# Maternal diet and gut microbiota influence predisposition to cardiovascular disease in the offspring

**DOI:** 10.1101/2022.03.12.480450

**Authors:** Hamdi Jama, Malathi S.I. Dona, Evany Dinakis, Michael Nakai, Madeleine R. Paterson, Waled Shihata, Crisdion Krstevski, Charles. D. Cohen, Kate L. Weeks, Gabriella E. Farrugia, Chad Johnson, Ekaterina Salimova, Daniel Donner, Helen Kiriazis, Harikrishnan Kaipananickal, Jun Okabe, Dovile Anderson, Darren J. Creek, Charles R. Mackay, Assam El-Osta, Alexander R. Pinto, David M. Kaye, Francine Z Marques

## Abstract

Cardiovascular disease is one of the most significant causes of death globally, especially in regions where unhealthy diets are prevalent and dietary fibre intake is low.^1,2^ Fibre, particularly prebiotic types that feed gut microbes, is essential for maintaining healthy gut microbial ecosystems.^3^ One assumption has been that cardiovascular health relates directly to lifestyle choices in adult life. Here, we show in mice that some of these benefits operate from the prenatal stage and relate to the diet and gut microbiome of the mother. Intake of fibre during pregnancy shaped the mothers’ gut microbiome, which had a lasting founding effect on the offspring’s microbial composition and function. Maternal fibre intake during pregnancy significantly changed the cardiac cellular and molecular landscape in the offspring, protecting them against the development of cardiac hypertrophy, remodelling, and inflammation. These suggest a role for foetal exposure to maternal-derived gut microbial metabolites, which are known to cross the placenta and drive epigenetic changes. Maternal fibre intake led to foetal epigenetic reprogramming of the atrial natriuretic peptide gene (*Nppa*), protective against heart failure. These results underscore the importance of dietary intake and the gut microbiome of the mother during pregnancy for cardiovascular disease in the offspring.

## Introduction

Diet is an important modifiable contributor to the development of cardiovascular disease (CVD).^2,4^ Dietary fibres are carbohydrates that remain undigested and unabsorbed until they reach the large intestine. Some of these fibres, such as resistant starches, are metabolised by gut microbial communities and considered prebiotic.^3,5^ Westernised diets, often very low in dietary fibre, are associated with increased risk of cardio-metabolic diseases.^6^ Conversely, the risk of CVD is decreased by diets high in fibre,^7^ such as the Mediterranean diet.^8,9^ Yet, the prevalence and incidence of CVD continue to rise, while fibre intake continues to decrease.^10,11^

Maternal nutrient intake and intrauterine exposures to nutrients and metabolites are fundamental determinants of postnatal outcomes.^11^ Nutrient quality and accessibility during foetal life underpin the developmental origins of health and disease.^11^ This hypothesis is supported by experimental, clinical and epidemiological studies, and result in the induction of cardio-metabolic and other non-communicable diseases.^12^ The concept of maternal nutritional constraint is a fundamental non-genetic factor regulating foetal development.^12–14^ Nutritional and metabolite allocation during foetal development significantly impacts growth and, more importantly, epigenetically regulates offspring’s postnatal phenotype and physiological variation.^13^ Growth restrictions during the perinatal period are associated with accelerated postnatal growth and subsequent development of coronary heart disease.^14^ However, the impact of maternal-derived gut metabolites and fibre intake as epigenetic mediators of the offspring’s cardiovascular outcome is yet to be determined.

Fibre intake has a significant impact on the composition and function of the gut microbiota,^15^ an emerging player in CVD.^16^ Increasing evidence supports an important role for the maternal gut microbiota in the offspring, both on the appropriate development, such as the priming of the immune system,^17^ and emergence of adverse phenotypes, such as metabolic syndrome.^18^ The prevailing theories suggest the relationship between offspring development and maternal gut microbiota is mediated by placental transfer of microbial-derived metabolites, such as short-chain fatty acids (SCFAs),^19^ which epigenetically influence the expression of genes.^19,20^ Therefore, *in utero* epigenetic changes, through regulation and modification of histone proteins, may be instrumental in determining the intergenerational risk of CVD. Thus, understanding the changes resulting from maternal fibre intake and cardiovascular health in the offspring can untangle the elusive ‘*why*’ in the continued rise in CVD prevalence globally.

Here we demonstrate that maternal fibre intake, through changes in the gut microbiota, associated metabolites and the chromatin landscape, modulates the cardiac phenotype of the adult offspring and their predisposition to CVD. We also discovered augmented and persistent metabolic changes in the offspring’s microbiome. Tackling CVD may require approaches that span all stages of life, including foetal development. Understanding microbiota and metabolite mechanisms has important implications for the prevention of CVD globally.

## Results

### Maternal fibre intake protects the offspring against CVD

Current experimental evidence supports the notion that high fibre intake elicits cardio-protective effects in the first generation via the gut microbiota.^21,22^ To determine whether this cardio-protection might persist in the second generation, pregnant C57BL/6J female mice were fed high-fibre or low-fibre isocaloric and nutrient-matched diets for the duration of pregnancy and breastfeeding (Supplementary Table 1). Although the high and low fibre diets were calorically matched, low-fibre fed dams had lower post-weaning weight, but we found no difference in these dams’ cardiac to body weight ratio (Supplementary Figure 1). After weaning, all offspring were placed on the same standard chow diet. The cardiac phenotypes of adult mice were compared after a hypertensive insult using angiotensin II (Ang II, 0.25mg/kg/day) starting at 6-weeks of age for four weeks (Figure 1A). Following Ang II administration, male offspring from low fibre-fed dams had significantly increased heart relative to both body weight and tibia ratio (Figure 1b and Supplementary Figure 2), while body weight remained similar (Supplementary Figure 2). The addition of fibre during pregnancy protected male offspring against an adverse hypertrophic response (Figure 1b and Supplementary Figure 2). We studied a small number of female offspring, and these did not have the same response to maternal fibre, which may be explained by an increase in their body weight with maternal fibre intake (Supplementary Figure 2). Next, we assessed heart function by echocardiography, and found male offspring from low-fibre mothers had increased left ventricular posterior wall thickness in diastole but not in systole (Figures 1c-d). Moreover, maternal low-fibre intake decreased the male offspring’s ejection fraction irrespective of Ang II treatment (Figure 1e). The cardiac phenotype was also assessed using cardiac catheterisation. Male offspring from low-fibre dams had an Ang II-induced increase in left ventricular systolic pressure and cardiac contractility (Supplementary Figure 3a-b). We observed no difference in heart rate and diastolic volume in echocardiography (Supplementary Figure 3c-d). Offspring from high-fibre dams had lower systolic and diastolic blood pressure; however, this difference in blood pressure was not maintained after Ang II-infusion (Supplementary Figure 4a-b), suggesting that the changes in the cardiac phenotype were independent of blood pressure.

**Figure 1.**
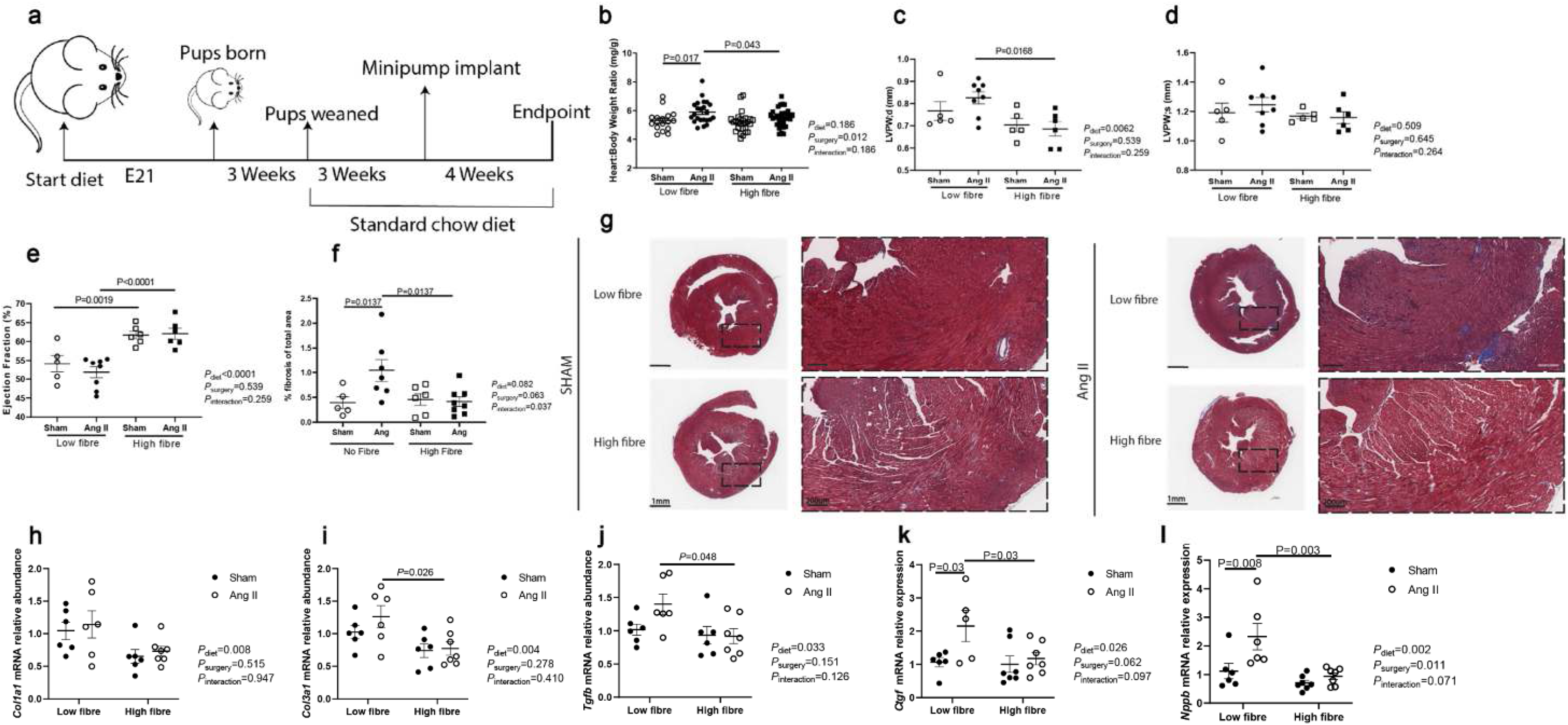
Maternal fibre intake protects against CVD. a) Schematic showing the experimental design. Following confirmation of a vaginal plugs, C57BL/6 dams were placed on a high-fibre or low-fibre diet for the duration of pregnancy and lactation. Once the offspring reached 3-weeks of age (standard weaning age), all offspring were placed on a standard chow diet for the duration of the experimental protocol. At 6-weeks of age, offspring were randomised to either sham (0.9% saline) or Ang II (0.25 mg/kg/d) treatment groups. b) Heart to body weight ratio of male offspring in mg/g, n=14-23/group; echocardiographic measurements showing c) left ventricular posterior wall (LVPW) thickness in diastole (mm), d) LVPW thickness in systole (mm) and e) ejection fraction as a percentage. f) Quantification of Masson’s trichome staining (collagen) as a percentage of total ventricular area and g) representative micrographs with the magnified comparison of high-fibre and low-fibre Ang II treated heart tissue. Scale bars are 1mm and 200μm respectively. Relative gene expression of h) Collagen 1 a1 (*Col1a1*), i) Collagen 3 a1 (*Col3a1*), j) transforming growth factor-beta (*Tgfb*), k) connective tissue growth factor (*Ctgf*), and l) natriuretic peptide b (*Nppb*) mRNA. All data is shown as mean±SEM. Panels C to J: n=5-8/treatment group, 2-way ANOVA with false discovery rate adjustment for multiple comparisons. P-values are adjusted for false discovery rate.

To further characterise the cardiac phenotype in offspring from low-fibre dams, we assessed the expression of markers associated with cardiac pathology. Maternal high-fibre significantly decreased collagen deposition in the heart (Figures 1f-g), and the expression of profibrotic collagen 1 *Col1a1*, collagen 3 *(Col3a1)*, transforming growth factor-beta (*Tgfb*), and connective tissue growth factor (*Ctgf*) mRNA (Figures 1h-k). Low-fibre in the maternal diet combined with the Ang-II insult increased the expression of natriuretic peptide B (*Nppb*); the addition of fibre in the maternal diet significantly blunted this response (Figure 1l). Together, these results support the notion that maternal fibre intake protects the adult offspring against an adverse cardiac phenotype.

### Maternal fibre intake leads to changes in the offspring’s gut microbiota

Alterations in the gut microbiota have emerged as a common feature and contributor to the development of CVD.^23^ Clinical and preclinical studies have identified distinct microbial signatures with fibre intake^24^ that are evident in CVD, such as heart failure.^25^ The gut microbial ecology is readily adaptable by dietary manipulations such as fibre.^26^ To determine the influence of maternal fibre intake on the offspring’s gut microbial composition, we first sequenced the bacterial 16S rRNA from caecal content collected from female C57BL/6J dams and their offspring. As expected, dams fed diets low or high in fibre had a different gut microbiota composition (Figure 2a and Supplementary Figure 5). Male offspring born to dams who received a high-fibre diet had distinct gut microbial colonisation that persisted into adulthood (Figure 2b), irrespective of sham or Ang II treatment. These findings were validated in independent cohorts (Supplementary Figure 5). This suggests the existence of a founder effect on the gut microbiota of the offspring in response to maternal fibre intake. We observed no differences in α-diversity measured by Shannon index and Chao1 between high-fibre and low-fibre samples (Supplementary Figure 6). Using linear discriminant analysis, we identified low-fibre offspring had higher abundance of *Akkermansia sp*., *Azosporillium sp*. and *Clostridiales bacterium*, while high-fibre offspring had more *Barnesiella sp*. (Figures 2c-d and Supplementary Figure 6).

**Figure 2.**
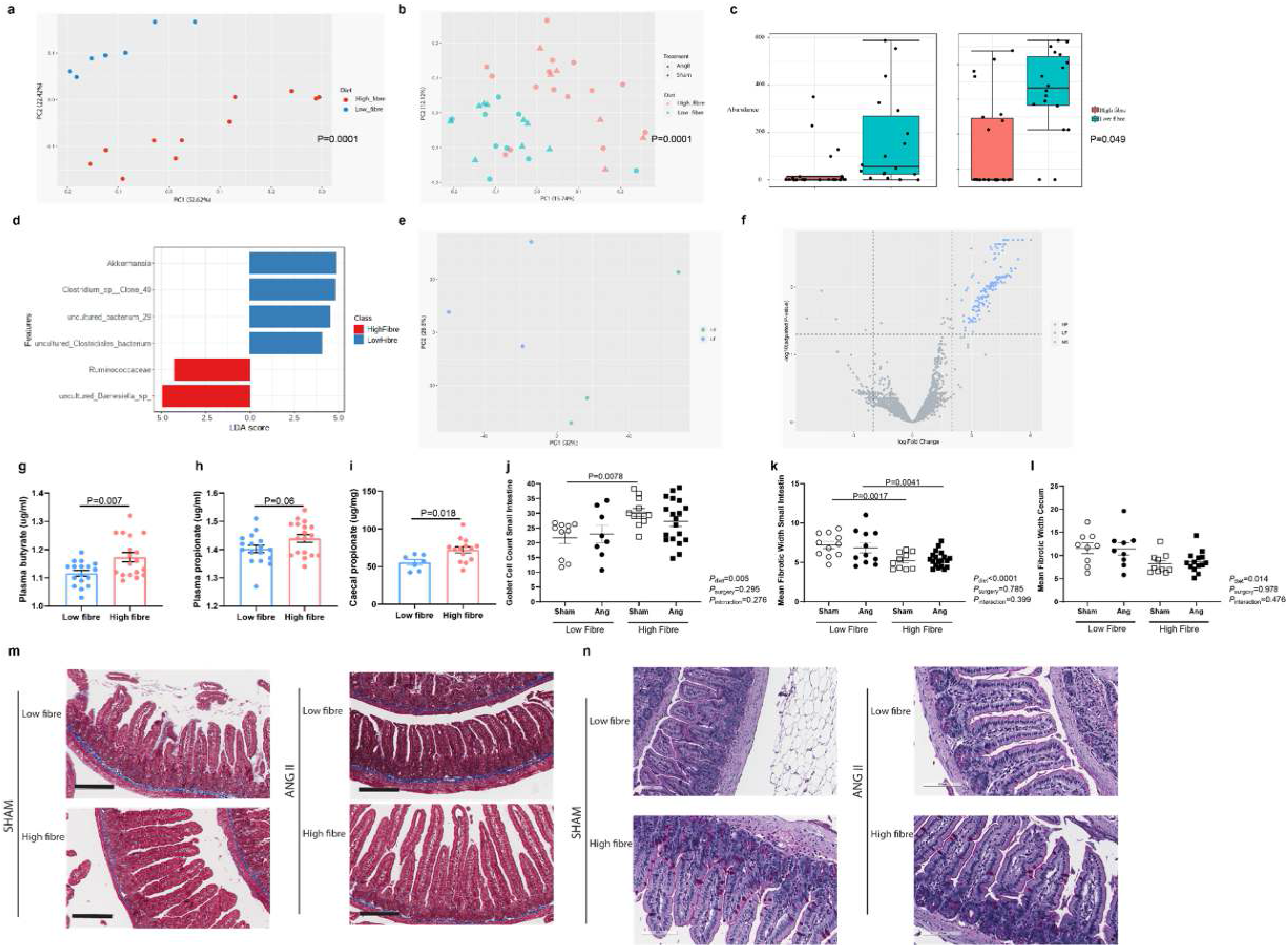
Maternal fibre intake alters microbial composition in dams and their offspring. a) Principal coordinate analysis showing β-diversity (weighted unifrac) in high-fibre (red) and low (blue)-fibre fed dams, n=7-11/diet, b) Principal coordinate analysis (PCoA) showing β-diversity (weighted unifrac) offspring from high-fibre(red) and low-fibre(blue) fed dams, n=6-15/group. c) Relative abundance of *Akkermansia* in high-fibre and low-fibre offspring, adjusted for multiple comparisons. d) Linear discriminant analysis effect size showing abundant features in the caecum of high-fibre and low-fibre offspring. Metagenomic analysis showing e) PCoA showing gene family abundance grouped by enzyme commission number in offspring from high-fibre and low-fibre dams, f) volcano plot showing differentially abundant gene families in high- and low-fibre offspring (n=3/diet). Short-chain fatty acid (SCFA) quantification in plasma and caecal samples from offspring: g) plasma butyrate, h) plasma propionate, and i) caecal propionate. Quantification of histological staining showing j) Goblet cell count in small intestines, k) mean fibrotic width in small intestine, and l) mean fibrotic width in the cecum. m) representative micrographs of high-fibre and low-fibre Ang II treated small intestine tissue. Scale bar 200μm. All data is shown as ±SEM. 2-way ANOVA with FDR adjustment for multiple comparisons. P-values are adjusted for false discovery rate.

We utilised shotgun metagenome sequencing of caecal content from the adult offspring to further validate these findings and to identify specific differential species and microbial genes. We identified 1,128 statistically significant differentially expressed gene families in the caecal content from low- and high-fibre offspring, confirming the differences in microbial colonisation (Figure 2e). Of these differentially expressed gene families, only 36 were in high-fibre samples (Supplementary Table 2). We then interrogated enzymatic pathways and found 174 enzyme signatures enriched in low-fibre offspring (Figure 2f, Supplementary Table 3). One hundred fifty-four of the identified enzyme signatures in low-fibre belonged to *Akkermansia muciniphila*, and one belonged to *Parabacteroides goldsteinii*. *Akkermansia muciniphila* upregulated genes encoding for mucolytic enzymes, including sulfatases, glycosylhydrolases, glycosyltransferases, β-N-acetylhexosaminidases, alpha-galactosidase and beta-galactosidase (Supplementary Table 3). Pathway analysis showed 10 microbial pathways enriched in low-fibre offspring, of which 3 were attributed to *Akkermansia muciniphila* (Supplementary Table 4). In contrast, high-fibre offspring had only 5 grouped enzyme signatures which belonged to *Bacteroides ovatus, Escherichia coli* and *Lactobacillus murinus;* the latter was previously shown to reduce inflammatory pathways and blood pressure associated with sodium intake.^27^

Dietary intake of fibre leads to the production of SCFAs by the gut microbiota. In preclinical models, SCFAs are cardioprotective.^21,22^ Indeed, we identified mice fed a high fibre diet during pregnancy had 50% higher plasma total SCFA and 71% higher plasma acetate levels (Supplementary Figure 7). We hypothesised that high-fibre offspring would have acquired at birth from their mothers a gut microbiota that may be more efficient in producing SCFAs, independent of their post-weaning diet. We quantified plasma and caecal levels of acetate, butyrate, propionate and total SCFAs, and found higher levels of butyrate and propionate between high-fibre and low-fibre male offspring (Figure 2g-i).

These gut microbial and metabolite composition changes can be accompanied by increased gut inflammation and subsequent gastrointestinal architecture and morphology changes. Thus, we assessed histological sections of the small and large intestines in the offspring. We observed no difference in villi length in the small intestine of offspring from low-fibre and high-fibre linage (Supplementary Figure 8). However, offspring from low-fibre fed dams had reduced goblet cell count and increased collagen deposition in the small and large intestines (Figures 2j-n). Similarly to gut microbial colonisation, chronic Ang II infusion did not impact these findings. Together, these data suggest maternal fibre intake during pregnancy induces marked microbial and intestinal changes that persist in the adult offspring.

### Single-cell RNA-sequencing reveals maternal fibre intake induces differential cardiac cellulome

We hypothesised that maternal fibre intake shaped, *in utero*, the epigenetic environment and modified the cardiac phenotype of offspring. We also hypothesised that maternal fibre intake, through SCFA production, might also induce marked cardiac transcriptional changes in the offspring. Thus, to understand how maternal high fibre intake shaped the cardiac cellulome, we performed single-cell RNA-sequencing (scRNA-seq) in 6-week-old naïve, unchallenged offspring (Supplementary Figure 9). A total of 25,557 cells were deep sequenced (94,006– 252,655/reads per cell). We identified and annotated 16 distinct clusters of cells using cluster differentiation markers, including mesenchymal cells, immune cells, and stromal cells (Supplementary Figure 9). Differential gene expression analysis revealed 2,686 up- and 2,379 down-regulated genes between high-fibre and low-fibre offspring across all cell types (Figure 3A, Supplementary Table 5). Gene ontology analyses revealed a markedly different transcriptional profile with several genes related to protein folding, and stabilisation upregulated in a number of cell types from high-fibre offspring (Figure 3c, Supplementary Table 6). Conversely, maternal high fibre intake down-regulated regulation of inflammatory and immune responses in macrophages and B cells, particularly related to interferon-β, and extracellular structure organisation, TGFβ, connective tissue development and collagen metabolic processes (Figure 3b, Supplementary Figure 10a-d), consistent with our findings regarding lower collagen deposition (Figure 1g-I).

**Figure 3.**
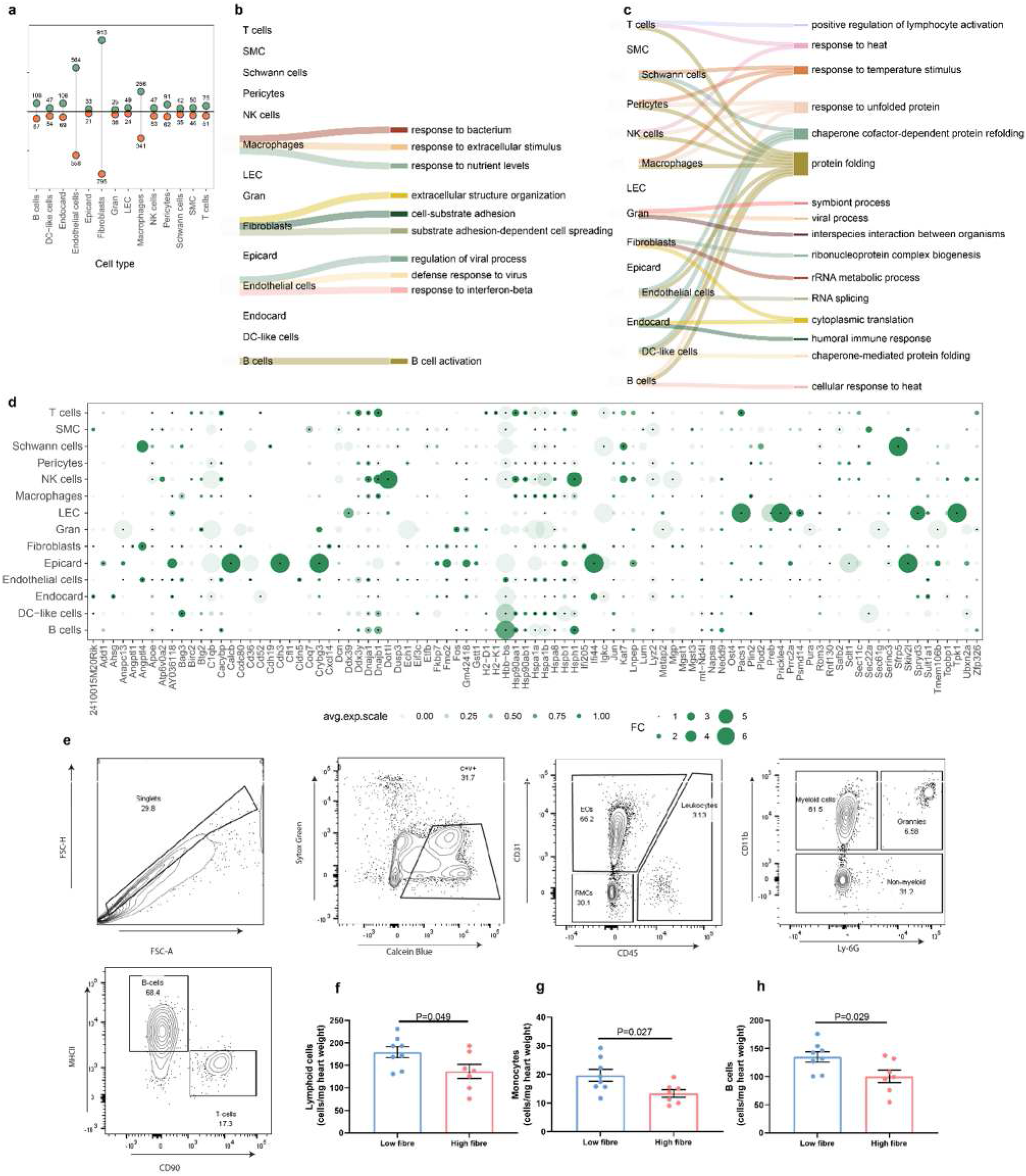
Maternal fibre intake influences the offspring’s cardiac transcriptome and cellulome. a) Lollipop plot showing the number of up- (green) and down- (red) regulated genes between high-fibre and low-fibre maternal diet (see Supplementary Table 6). Sankey plot summarising top 3 b) down-regulated and c) up-regulated Gene Ontology biological processes in high-fibre offspring. d) Dot plot showing top 10 up-regulated genes in each cell type in high-fibre offspring (n=8/diet, 4 male and 4 female). For data analysis please see methods. e) Flow cytometry gating strategy. F) Flow cytometric assessment of heart tissue of male offspring f) lymphoid cells, g) monocytes, and F) B cells normalised to heart weight and expressed as cells/mg heart weight (n=7-8/group). All data is shown as ±SEM. 2-way ANOVA with FDR adjustment for multiple comparisons. P-values are adjusted for false discovery rate.

We then utilised a flow cytometry panel in independent animals to determine the impact of maternal fibre intake on the cardiac cellular composition (Figure 3e). Maternal high-fibre intake reduced the abundance of cardiac lymphoid cells, particularly B cells, independent of sex (Figure 3f-h). Thus, these findings suggest maternal fibre intake prevents aberrant cardiac inflammation and fibrosis at both cellular and molecular levels in the offspring.

### Maternal fibre intake induces epigenetic regulation of atrial natriuretic peptide

A previous study in a model of allergic airway disease showed the protective effect of maternal fibre intake was mediated through the inhibition of HDAC9.^19^ This histone deacetylase is primarily involved.16 We hypothesised that lack of HDAC9 would result in a lack of response to a high-fibre diet, which we tested in HDAC9^−/−^ offspring. We found HDAC9^−/−^ offspring from high-fibre mothers were still protected from Ang II-induced cardiac hypertrophy independent of an increase in blood pressure (Supplementary Figure 11). To determine if these changes were reflected in gene expression, we quantified *Col1a1* and *Nppb* mRNA, and found no difference in their expression (Supplementary Figure 11). This may suggest high fibre-driven cardio-protection is not mediated through inhibition of HDAC9. Along with HDAC9, HDAC5 regulates cardiac hypertrophic response to stress.^28^ Indeed, *Hdac5* gene expression was affected by fibre intake, with a high-fibre diet significantly increasing its expression in wild-type male offspring independently of Ang II treatment (Supplementary Figure 11). This data suggests that in our model, HDAC5, and not HDAC9, may be a critical mediator of cardioprotection by maternal dietary fibre.

Besides HDAC inhibition, SCFAs were recently discovered to impact acetylation.^29^ Thus, we assessed the level of acetylated histones in cardiac tissue using western blot. We found an increase in global histone3 acetylation at lysines 9 and 14 (H3K9/14ac) in low-fibre compared to high-fibre Ang II-treated offspring (Figure 4a). We observed high-fibre offspring had decreased expression of the atrial natriuretic peptide A gene (*Nppa*) compared to low-fibre offspring (Figure 4b). We performed a chromatin-immunoprecipitation (ChIP) assay to elucidate further the epigenetic changes in this gene (Figure 4c). The ChIP assay revealed a decrease in H3K9/14ac14ac at the promoter region of the *Nppa* gene in Ang II-treated high-fibre offspring (Figure 4d), suggesting that maternal fibre intake influences the epigenetic changes of the *Nppa* gene in the offspring’s heart. Together, these data reveal maternal high-fibre intake leads to foetal epigenetic reprogramming, likely through maternal to foetal transfer of gut microbial-derived metabolites that impact at least one crucial gene for cardiac function, *Nppa*.

**Figure 4.**
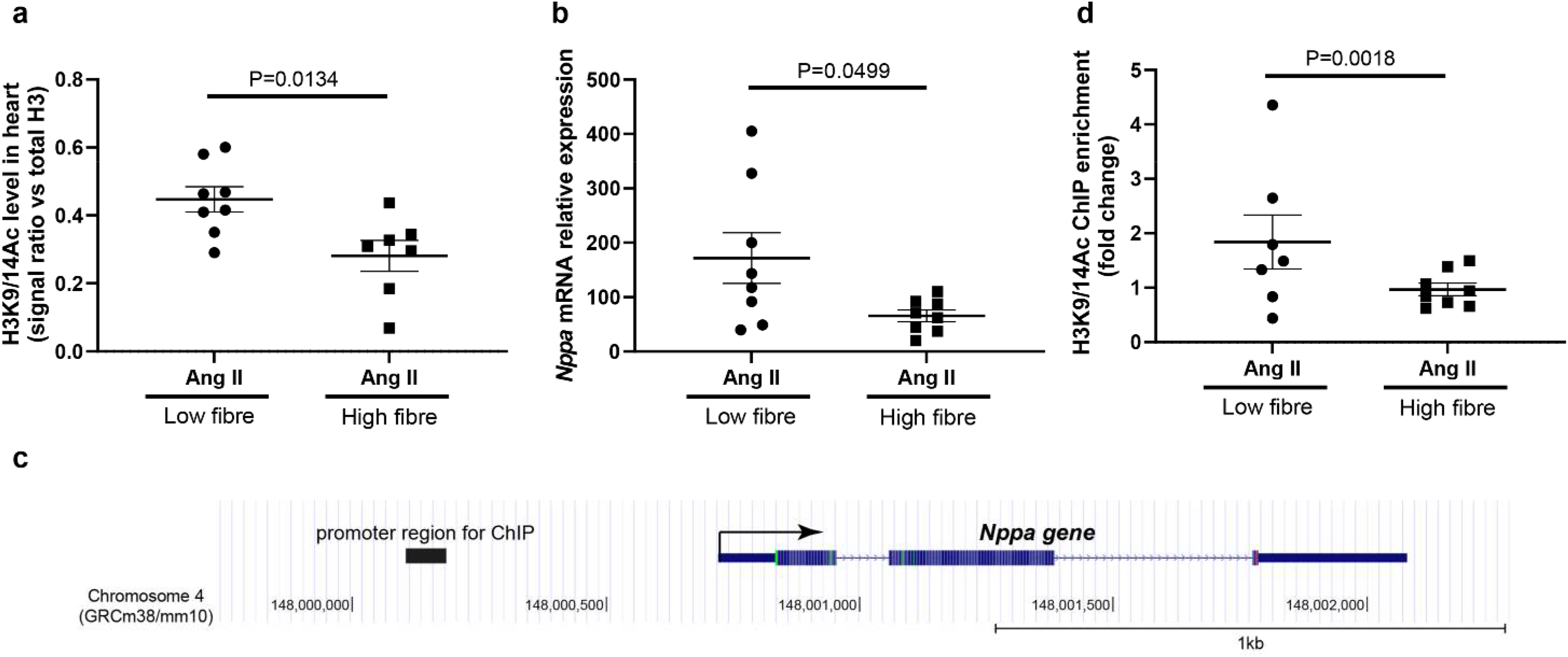
Maternal fibre intake promotes cardioprotective-acetylation acetylation changes. a) Quantification of histone H3 lysine 9 and 14acetylation (H3K9acH3K9/14ac14acH3K9ac) in cardiac tissue from male offspring. b) Relative mRNA gene expression of *Nppa*. c) Relative enrichment of H3K9acH3K9/14ac at the promoter H3K9ac region of *Nppa* gene in cardiac tissue samples from high-fibre and low-fibre Ang II-treated male offspring. d) Representation of natriuretic peptide A (*Nppa*) gene promoter region. The solid box the region targeted for ChIP assay. All data is shown as ±SEM. n=7-8/group, Mann-Whitney test.

## Discussion

Our study provides an important conceptual advancement in the developmental origins of CVD. We found that maternal fibre intake during pregnancy protected offspring against the development of an adverse cardiac phenotype associated with hypertension (using chronic Ang II-infusion), including cardiac hypertrophy, haemodynamic function and collagen deposition. Moreover, maternal fibre intake induced marked changes in the offspring’s gut microbial ecology and intestine irrespective of Ang II treatment, suggesting an intergenerational founding effect of the gut microbiota. These changes were also accompanied by a distinct cardiac cellular and transcriptomic phenotype in the offspring, even in the absence of Ang II, and epigenetic reprogramming of the atrial natriuretic peptide *Nppa*.

A longitudinal study of 25 human mother-infant pairs showed substantial early seeding of the infant gut microbial communities from strains present in the maternal gut.^25^ Our study supports that the acquisition and development of the infant microbiome is seeded early in life, is dependent on the maternal diet during pregnancy, and persists into later life. Our findings advance this concept by demonstrating maternal fibre intake profoundly impacted offspring gut microbial taxonomy and function, supporting the notion that early life dietary exposure shapes later life microbial ecology. Thus, a shift in microbial composition, facilitated through diet, is vertically transmitted to offspring and leads to long-lasting colonisation that may drive health and disease states.

Dietary constituents, including protein and fat, alter and shape the offspring’s microbiome.^30–33^ Our findings add prenatal and neonatal exposure to fibre as an essential dietary component that shapes the gut microbiome. This finding is impactful, as there is a rise in diets low in dietary fibre.^2^ Increasing evidence supports that the maternal microbial environment is an important determinant of autoimmune disorders, neurodevelopmental disorders and metabolic syndrome.^18,19,34^ We demonstrated that low-fibre offspring had a distinct microbiome with altered metabolic function and gastrointestinal homeostasis, including reduced number of mucus-producing goblet cells. Low-fibre offspring had more abundant *Akkermansia muciniphila* and upregulated several genes encoding for enzymes necessary for the degradation of host mucins. This data is also consistent with previous studies demonstrating that *Akkermansia muciniphila* is not strictly fibrolytic and can utilise host glycans and mucins without dietary fibre.^35^ This switch in the metabolic program has been observed in humanised mice with selective bacteria and those directly fed a low-fibre diet.^35,36^ However, in our model, *Akkermansia muciniphila* seeded from low-fibre mothers, presumably at birth, were metabolically imprinted with genes necessary to survive in a low-fibre environment. Even in the presence of a normal chow-diet (and, thus, normal fibre levels), these microbes persisted with their low-fibre metabolic program and likely contributed to the observed reduction in mucus-producing goblet cells. It is unclear why microbiota seeded from low-fibre mothers persist with this augmented metabolic program as previous studies have shown that the gut microbiome is rapidly modulated by diet.^37,38^ However, this sheds light on a potential mechanism in which maternal low-fibre influences offspring susceptibility to later life disease by augmenting metabolic changes in acquired microbes. Nevertheless, excessive mucus degradation by commensal microbes weakens the gut barrier and exacerbates intestinal inflammation.^39^ In severe gut inflammation, the presence of *Akkermansia muciniphila* resulted in worsened inflammation and disturbed mucus homeostasis.^40^ Expansion of *Akkermansia muciniphila* also impacts the microbial ecosystem, particularly in low fibre settings, substantially reducing fibrolytic and SCFA-producing bacteria.^36^ Together, our data support lack of fibre in the maternal diet promotes the seeding and expansion of a metabolically altered *Akkermansia muciniphila*, fostering a gut microbiota ecosystem that supports an inflammatory environment, degradation of intestinal mucin and increased intestinal fibrosis.

Interrogation of the single-cell transcriptome revealed remarkable differences in the gene expression of low- and high fibre offspring. These fundamental differences in the naïve heart undoubtedly influence cardiovascular outcomes. In the absence of fibre, offspring had a baseline transcriptomic profile that, under secondary stress such as Ang II, may exacerbate cardiac phenotype and hypertrophy, notably increasing collagen deposition in cardiac fibroblasts. Indeed, this was observed when we challenged offspring with an Ang II insult, supporting the worse hemodynamic function observed. Together, these data strongly implicate maternal dietary fibre during pregnancy as a driver of cardiac outcome in the offspring. Low-fibre intake is implicated in the development of obesity, lupus and cardiovascular disease in the first generation.^22,41^ Our current findings importantly demonstrate the development of these may indeed originate from lack of prenatal fibre intake and the constraint of maternal-derived metabolites, such as SCFA, to the developing foetus. Maternal high-fibre functionally shaped the cardiac cellulome and promoted a cardio-protective phenotype in offspring. Thus, pre-and postnatal exposure to fibre and maternal gut-derived metabolites may be novel therapeutic interventions to improve cardiac health and prevent CVD.

We showed maternal fibre intake during pregnancy influenced offspring response to cardiovascular stress. Previous studies demonstrate the protective effects of fibre are mediated predominately through microbial-derived metabolites such as SCFAs.^18,22^ The precise mechanisms by which maternal gut microbiota-derived metabolites influence foetal development are yet to be elucidated. However, the current dogma suggests these maternal gut microbial-derived metabolites traverse the placental circulation and trigger several signalling cascades through G-protein coupled receptors and inhibition of histone deacetylases, thus altering gene expression.^19^ Our data demonstrate microbiota seeded from mothers receiving a high fibre diet results in microbial communities adapted to producing higher levels of propionate and butyrate in the offspring. Thus, disturbances in fibre intake during pregnancy may perturb offspring’s SCFA-producing capability in later life. Given the increasingly poor global consumption of dietary fibre and maternal constraint of beneficial fibre-derived metabolites, the physiological consequences may be far-reaching.

SCFA are described as being inhibitors of histone deacetylases.^42,43^ Recent studies show the therapeutic benefit of pharmacological HDAC inhibition regulating the acetylation index in preclinical cardiomyopathy.^44^ Given previously published data implicated HDAC9, we investigated whether the cardioprotection following maternal fibre intake was mediated through inhibition of HDAC9 and subsequent transcriptional regulation of genes. HDAC9 knockout offspring were protected from Ang II-induced cardiac hypertrophy. In addition, no differences in HDAC9 protein levels were observed in the cardiac tissue of offspring, suggesting the cardiac phenotype imprinted by maternal fibre is through a HDAC9-independent mechanism. To this end, we investigated the chromatin acetylation profile of offspring cardiac tissue, focusing on a gene highly relevant to cardiac function: *Nppa*. Maternal fibre intake during pregnancy leads to a decrease in histone H3 acetylation at both genome-wide and the cis-regulatory region of *Nppa*. These changes were accompanied by a reduction in the expression of the *Nppa* gene in the presence of maternal fibre, suggesting maternal fibre intake influences the epigenetic changes of the *Nppa* gene in the offspring’s heart. However, the precise maternal-derived epigenetic mediator responsible for these changes is yet to be elucidated.

In conclusion, gut microbes, particularly *Akkermansia muciniphila* and maternal-derived epigenetic mediators, lead to cardiac and gastrointestinal phenotypic changes in the offspring in response to maternal fibre intake. Maternal low-fibre diet had a profound effect on gut microbiota’s metabolic function, which persisted in offspring into adulthood. Maternal fibre intake significantly altered the molecular, cellular and functional cardiac outcome of offspring. Thus, during pregnancy, maternal fibre intake, particularly prebiotic fibre, promotes a protective cardiac cellular and molecular environment advantageous for the offspring’s future cardiovascular health. With the continued growing global burden of cardiovascular disease, maternal fibre intake may represent an inexpensive and readily available preventative opportunity.

## Methods

Please see the Online Supplementary file.

## Acknowledgements

We would like to acknowledge the Monash Animal Research Platform (MARP) for their help with animal housing, the Monash Histology Platform and Micro Imaging Facility for their help with histological staining, microscopy and analysis, and the Monash Proteomics and Metabolomics Facility for SCFA analysis. We would like to also acknowledge Monash Bioinformatics Platform for their assistance in analysing microbiome data and access to M3 servers.

## References

1 James, S. L. et al. Global, regional, and national incidence, prevalence, and years lived with disability for 354 diseases and injuries for 195 countries and territories, 1990–2017: a systematic analysis for the Global Burden of Disease Study 2017. The Lancet 392, 1789–1858, doi:10.1016/S0140-6736(18)32279-7 (2018).

2 Reynolds, A. et al. Carbohydrate quality and human health: a series of systematic reviews and meta-analyses. The Lancet 393, 434–445, doi:10.1016/S0140-6736(18)31809-9 (2019).

3 Slavin, J. Fiber and prebiotics: mechanisms and health benefits. Nutrients 5, 1417–1435, doi:10.3390/nu5041417 (2013).

4 Yu, E., Malik, V. S. & Hu, F. B. Cardiovascular Disease Prevention by Diet Modification: JACC Health Promotion Series. Journal of the American College of Cardiology 72, 914–926, doi:https://doi.org/10.1016/j.jacc.2018.02.085 (2018).

5 Gill, S. K., Rossi, M., Bajka, B. & Whelan, K. Dietary fibre in gastrointestinal health and disease. Nat Rev Gastroenterol Hepatol 18, 101–116, doi:10.1038/s41575-020-00375-4 (2021).

6 Benjamin Emelia, J. et al. Heart Disease and Stroke Statistics—2019 Update: A Report From the American Heart Association. Circulation 139, e56–e528, doi:10.1161/CIR.0000000000000659 (2019).

7 Reynolds, A. et al. Carbohydrate quality and human health: a series of systematic reviews and meta-analyses. Lancet 393, 434–445, doi:10.1016/S0140-6736(18)31809-9 (2019).

8 de Lorgeril, M. et al. Mediterranean diet, traditional risk factors, and the rate of cardiovascular complications after myocardial infarction: final report of the Lyon Diet Heart Study. Circulation 99, 779–785 (1999).

9 Ndanuko, R. N., Tapsell, L. C., Charlton, K. E., Neale, E. P. & Batterham, M. J. Dietary Patterns and Blood Pressure in Adults: A Systematic Review and Meta-Analysis of Randomized Controlled Trials. Adv Nutr 7, 76–89, doi:10.3945/an.115.009753 (2016).

10 O’Keefe, S. J. The association between dietary fibre deficiency and high-income lifestyle-associated diseases: Burkitt’s hypothesis revisited. Lancet Gastroenterol Hepatol 4, 984–996, doi:10.1016/S2468-1253(19)30257-2 (2019).

11 Popkin, B. M. Global nutrition dynamics: the world is shifting rapidly toward a diet linked with noncommunicable diseases. Am J Clin Nutr 84, 289–298, doi:10.1093/ajcn/84.1.289 (2006).

12 Gluckman, P. D., Hanson, M. A., Cooper, C. & Thornburg, K. L. Effect of in utero and early-life conditions on adult health and disease. N Engl J Med 359, 61–73, doi:10.1056/NEJMra0708473 (2008).

13 Charalambous, M., da Rocha, S. T. & Ferguson-Smith, A. C. Genomic imprinting, growth control and the allocation of nutritional resources: consequences for postnatal life. Curr Opin Endocrinol Diabetes Obes 14, 3–12, doi:10.1097/MED.0b013e328013daa2 (2007).

14 Barker, D. J., Osmond, C., Forsén, T. J., Kajantie, E. & Eriksson, J. G. Trajectories of growth among children who have coronary events as adults. N Engl J Med 353, 1802–1809, doi:10.1056/NEJMoa044160 (2005).

15 Xu, Z. & Knight, R. Dietary effects on human gut microbiome diversity. Br J Nutr 113 Suppl, S1–5, doi:10.1017/S0007114514004127 (2015).

16 Marques, F. Z., Mackay, C. R. & Kaye, D. M. Beyond gut feelings: how the gut microbiota regulates blood pressure. Nature Reviews Cardiology 15, 20–32, doi:10.1038/nrcardio.2017.120 (2018).

17 Nyangahu, D. D. et al. Disruption of maternal gut microbiota during gestation alters offspring microbiota and immunity. Microbiome 6, 124–124, doi:10.1186/s40168-018-0511-7 (2018).

18 Kimura, I. et al. Maternal gut microbiota in pregnancy influences offspring metabolic phenotype in mice. Science 367, doi:10.1126/science.aaw8429 (2020).

19 Thorburn, A. N. et al. Evidence that asthma is a developmental origin disease influenced by maternal diet and bacterial metabolites. Nat Commun 6, 7320, doi:10.1038/ncomms8320 (2015).

20 Roseboom, T. J. Epidemiological evidence for the developmental origins of health and disease: effects of prenatal undernutrition in humans. The Journal of endocrinology 242, T135–t144, doi:10.1530/joe-18-0683 (2019).

21 Marques, F. Z. et al. High-Fiber Diet and Acetate Supplementation Change the Gut Microbiota and Prevent the Development of Hypertension and Heart Failure in Hypertensive Mice. Circulation 135, 964–977, doi:10.1161/CIRCULATIONAHA.116.024545 (2017).

22 Kaye, D. M. et al. Deficiency of Prebiotic Fiber and Insufficient Signaling Through Gut Metabolite-Sensing Receptors Leads to Cardiovascular Disease. Circulation 141, 1393–1403, doi:10.1161/circulationaha.119.043081 (2020).

23 Tang, W. H. W., Kitai, T. & Hazen, S. L. Gut Microbiota in Cardiovascular Health and Disease. Circ Res 120, 1183–1196, doi:10.1161/CIRCRESAHA.117.309715 (2017).

24 Jama, H., Beale, A., Shihata, W. A. & Marques, F. Z. The effect of diet on hypertensive pathology: is there a link via gut microbiota-driven immune-metabolism? Cardiovasc Res 115, 1435–1447, doi:10.1093/cvr/cvz091 (2019).

25 Beale, A. L. et al. The Gut Microbiome of Heart Failure With Preserved Ejection Fraction. J Am Heart Assoc 10, e020654, doi:10.1161/JAHA.120.020654 (2021).

26 David, L. A. et al. Diet rapidly and reproducibly alters the human gut microbiome. Nature 505, 559–563, doi:10.1038/nature12820 (2014).

27 Wilck, N. et al. Salt-responsive gut commensal modulates TH17 axis and disease. Nature 551, 585–589, doi:10.1038/nature24628 (2017).

28 Chang, S. et al. Histone deacetylases 5 and 9 govern responsiveness of the heart to a subset of stress signals and play redundant roles in heart development. Mol Cell Biol 24, 8467–8476, doi:10.1128/MCB.24.19.8467-8476.2004 (2004).

29 Thomas, S. P. & Denu, J. M. Short-chain fatty acids activate acetyltransferase p300. Elife 10, e72171, doi:10.7554/eLife.72171 (2021).

30 Ma, J. et al. High-fat maternal diet during pregnancy persistently alters the offspring microbiome in a primate model. Nat Commun 5, 3889, doi:10.1038/ncomms4889 (2014).

31 Schutkowski, A. et al. Impact of a high-protein diet during lactation on milk composition and offspring in a pig model. Eur J Nutr 58, 3241–3253, doi:10.1007/s00394-018-1867-y (2019).

32 Hsu, C. N., Hou, C. Y., Lee, C. T., Chan, J. Y. H. & Tain, Y. L. The Interplay between Maternal and Post-Weaning High-Fat Diet and Gut Microbiota in the Developmental Programming of Hypertension. Nutrients 11, doi:10.3390/nu11091982 (2019).

33 Warren, M. F., Hallowell, H. A., Higgins, K. V., Liles, M. R. & Hood, W. R. Maternal Dietary Protein Intake Influences Milk and Offspring Gut Microbial Diversity in a Rat (Rattus norvegicus) Model. Nutrients 11, doi:10.3390/nu11092257 (2019).

34 Kim, E. et al. Maternal gut bacteria drive intestinal inflammation in offspring with neurodevelopmental disorders by altering the chromatin landscape of CD4+ T cells. Immunity, doi:https://doi.org/10.1016/j.immuni.2021.11.005 (2021).

35 Kovatcheva-Datchary, P. et al. Simplified Intestinal Microbiota to Study Microbe-Diet-Host Interactions in a Mouse Model. Cell Reports 26, 3772–3783.e3776, doi:https://doi.org/10.1016/j.celrep.2019.02.090 (2019).

36 Desai, M. S. et al. A Dietary Fiber-Deprived Gut Microbiota Degrades the Colonic Mucus Barrier and Enhances Pathogen Susceptibility. Cell 167, 1339–1353.e1321, doi:10.1016/j.cell.2016.10.043 (2016).

37 Sonnenburg, J. L. & Bäckhed, F. Diet-microbiota interactions as moderators of human metabolism. Nature 535, 56–64, doi:10.1038/nature18846 (2016).

38 Leeming, E. R., Johnson, A. J., Spector, T. D. & Le Roy, C. I. Effect of Diet on the Gut Microbiota: Rethinking Intervention Duration. Nutrients 11, 2862, doi:10.3390/nu11122862 (2019).

39 Paone, P. & Cani, P. D. Mucus barrier, mucins and gut microbiota: the expected slimy partners? Gut 69, 2232, doi:10.1136/gutjnl-2020-322260 (2020).

40 Ganesh, B. P., Klopfleisch, R., Loh, G. & Blaut, M. Commensal Akkermansia muciniphila Exacerbates Gut Inflammation in Salmonella Typhimurium-Infected Gnotobiotic Mice. PLOS ONE 8, e74963, doi:10.1371/journal.pone.0074963 (2013).

41 Schäfer, A.-L. et al. Low Dietary Fiber Intake Links Development of Obesity and Lupus Pathogenesis. Frontiers in Immunology 12, doi:10.3389/fimmu.2021.696810 (2021).

42 Kasubuchi, M., Hasegawa, S., Hiramatsu, T., Ichimura, A. & Kimura, I. Dietary gut microbial metabolites, short-chain fatty acids, and host metabolic regulation. Nutrients 7, 2839–2849, doi:10.3390/nu7042839 (2015).

43 Koh, A., De Vadder, F., Kovatcheva-Datchary, P. & Bäckhed, F. From Dietary Fiber to Host Physiology: Short-Chain Fatty Acids as Key Bacterial Metabolites. Cell 165, 1332–1345, doi:10.1016/j.cell.2016.05.041 (2016).

44 Khurana, I. et al. SAHA attenuates Takotsubo-like myocardial injury by targeting an epigenetic Ac/Dc axis. Signal Transduction and Targeted Therapy 6, 159, doi:10.1038/s41392-021-00546-y (2021).

