## Supplementary material for "Maternal diet and gut microbiota influence predisposition to cardiovascular disease in the offspring": Online supplemental tables and figures

^1^Hypertension Research Laboratory, School of Biological Sciences, Monash University, Melbourne, VIC., Australia; ^2^Heart Failure Research Group Baker Heart and Diabetes Institute, Melbourne, VIC., Australia; ^3^Cardiac Cellular Systems, Baker Heart and Diabetes Institute, Melbourne, VIC, Australia; ^4^Centre for Cardiovascular Biology and Disease Research, La Trobe University, Melbourne, Victoria, Australia; ^5^Department of Anatomy & Physiology, The University of Melbourne, Parkville, VIC, Australia; ^6^Baker Department of Cardiometabolic Health, The University of Melbourne, Parkville, VIC, Australia; ^7^Monash Micro Imaging, Monash University, Melbourne, Australia; ^8^La Trobe Institute of Molecular Science, La Trobe University, Melbourne, Victoria, Australia; ^9^Burnet Institute, Melbourne, VIC, Australia; ^10^Monash Biomedical Imaging, Monash University, Melbourne, Australia; ^11^Epigenetics in Human Health and Disease, Central Clinical School, Alfred Centre, Monash University Melbourne, VIC, Australia; ^12^Department of Clinical Pathology, University of Melbourne, VIC, Australia; ^13^Department of Diabetes, Central Clinical School, Monash University, Melbourne, VIC, Australia; ^14^Monash Institute of Pharmaceutical Sciences, Monash University, Melbourne, Australia; ^15^Infection and Immunity Program, Monash Biomedicine Discovery Institute, Monash University, Melbourne, Australia; ^16^Department of Biochemistry and Molecular Biology, Monash University, Melbourne, Australia; ^17^Department of Cardiology, Alfred Hospital, Melbourne, Australia; ^18^Central Clinical School, Faculty of Medicine Nursing and Health Sciences, Monash University, Melbourne, Australia;

***Corresponding author**: A/Prof Francine Z. Marques, PhD, Hypertension Research Laboratory, School of Biological Sciences, Faculty of Science, Monash University, Melbourne, Australia, Phone: +61-03-9905 6958.

**Methods**

**Animal experiments**

Male and female C57BL/6J mice were obtained from Monash Animal Research Facility and allowed to mate. Experiments were validated across 2 independent animal facilities (Monash University, Clayton Campus, and Baker Heart and Diabetes Institute) and laboratories (F.Z.M. and D.M.K.). Histone deacetylase 9 knockout (HDAC9-/-) mice were obtained from a local colony maintained by Prof Charles Mackay at Monash University (approval number 17322). All animal experiments were approved by Monash Animal Ethics Committee (approval number 17465) and the Alfred Medical Research and Education Precinct Animal Experimentation Ethics Committee (approval number E/1626/2016/B) in compliance with guidelines by the National Medical and Health Research Council of Australia.

**Dietary intervention**

Female mice were allowed to feed ad libitum for the duration of pregnancy and lactation. We used diets high in resistant starches, a prebiotic type of fibre (referred to as "high-fibre", SF11-025) or low in resistant starches ("low-fibre", SF09-028) both obtained from Specialty Feeds (see Supplementary Table 1 for nutritional information). Once offspring reached 3-weeks of age (weaning age), all animals were placed on a standard chow from Specialty Feeds. Food was replenished twice a week for the duration of the intervention. All animals were kept in clean breeding facility at Monash University followed by specific pathogen-free facilities.

**Minipump implantation**

Prior to minipump implantation, littermates from the same maternal diet (low-fibre or high-fibre) were randomised to sham (0.9% saline) or Angiotensin II (0.25 mg·kg-1·d-1 Ang II; Auspep) treatment using an automated Excel spreadsheet (developed by K.W). ALZET osmotic minipump (model 2004) were prepared and primed for 40 hours before implanting subcutaneously under isoflurane anaesthesia in the lower flank.

**Characterisation of the cardiovascular system**

Blood pressure (BP) was measured using the CODA High Throughput Non-invasive Tail cuff system (Kent Scientific Corporation) in a quiet room. Measurements and analysis were performed according to manufactures protocol. Briefly, animals underwent 3 consecutive days of acclimatisation where they were placed in restrainers for 10 full measurement cycles. Care was taken to ensure animals were not in distress before, during or after measurements to accurately reflect resting BP. Baseline BP measurements were obtained one day prior to minipump surgeries using the same protocol of 10 full measurement cycles. Following minipump implantation all animals recovered for a full week before subsequent weekly BP was measured over a 4-week period. Data was analysed using CODA software and reported as change over time per treatment group. Echocardiography was performed in the final week using the Visualsonics Vevo 2100 Ultra High Frequency ultrasound imaging system and software. Mice were anaesthetised using 1-2% isoflurane and parasternal long axis, short axis and M-mode images were obtained by an experienced technician. In an independent cohort and institute, cardiac catherization was performed under isoflurane using a 1.4F Mikro-Tip pressure catheter (Millar) to obtain left ventricular pressure.

**Tissue collection**

Following euthanasia cardiac puncture was performed to remove blood. Right atria were snipped and organs were perfused with 10ml cold PBS. Organs were then rapidly removed and either processed for histology (paraffin, OCT) or snap frozen in liquid nitrogen and stored at -80^o^C for future analysis. Intestinal content was rapidly removed and snap frozen and stored at -80^o^C until DNA extraction or short-chain fatty acids (SCFAs) measurement. Blood was collected in heparinised collection tubes tubes and centrifugated for 10 minutes. Plasma was collected, place in clean tubes and stored in -80^o^C until short-chain fatty acids (SCFAs) measurement were performed.

**Histological staining and analysis**

After dissection, heart was cut through the transverse axis. The apical portion of the heart was snap frozen and remaining tissue was placed in 10% formaldehyde and paraffin wax embedded. Small intestinal tissue was divided into three equal segments, cut open, gently rinsed with PBS to remove gut content and rolled into Swiss rolls. After removal of caecal content, caecal tissue was rinsed with PBS and rolled into Swiss rolls. Intestinal tissue was then placed in 10% formaldehyde and paraffin wax embedded. Heart and intestinal tissue were sliced into 4 μm thick sections and stained for collagen (Masson's trichrome). Sequential 4 μm thick sections of gut tissue were also stained for goblet cells (Alcian blue/periodic acid–Schiff AB/PAS). Whole tissue sections were scanned using the Scanscope AT Turbo (Aperio) at 40X magnification. Total collagen in the heart tissue was quantified using a colour thresholding macro (developed by C.J.) using ImageJ software (FIJI). Collagen levels were expressed as a percentage of the total area of heart tissue scanned. Intestinal fibrosis was quantified manually using FIJI ImageJ software, with each intestinal tissue section was imaged three times in different fields of view. All images analysed were captured at 10x magnification using the ImageScope software (Aperio), v12.4.0.5043). Ten consecutive measurements per image were taken to quantify fibrotic area (thickness of collagen deposition) within the submucosa. An average thickness was then obtained per field of view (average of ten measurements). An overall average of the three fields of view was then taken per tissue section which was then reported as fibrosis thickness.

**Quantitative Real-time PCR (qPCR)**

To quantify gene expression, RNA was extracted from frozen heart tissue using TRIzol Reagent (Applied Biosystems) according to manufacturer specifications, and was DNase treated. RNA quality and concentration were checked using a Nanodrop machine. RNA concentration was normalised across samples prior to reverse transcription of cDNA using the High Capacity cDNA RT kit (Thermo Fisher Scientific), primers used and PCR conditions are as previously published.^1,2^ Gene expression was measured using Fast SYBRGreen method on QuantStudio 7 Instrument (Thermo Fisher Scientific). Expression was reported as relative abundance to the housekeeping gene *Gapdh* using the 2^-ΔCt^ method.

**Caecal DNA extraction, library preparation and 16S sequencing**

Caecal DNA (0.1g) was extracted using the DNeasy PowerSoil DNA isolation kit (Qiagen). DNA libraries were prepared by PCR amplification of the V4 region of bacterial 16S ribosomal RNA as previously described^1,2^ using 20 ng of caecal DNA, 515F and 926R primers and Platinum Hot Start PCR master mix in a Thermal Cycler (BioRad). DNA libraries were checked for concentration on a Qubit. Two hundred and forty ng of each library were sequenced in an Ilumina MiSeq sequencer (300bp paired-end reads).

**16S rRNA microbiome data analysis**

Sequence reads from samples were first analysed using the QIIME2 framework.^3^ Forward and reverse reads were first truncated at base number 243 for forward reads and base number 224 for reverse reads, then were denoised, merged, and chimera filtered using the DADA2 plugin^4^, resulting in an amplicon sequence variant (ASV) table (via q2-dada2) with resolution at the single nucleotide level. A phylogeny was then created using fasttree2^5^ from mafft-aligned^6^ ASVs, and were then subsampled without replacement to 29,000 reads per sample (via q2-alignment and q2-phylogeny). β diversity metrics were generated from the rarefied samples (via q2-diversity), reported here as weighted Unifrac metrics along with associated Principle Coordinate Analysis (PCoA) distance tables. α-diversity metrics, including Chao1 and Shannon indices were analysed using MicrobiomeAnalyst.^7,8^ Linear discriminant analysis (LDA) effect size (LEfSe) corrected for multiple adjustments using FDR q<0.05 was performed on MicrobiomeAnalyst, together with edgeR using FDR q<0.05 to compare between low and high fibre offspring.

**Shotgun metagenome sequencing**

Given there were no differences between sham and Ang II treatment in the gut microbiome phenotype (Figure 2B), we utilised sham low-fibre and high-fibre for shotgun metagenome sequencing. Metagenomes were sequenced in an Illumina NovaSeq instrument with 30 million/reads per sample. FASTQ files were first inspected with FastQC (version 0.11.9).^9^ Trimmomatic (version 0.38) was then used with the paired-end flag for Illumina adapter trimming, leading and trailing removal of bases with a quality score below 3, and for cutting each read using a 4-base wide sliding window where the average per-base quality dropped below 15.^10^ Contig and scaffold assembly was conducted using metaSPAdes (version 3.13.1) with 96 GB of RAM and 64 threads.^11^ Separately, FASTQ files were passed through HumanN2, supported by Bowtie2 (version 2.3.5) and the TBB library (version 20180312oss).^12,13^ HumanN2 was run with the protein-database, nucleotide-database, and metaphlan-options flags, pointing to the UniRef, chocoPhlan, and MetaPhlAn databases downloaded with the included humann_databases utility script. The humann_rename_table utility script was used to reattach full gene and pathway names, and the humann_renorm_table script was used to convert reads per kilobases into normalised copies per million count. The gene count outputs were further condensed by grouping genes by level-4 Enzyme Commission categories using the humann_regroup_table utility script. Output files were then passed to R (version 4.0.3), where gene counts, level-4 Enzyme Commission category-grouped genes, and pathway abundance counts were analysed for differential expression between both maternal diet and treatment using the limma package (version 3.46.0), supported by the Tidyverse package (version 1.3.1).^14-17^ Differential abundance of taxa was analysed using the ALDEx2 package (version 1.22.0) using the main aldex function with "zero" as the set of features to retain as the denominator for the subsequent geometric mean calculation.^18^ PCA plots and volcano plots were then created using ggplot2 (version 3.3.5).^19^

**Short-chain fatty acids quantification by LCMS**

*Plasma extraction and derivatization*

Fresh stock solutions 5 mg/ml AA, BA, PA, IBA, IVA, VA, CA in water, 200 mM EDC in 50% acetonitrile, 200 mM 3-NPH in 50% acetonitrile and 100 mM 13C6-3-NPH in 50% acetonitrile were prepared on the day and kept covered from light where possible. SCFAs were mixed together in a 10:1 ratio (AA, BA, PA: IBA, IVA, VA, CA) to match plasma SCFA levels. Mixed working solution 100/10 µg/ml was prepared for ISTD derivatisation, QC sample spiking and calibrant spiking: AA, BA and PA were mixed together to make 1 mg/ml solution. IBA, IVA, VA and CA were mixed together to make 1 mg/ml solution which is then diluted 10x with water to make 100 µg/ml. 80 µL of this solution and 1 mg/ml AA, BA and PA solution and 640 µL of water were mixed together to make mixed SCFA 100/10 µg/ml solution which was then further diluted with acetonitrile to make 50/5 µg/ml, 10/1 µg/ml, 1/0.1 µg/ml solutions. QC samples were prepared by spiking 20 µl of pooled mice plasma before extraction with 0.5/0.05 µg/ml and 1/0.1 µg/ml SCFA and adjusting acetonitrile volume to total of 60 µL. An unspiked QC sample was also extracted to determine natural SCFA level in QC samples. Calibrants were prepared by spiking 20 µL of water with 1/0.1 µg/ml, 50/5 µg/ml, 100/10 µg/ml, 150/15 µg/m, 200/20 µg/ml and adjusting acetonitrile volume to total 60 µL. These samples were mixed together with the study samples in a cool room for 30 min. 10 µL of plasma (study samples only) extracted with 30 µl of acetonitrile and mixed in a cool room for 30 min to crush out protein. All samples were centrifuged and 20 µL of supernatant were transferred to new tubes (equal to 5 µL of plasma). Derivatising reagents were added 40 uL 7.5% pyridine solution, 20 µL 200 mm EDC, 20 µL 200 mM 3-NPH and mixed at room temp covered from light for 45 min. Alongside ISTD was derivatised using 13C-3-NPH. 200 µL of 100/10 µg/ml with 80 µL of EDC and 80 µL of 13C6-3-NPH and 160 µL pyridine solution. After derivatisation 480 µL of 50% acetonitrile was added to dilute to final 1000 µL which produced 20/2 µg/ml ISTD solution. After derivatisation was completed, 10 µL of 20/2 µg/ml ISTD solution was added followed by 90 µl of 0.1% formic acid solution, vortexed, quickly centrifuged and transferred to vials. 6 µL was injected onto LCMS. This produced 40/4 µg/ml ISTD spike in plasma.

*Caecal content SCFAs analysis*

Caecal content SCFAs analysis performed in a similar fashion with some changes in concentrations for calibrants, QC and ISTD to better match caecal content SCFA levels. SCFA were mixed together in a ratio 1000 (AA, BA) : 500 (PA) : 20 (IBA, VA) : 10 (IVA) : 1 (CA). Briefly, 20-40 mg of caecal content was recorded and weighed into Eppendorf tubes and extracted with 6 µL of acetonitrile per 1 mg caecal content. The samples were shaken at 4^o^C for 30 min, centrifuged and 30 µL of supernatant (equivalent to 5 mg of caecal content) was transferred to new Eppendorf tubes. For QC pooled caecal content was extracted and spiked with 150, 600 and 2500 µg/ml SCFAs (these concentrations refer to AA while other SCFAs scale down). Calibrants were prepared at 100, 500, 1000, 2000, 4000 and 6000 µg/ml. Study samples, QC samples and calibrants were derivatised with 70 uL 7.5% pyridine solution, 30 uL 200 mm EDC and 30 uL 200 mM 3-NPH. ISTD was prepared alongside and spiked at 200 ug/ml for AA.

*LC-MS analysis*

LCMS data was acquired on Q-Exactive Orbitrap mass spectrometer (Thermo Fisher Scientific) coupled with high-performance liquid chromatography (HPLC) system Dionex Ultimate® 3000 RS (Thermo Fisher Scientific). A C18 chromatographic column Zorbax Eclipse Plus (1.8 µm, 2.1 × 100 mm, Agilent Technologies) was used for separation. The mobile phase (A) was 0.1% formic acid and mobile phase (B) was 0.1% formic acid in acetonitrile, needle wash solution was 50% isopropanol. The gradient program started at 1 % B, kept at 15% B until 2 min, then increased to 45 %B at 11 min, to 100%B at 11.5 min, kept at 100 % B until 13 min, reduced to 15% B at 13.5 min and followed by equilibration at 15 %B until 17 min. The flow rate was 0.3 mL/min, column compartment temperature was set to 40 ºC, and the injection volume was 6 µL. The total run time was 17 min with the diversion of the flow to waste for the first 2 min and the last 4 min of the run. Mass spectrometer operated in parallel reaction monitoring mode (PRM) with inclusion list enabled in negative ion mode at 17.5k resolution at 200 m/z with detection range of 85 to 1250 m/z. Maximum injection time was 100 ms and automatic gain control was set to 2e5. Normalized collision energy was 45 for all SCFA analytes. Electro-spray ionization source (HESI) was set to 4 kV voltage, sheath gas was set to 20, aux gas to 7 and sweep gas to 1 arbitrary unit, capillary temperature 300°C, probe heater temperature 120°C.

*Data analysis*

Raw data was processed using Tracefinder application (Thermo Scientific). Integrating peaks areas of 137.0357 m/z for natural SCFA and 143.0558 m/z for their internal standards produced by respective deprotonated precursor ions within 10 ppm mass error window. QC of the measurement was assessed by measuring plasma spiked with variable amounts of SCFA in duplicate (Unspiked, 0.5/0.05 ug/ml, 1/0.1 ug/ml and 5/0.5 ug/ml). Plasma contains endogenous SCFA, therefore first endogenous concentrations of SCFAs were determined from Unspiked sample using calibration curve (1/0.1 – 200/20 ng/ml, weighting 1/x). Endogenous concentrations and spiked concentrations were summed to workout total concentrations of SCFAs in QC samples. Accuracy of the QC samples were within ±15% difference. Area ratio of target analyte and internal standard was used to construct standard curves in the range 1-200 ug/ml for AA, PA and BA and 100-20000 ng/ml for IBA, IVA, VA and CA. 1/x weighting was used.

**Flow cytometry**

Animals were euthanised with CO_2_ inhalation and promptly perfused with cold 0.9 mM CaCl_2_ in PBS through the left ventricle. Hearts were harvested and cleaned from atria and connective tissue and minced with curved scissors. Enzymatic digestion of hearts was done in 3 mL of 2 mg/mL collagenase IV (LS004188, Worthington Biochem) and 1 mg/mL dispase II (04942078001, Roche) at 37°C for a duration of 45 minutes with trituration at the 15-minute time points. Following digestion, cell suspension was filtered through 70 µm mesh into 15 mL conical bottom tubes and subjected to a debris clearance spin at 200×g, at 4°C for 15 minutes with no breaks. Supernatant aspirated, and cell pellet washed with 2% F.C.S. (heat inactivated) and 0.9 mM CalCl_2_ in 1XHBSS. Cells were stained with an antibody cocktail which included anti: I-A/I-E (2G9, BD Biosciences), CD11b (M1/70, BD Biosciences), CD64 (a & b alloantigens, X54-5/7.1, BD Biosciences), CD146 (ME-9F1, BD Biosciences), CD31 (390, BD Biosciences), Ly6C (HK1.4, Biolegend), CD59a (REA287, Miltenyi Biotec), Ly6G (1A8, Biolegend), NK1.1 (PK136, Biolegend), CD39 (Duha59, Biolegend), CD90.2 (30-H12, BD Biosciences) and CD45 (30-F11, BD Biosciences). SYTOX™ Green Dead Cell Stain (S34860, Invitrogen) and eBioscience™ Calcein Blue AM Viability Dye (65-0855-39, Invitrogen) were used to gate live and metabolically active cells. Cells were acquired on the BD LSR Fortessa-X20 and analysed using FlowJo(version 10.5.3).

**Single-Cell Library Preparation**

Animals were euthanised in CO_2_ chamber and hearts were quickly harvested, atria were carefully removed and hearts were weighed. Hearts were clamped and attached to a 3D printed perfusion device and continuously perfuse with enzymatic digestion containing 3 ml of 2mg/mL collagenase IV (LS004188, Worthington Biochem) and 1 mg/ml dispase II (04942078001, Roche) in 0.9 mM CaCl2 in PBS. Tissue was incubated at 37°C for a total of 45 min with trituration using wide bore tips at 15-minute intervals. Post-incubation, cell suspension was filtered into 10 mL PBS in 15 ml conical tube. Debris was cleared through centrifugation at 200×g, at 4°C for 15 minutes. Supernatant aspirated, and cell pellet washed with a buffer containing 2% F.C.S. (heat inactivated) and 0.9 mM CalCl2 in 1XHBSS. Cells were stained with Sytox™ Green (S34860, Invitrogen) and Vybrant™ DyeCycle™ (V10273, Invitrogen) with the addition of verapamil and sorted for live, metabolically active and nucleated cells using FACSAria™ III cell sorter (B.D.). Libraries were prepared using according to the 10x Genomics chemistry 3` v2 kit and sequenced in an Illumina HiSeq instrument.

**Analysis of single-cell RNA-Seq data**

Analysis was performed using 10X Genomics pipeline. Briefly, raw sequencing data files by converting basecall files to fastq format, aligning sequencing reads to mm10 transcriptome and quantifying the expression of each transcript in each cell using Cell ranger version 3.1.0 (10X Genomics). Data was further analysed and filtered under the following criteria: <200 unique genes expressed, >10,000 UMIs, or >10% of reads mapping to mitochondria using Seurat R package (version 3.2.0.5). Default normalisation and scaling parameters were used to assess highly variable genes. To visualise high dimensional clustering of cells t-distributed stochastic neighbour embedding (t-SNE) was used. Identified clusters were manually annotated based on features corresponding to canonical cell-type genes (Supplementary Table 5).

To interrogate and identify differentially expressed genes, at least 10% of cells in at least one of the groups being compared were first filtered. The differential expression (D.E.) testing method MAST with cellular detection rate as covariate (MASTcpmDetRate) was used to identify D.E. genes between groups. Statistical significance was determined using the uncorrected p value cut-off 0.01.

Gene Ontology (GO) enrichment analysis for differentially expressed gene lists (uncorrected p <0.01) was performed using the enrichGO function from ClusterProfiler R package version 3.12.0. Data was mapped and annotated using data from org.Mm.eg.db: Genome wide annotation for Mouse, R package version 3.8.2. Data were further filtered using the list of genes identified in the experiment as the background gene list for Mus musculus with minimum and maximum gene set sizes 10 and 500, respectively, to determine G.O. Biological Process terms (GO-BP). Semantic similarity higher than the assigned cut-off of 0.7 were treated as redundant and discarded using clusterProfiler R package. The Benjamini-Hochberg adjusted p-value cut-off of 0.05 was used to determine statistically significant GO-BP terms.

**Chromatin immunoprecipitation (ChIP) assay**

ChIP assay from mouse heart was performed as described previously.^20,21^ Mouse left ventricles were cut into tiny pieces in ice cold PBS and chromatin was fixed for 10 minutes using 1% formaldehyde. Excessive formaldehyde was quenched using 0.125 mM Glycine for 10 minutes at room temperature. Tissues were washed and homogenised in 1x PBS followed by lysis in SDS lysis buffer. Lysates were sonicated to 300-500 bp fragments and the sonication (Qsonica, Q-700) pattern was checked using MultiNA capillary electrophoresis system (Shimadzu). Approximately 5 ug of chromatin was used for immunoprecipitation using antibody specific to H3K9/14 acetylation (cat#06-599 Millipore). Non-specific IgG was incubated simultaneously to each reaction as a control. ChIP enriched DNA was purified using DNA binding columns (MN,Nucleospin) according to the manufacturer’s protocol . Relative enrichment of DNA in input, ChIP and IgG samples were quantified using Fast SYBR Green qPCR system (Applied biosystems) using primers specific to *Nppa* gene promoter region (mNppa ChIP For: GGCTTCCTGGCTGACTTCAT, mNppa ChIP Rev: CCCACCCTAGATGTCCCTGT).

**Statistical analysis**

GraphPad Prism (version 9) package was used for the statistical analysis and graphing. Normal distribution of data was verified using Shapiro-Wilk's normality test. Independent sample t-test was used to compare 2 groups. Two-factor analysis of variance (2-way ANOVA with Benjamini and Hochberg's false discovery rate adjustment for multiple comparisons) not for repetitive measures was used to compare the data between the treatment groups (sham and Ang II) and maternal diet (high-fibre and low-fibre). Repetitive measures analyses were used for tail-cuff BP only. All values are represented as mean±SEM, and P<0.05 (or FDR adjusted P<0.05 for genomic comparisons) were considered significant.

**Supplementary Tables**

**Supplementary Table 1.** Nutrient break down of diets used in study (supplied by Specialty Feeds).

|  | **High-fibre** | **Low-fibre** | **Standard Chow** |
| --- | --- | --- | --- |
| **Crude fibre** | 9.7 % | 0 % | 5.2% |
| **Acid detergent fibre** | 9.7 % | 0 % | 7.7% |
| **Protein** | 19.4 % | 19 % | 19.0% |
| **Total Fat** | 7.0% | 7.0% | 4.6% |
| **% Total calculated energy from lipids** | 16.4% | 16.0% | 12.0% |
| **% Total calculated energy from protein** | 21.9% | 21.0% | 23.0% |
| **% Total calculated energy from carbohydrates** | 61.7% (all resistant starches) | 63% (all non-resistant starches) | 65% |
| **Digestible energy** | 16.3 % | 16.9 % | N/A |

**Supplementary Table 2.** Enzymatic pathways identified in low and high fibre offspring.

| Enzyme commission number | logFC | Adjusted P-value | Direction |
| --- | --- | --- | --- |
| 5.4.99.2\|g__Akkermansia.s__Akkermansia_muciniphila | 6 | 0.002 | Low-fibre |
| 2.7.7.7\|g__Akkermansia.s__Akkermansia_muciniphila | 5.6 | 0.002 | Low-fibre |
| UNGROUPED\|g__Akkermansia.s__Akkermansia_muciniphila | 5.1 | 0.002 | Low-fibre |
| 3.2.1.52\|g__Akkermansia.s__Akkermansia_muciniphila | 5.4 | 0.002 | Low-fibre |
| 3.2.1.23\|g__Akkermansia.s__Akkermansia_muciniphila | 5.6 | 0.002 | Low-fibre |
| 3.6.3.14\|g__Akkermansia.s__Akkermansia_muciniphila | 4.9 | 0.002 | Low-fibre |
| 2.7.4.1\|g__Akkermansia.s__Akkermansia_muciniphila | 4.7 | 0.002 | Low-fibre |
| 6.1.1.4\|g__Akkermansia.s__Akkermansia_muciniphila | 4.6 | 0.002 | Low-fibre |
| 3.6.1.1\|g__Akkermansia.s__Akkermansia_muciniphila | 5.4 | 0.002 | Low-fibre |
| 6.1.1.7\|g__Akkermansia.s__Akkermansia_muciniphila | 4.4 | 0.002 | Low-fibre |
| 6.3.5.2\|g__Akkermansia.s__Akkermansia_muciniphila | 4.7 | 0.002 | Low-fibre |
| 2.2.1.9\|g__Akkermansia.s__Akkermansia_muciniphila | 4.2 | 0.002 | Low-fibre |
| 6.3.2.10\|g__Akkermansia.s__Akkermansia_muciniphila | 4.4 | 0.002 | Low-fibre |
| 1.4.4.2\|g__Akkermansia.s__Akkermansia_muciniphila | 4.6 | 0.002 | Low-fibre |
| 5.1.3.8\|g__Akkermansia.s__Akkermansia_muciniphila | 4.2 | 0.002 | Low-fibre |
| 2.7.11.1\|g__Akkermansia.s__Akkermansia_muciniphila | 4.4 | 0.002 | Low-fibre |
| 6.1.1.20\|g__Akkermansia.s__Akkermansia_muciniphila | 4.5 | 0.002 | Low-fibre |
| 2.7.10.2 | 3 | 0.003 | Low-fibre |
| 1.11.1.5 | 4 | 0.003 | Low-fibre |
| 3.6.3.27\|g__Akkermansia.s__Akkermansia_muciniphila | 4.6 | 0.003 | Low-fibre |
| 2.7.1.40\|g__Akkermansia.s__Akkermansia_muciniphila | 4.2 | 0.003 | Low-fibre |
| 6.3.4.4\|g__Akkermansia.s__Akkermansia_muciniphila | 4 | 0.003 | Low-fibre |
| 1.1.1.205\|g__Akkermansia.s__Akkermansia_muciniphila | 4.3 | 0.003 | Low-fibre |
| 4.1.1.70\|g__Akkermansia.s__Akkermansia_muciniphila | 4.6 | 0.003 | Low-fibre |
| 2.7.7.72\|g__Akkermansia.s__Akkermansia_muciniphila | 4.4 | 0.003 | Low-fibre |
| 2.7.1.90\|g__Akkermansia.s__Akkermansia_muciniphila | 4.2 | 0.003 | Low-fibre |
| 3.4.24.70 | 5.4 | 0.004 | Low-fibre |
| 6.1.1.21\|g__Akkermansia.s__Akkermansia_muciniphila | 4.2 | 0.004 | Low-fibre |
| 6.1.1.22\|g__Akkermansia.s__Akkermansia_muciniphila | 4.1 | 0.004 | Low-fibre |
| 2.8.1.10\|g__Akkermansia.s__Akkermansia_muciniphila | 3.7 | 0.004 | Low-fibre |
| 1.6.5.11\|g__Akkermansia.s__Akkermansia_muciniphila | 4.5 | 0.005 | Low-fibre |
| 5.2.1.8\|g__Akkermansia.s__Akkermansia_muciniphila | 5 | 0.005 | Low-fibre |
| 2.4.1.129\|g__Akkermansia.s__Akkermansia_muciniphila | 5.2 | 0.005 | Low-fibre |
| 4.2.1.3\|g__Akkermansia.s__Akkermansia_muciniphila | 5 | 0.006 | Low-fibre |
| 4.1.1.68 | 3.1 | 0.007 | Low-fibre |
| 1.8.1.2 | 3 | 0.007 | Low-fibre |
| 2.1.1.14 | 4.3 | 0.007 | Low-fibre |
| 4.1.1.68\|g__Akkermansia.s__Akkermansia_muciniphila | 3.1 | 0.007 | Low-fibre |
| 6.3.5.3\|g__Akkermansia.s__Akkermansia_muciniphila | 4.8 | 0.007 | Low-fibre |
| 4.2.1.9\|g__Akkermansia.s__Akkermansia_muciniphila | 4.8 | 0.007 | Low-fibre |
| 5.1.1.1\|g__Akkermansia.s__Akkermansia_muciniphila | 4.8 | 0.007 | Low-fibre |
| 6.3.4.3\|g__Akkermansia.s__Akkermansia_muciniphila | 4.8 | 0.007 | Low-fibre |
| 2.4.1.227\|g__Akkermansia.s__Akkermansia_muciniphila | 4.7 | 0.007 | Low-fibre |
| 2.1.2.1\|g__Akkermansia.s__Akkermansia_muciniphila | 4.9 | 0.007 | Low-fibre |
| 3.6.4.12\|g__Akkermansia.s__Akkermansia_muciniphila | 4.9 | 0.007 | Low-fibre |
| 6.1.1.11\|g__Akkermansia.s__Akkermansia_muciniphila | 4.7 | 0.007 | Low-fibre |
| 2.7.1.51\|g__Akkermansia.s__Akkermansia_muciniphila | 4.6 | 0.007 | Low-fibre |
| 1.6.99.5\|g__Akkermansia.s__Akkermansia_muciniphila | 4.6 | 0.007 | Low-fibre |
| 2.7.1.11\|g__Akkermansia.s__Akkermansia_muciniphila | 4.7 | 0.007 | Low-fibre |
| 1.11.1.5\|g__Akkermansia.s__Akkermansia_muciniphila | 4.7 | 0.007 | Low-fibre |
| 2.4.1.1\|g__Akkermansia.s__Akkermansia_muciniphila | 5 | 0.007 | Low-fibre |
| 6.3.2.9\|g__Akkermansia.s__Akkermansia_muciniphila | 4.5 | 0.007 | Low-fibre |
| 3.6.5.n1\|g__Akkermansia.s__Akkermansia_muciniphila | 4.5 | 0.007 | Low-fibre |
| 2.7.10.2\|g__Akkermansia.s__Akkermansia_muciniphila | 4.5 | 0.007 | Low-fibre |
| 1.1.1.42\|g__Akkermansia.s__Akkermansia_muciniphila | 4.7 | 0.007 | Low-fibre |
| 2.8.1.8\|g__Akkermansia.s__Akkermansia_muciniphila | 4.5 | 0.007 | Low-fibre |
| 3.2.1.51\|g__Akkermansia.s__Akkermansia_muciniphila | 4.4 | 0.007 | Low-fibre |
| 2.1.3.3\|g__Akkermansia.s__Akkermansia_muciniphila | 4.2 | 0.007 | Low-fibre |
| 2.4.2.18\|g__Akkermansia.s__Akkermansia_muciniphila | 4.3 | 0.007 | Low-fibre |
| 2.5.1.3\|g__Akkermansia.s__Akkermansia_muciniphila | 4.4 | 0.007 | Low-fibre |
| 4.2.1.46\|g__Akkermansia.s__Akkermansia_muciniphila | 4.5 | 0.007 | Low-fibre |
| 1.1.1.86\|g__Akkermansia.s__Akkermansia_muciniphila | 4.6 | 0.007 | Low-fibre |
| 4.2.1.20\|g__Akkermansia.s__Akkermansia_muciniphila | 4.5 | 0.007 | Low-fibre |
| 4.2.99.18\|g__Akkermansia.s__Akkermansia_muciniphila | 4.5 | 0.007 | Low-fibre |
| 1.3.99.22\|g__Akkermansia.s__Akkermansia_muciniphila | 4.4 | 0.008 | Low-fibre |
| 4.2.1.11\|g__Akkermansia.s__Akkermansia_muciniphila | 4.3 | 0.008 | Low-fibre |
| 6.1.1.6\|g__Akkermansia.s__Akkermansia_muciniphila | 4.3 | 0.008 | Low-fibre |
| 1.1.1.37\|g__Akkermansia.s__Akkermansia_muciniphila | 4.5 | 0.008 | Low-fibre |
| 1.8.1.4\|g__Akkermansia.s__Akkermansia_muciniphila | 4 | 0.008 | Low-fibre |
| 3.4.21.92\|g__Akkermansia.s__Akkermansia_muciniphila | 4 | 0.008 | Low-fibre |
| 2.7.8.6 | 4.2 | 0.009 | Low-fibre |
| 2.5.1.55\|g__Parabacteroides.s__Parabacteroides_goldsteinii | 2.2 | 0.009 | Low-fibre |
| 3.2.1.22\|g__Akkermansia.s__Akkermansia_muciniphila | 4.2 | 0.009 | Low-fibre |
| 4.3.2.1\|g__Akkermansia.s__Akkermansia_muciniphila | 4.3 | 0.009 | Low-fibre |
| 1.17.1.2\|g__Akkermansia.s__Akkermansia_muciniphila | 4.4 | 0.009 | Low-fibre |
| 2.5.1.47\|g__Akkermansia.s__Akkermansia_muciniphila | 4.2 | 0.009 | Low-fibre |
| 3.1.26.3\|g__Akkermansia.s__Akkermansia_muciniphila | 4 | 0.009 | Low-fibre |
| 1.3.99.1\|g__Akkermansia.s__Akkermansia_muciniphila | 4.7 | 0.009 | Low-fibre |
| 4.1.1.15\|g__Akkermansia.s__Akkermansia_muciniphila | 4.1 | 0.009 | Low-fibre |
| 3.1.13.1\|g__Akkermansia.s__Akkermansia_muciniphila | 4 | 0.009 | Low-fibre |
| 1.1.1.271\|g__Akkermansia.s__Akkermansia_muciniphila | 4.5 | 0.009 | Low-fibre |
| 2.3.1.79\|g__Akkermansia.s__Akkermansia_muciniphila | 3.4 | 0.009 | Low-fibre |
| 2.7.8.6\|g__Akkermansia.s__Akkermansia_muciniphila | 4.2 | 0.009 | Low-fibre |
| 4.1.2.13\|g__Akkermansia.s__Akkermansia_muciniphila | 4.1 | 0.009 | Low-fibre |
| 2.5.1.49\|g__Akkermansia.s__Akkermansia_muciniphila | 4 | 0.009 | Low-fibre |
| 6.1.1.15\|g__Akkermansia.s__Akkermansia_muciniphila | 4.1 | 0.009 | Low-fibre |
| 4.1.99.1\|g__Akkermansia.s__Akkermansia_muciniphila | 4 | 0.009 | Low-fibre |
| 6.2.1.5\|g__Akkermansia.s__Akkermansia_muciniphila | 4.1 | 0.009 | Low-fibre |
| 2.4.1.182\|g__Akkermansia.s__Akkermansia_muciniphila | 4 | 0.009 | Low-fibre |
| 1.8.1.2\|g__Akkermansia.s__Akkermansia_muciniphila | 4.1 | 0.009 | Low-fibre |
| 4.1.1.19\|g__Akkermansia.s__Akkermansia_muciniphila | 4.2 | 0.009 | Low-fibre |
| 2.6.1.44 | 3.8 | 0.01 | Low-fibre |
| 2.4.2.19 | 2.8 | 0.01 | Low-fibre |
| 2.3.1.234\|g__Akkermansia.s__Akkermansia_muciniphila | 3.9 | 0.01 | Low-fibre |
| 4.2.3.4\|g__Akkermansia.s__Akkermansia_muciniphila | 3.7 | 0.01 | Low-fibre |
| 2.6.1.44\|g__Akkermansia.s__Akkermansia_muciniphila | 3.8 | 0.01 | Low-fibre |
| 4.3.2.2\|g__Akkermansia.s__Akkermansia_muciniphila | 4 | 0.01 | Low-fibre |
| 4.2.1.47\|g__Akkermansia.s__Akkermansia_muciniphila | 3.8 | 0.01 | Low-fibre |
| 4.1.1.48\|g__Akkermansia.s__Akkermansia_muciniphila | 4.1 | 0.01 | Low-fibre |
| 2.7.7.4\|g__Akkermansia.s__Akkermansia_muciniphila | 3.9 | 0.01 | Low-fibre |
| 2.7.1.2\|g__Bacteroides.s__Bacteroides_ovatus | -4.6 | 0.011 | High-fibre |
| 3.5.4.25\|unclassified | 2.7 | 0.011 | Low-fibre |
| 3.5.4.25 | 2.7 | 0.011 | Low-fibre |
| 3.4.23.36\|g__Akkermansia.s__Akkermansia_muciniphila | 3.9 | 0.011 | Low-fibre |
| 6.1.1.17\|g__Akkermansia.s__Akkermansia_muciniphila | 3.6 | 0.011 | Low-fibre |
| 6.3.2.4\|g__Akkermansia.s__Akkermansia_muciniphila | 3.9 | 0.011 | Low-fibre |
| 2.7.1.130\|g__Akkermansia.s__Akkermansia_muciniphila | 4.2 | 0.011 | Low-fibre |
| 3.5.1.2\|g__Akkermansia.s__Akkermansia_muciniphila | 3.7 | 0.011 | Low-fibre |
| 3.5.3.12\|g__Akkermansia.s__Akkermansia_muciniphila | 3.7 | 0.011 | Low-fibre |
| 2.7.8.13\|g__Akkermansia.s__Akkermansia_muciniphila | 4.1 | 0.011 | Low-fibre |
| 1.3.5.2\|g__Akkermansia.s__Akkermansia_muciniphila | 3.7 | 0.011 | Low-fibre |
| 6.3.4.2\|g__Akkermansia.s__Akkermansia_muciniphila | 4.1 | 0.011 | Low-fibre |
| 2.1.1.198\|g__Akkermansia.s__Akkermansia_muciniphila | 3.7 | 0.011 | Low-fibre |
| 4.2.1.59\|g__Akkermansia.s__Akkermansia_muciniphila | 3.7 | 0.011 | Low-fibre |
| 2.5.1.78\|g__Akkermansia.s__Akkermansia_muciniphila | 3.5 | 0.011 | Low-fibre |
| 2.1.1.72\|g__Akkermansia.s__Akkermansia_muciniphila | 3.5 | 0.012 | Low-fibre |
| 2.1.1.199\|g__Akkermansia.s__Akkermansia_muciniphila | 3.7 | 0.012 | Low-fibre |
| 2.3.1.180\|g__Akkermansia.s__Akkermansia_muciniphila | 3.8 | 0.013 | Low-fibre |
| 1.1.1.103\|g__Akkermansia.s__Akkermansia_muciniphila | 3.6 | 0.014 | Low-fibre |
| 2.7.7.87\|g__Akkermansia.s__Akkermansia_muciniphila | 3.8 | 0.014 | Low-fibre |
| 3.6.1.19\|g__Akkermansia.s__Akkermansia_muciniphila | 3.8 | 0.014 | Low-fibre |
| 6.1.1.19\|g__Akkermansia.s__Akkermansia_muciniphila | 3.7 | 0.014 | Low-fibre |
| 2.7.7.38\|g__Akkermansia.s__Akkermansia_muciniphila | 3.5 | 0.014 | Low-fibre |
| 5.4.99.12\|g__Akkermansia.s__Akkermansia_muciniphila | 3.7 | 0.014 | Low-fibre |
| 6.3.4.5\|g__Akkermansia.s__Akkermansia_muciniphila | 3.5 | 0.014 | Low-fibre |
| 2.1.1.163\|g__Akkermansia.s__Akkermansia_muciniphila | 3.3 | 0.014 | Low-fibre |
| 2.2.1.6\|g__Akkermansia.s__Akkermansia_muciniphila | 3.8 | 0.014 | Low-fibre |
| 2.1.1.228\|g__Akkermansia.s__Akkermansia_muciniphila | 3.5 | 0.015 | Low-fibre |
| 2.7.1.50\|g__Akkermansia.s__Akkermansia_muciniphila | 3.4 | 0.015 | Low-fibre |
| 4.1.1.65\|g__Akkermansia.s__Akkermansia_muciniphila | 3.5 | 0.015 | Low-fibre |
| 1.3.5.2 | 3.3 | 0.016 | Low-fibre |
| 2.7.7.72 | 2.5 | 0.016 | Low-fibre |
| 2.1.2.11\|g__Akkermansia.s__Akkermansia_muciniphila | 3.2 | 0.016 | Low-fibre |
| 5.4.2.12\|g__Akkermansia.s__Akkermansia_muciniphila | 3.8 | 0.016 | Low-fibre |
| 3.5.2.6\|g__Akkermansia.s__Akkermansia_muciniphila | 3.4 | 0.017 | Low-fibre |
| 4.2.3.5\|g__Akkermansia.s__Akkermansia_muciniphila | 3.5 | 0.017 | Low-fibre |
| 2.5.1.72\|g__Akkermansia.s__Akkermansia_muciniphila | 3.6 | 0.017 | Low-fibre |
| 6.1.1.16\|g__Akkermansia.s__Akkermansia_muciniphila | 3.9 | 0.017 | Low-fibre |
| 4.1.1.32\|g__Akkermansia.s__Akkermansia_muciniphila | 3.6 | 0.018 | Low-fibre |
| 1.1.1.85\|g__Akkermansia.s__Akkermansia_muciniphila | 3.4 | 0.018 | Low-fibre |
| 5.3.1.13\|g__Akkermansia.s__Akkermansia_muciniphila | 3.6 | 0.019 | Low-fibre |
| 5.1.3.1\|g__Akkermansia.s__Akkermansia_muciniphila | 3.2 | 0.019 | Low-fibre |
| 2.4.99.17\|g__Akkermansia.s__Akkermansia_muciniphila | 3.2 | 0.019 | Low-fibre |
| 3.4.13.22\|g__Akkermansia.s__Akkermansia_muciniphila | 3.2 | 0.019 | Low-fibre |
| 2.3.1.129\|g__Akkermansia.s__Akkermansia_muciniphila | 3.8 | 0.02 | Low-fibre |
| 2.8.4.4\|g__Akkermansia.s__Akkermansia_muciniphila | 3.3 | 0.02 | Low-fibre |
| 2.7.4.8\|g__Akkermansia.s__Akkermansia_muciniphila | 3.1 | 0.02 | Low-fibre |
| 3.1.22.4\|g__Akkermansia.s__Akkermansia_muciniphila | 3.1 | 0.02 | Low-fibre |
| 2.4.2.17\|g__Akkermansia.s__Akkermansia_muciniphila | 3.4 | 0.02 | Low-fibre |
| 3.5.3.12 | 3.1 | 0.022 | Low-fibre |
| 1.8.4.8\|g__Akkermansia.s__Akkermansia_muciniphila | 3.1 | 0.022 | Low-fibre |
| 2.7.7.9\|g__Akkermansia.s__Akkermansia_muciniphila | 3.1 | 0.022 | Low-fibre |
| 4.1.1.36\|g__Akkermansia.s__Akkermansia_muciniphila | 3.4 | 0.022 | Low-fibre |
| 1.4.3.5 | 3.9 | 0.023 | Low-fibre |
| 6.3.2.1\|g__Akkermansia.s__Akkermansia_muciniphila | 3.4 | 0.023 | Low-fibre |
| 2.3.1.39\|g__Akkermansia.s__Akkermansia_muciniphila | 3.4 | 0.023 | Low-fibre |
| 2.6.1.9\|g__Akkermansia.s__Akkermansia_muciniphila | 3.2 | 0.023 | Low-fibre |
| 2.7.3.9\|g__Akkermansia.s__Akkermansia_muciniphila | 3.3 | 0.026 | Low-fibre |
| 2.7.1.51 | 2.6 | 0.027 | Low-fibre |
| 2.7.8.13\|g__Bacteroides.s__Bacteroides_ovatus | -5.4 | 0.029 | High-fibre |
| 2.5.1.54\|g__Akkermansia.s__Akkermansia_muciniphila | 3.1 | 0.029 | Low-fibre |
| 6.3.3.3\|g__Akkermansia.s__Akkermansia_muciniphila | 2.8 | 0.029 | Low-fibre |
| 3.5.2.3\|g__Akkermansia.s__Akkermansia_muciniphila | 3.2 | 0.029 | Low-fibre |
| 2.8.4.3\|g__Akkermansia.s__Akkermansia_muciniphila | 2.9 | 0.029 | Low-fibre |
| 3.1.21.1\|g__Escherichia.s__Escherichia_coli | -3.9 | 0.03 | High-fibre |
| 3.1.21.1 | -3.9 | 0.03 | High-fibre |
| 3.2.2.20 | 2.9 | 0.03 | Low-fibre |
| 6.3.2.6\|g__Akkermansia.s__Akkermansia_muciniphila | 3.4 | 0.03 | Low-fibre |
| 2.4.2.10\|g__Akkermansia.s__Akkermansia_muciniphila | 3 | 0.031 | Low-fibre |
| 1.8.4.8 | 2.6 | 0.032 | Low-fibre |
| 2.2.1.7\|g__Akkermansia.s__Akkermansia_muciniphila | 3 | 0.032 | Low-fibre |
| 2.6.99.2\|g__Akkermansia.s__Akkermansia_muciniphila | 3.2 | 0.032 | Low-fibre |
| 1.3.1.12 | 2.9 | 0.033 | Low-fibre |
| 1.1.1.262\|g__Akkermansia.s__Akkermansia_muciniphila | 3.1 | 0.033 | Low-fibre |
| 2.5.1.19\|g__Akkermansia.s__Akkermansia_muciniphila | 2.7 | 0.04 | Low-fibre |
| 3.1.1.31 | 2 | 0.041 | Low-fibre |
| 6.1.1.6\|g__Lactobacillus.s__Lactobacillus_murinus | -2.4 | 0.043 | High-fibre |
| 5.1.1.7\|g__Akkermansia.s__Akkermansia_muciniphila | 2.7 | 0.044 | Low-fibre |
| 2.3.1.n2\|g__Akkermansia.s__Akkermansia_muciniphila | 3.1 | 0.046 | Low-fibre |

**Supplementary Table 3.** Pathway analysis of microbial pathways enriched in low and high fibre offspring.

| Gene family description | UniRefID | logFC | Adjusted P-value | Direction |
| --- | --- | --- | --- | --- |
| Mobilization protein\|g__Bacteroides.s__Bacteroides_ovatus | W0ETG2 | -3.63 | 0.009698 | High-fibre |
| Outer membrane autotransporter barrel domain protein | B2UQY0 | 5.57 | 0.009698 | Low-fibre |
| Outer membrane autotransporter barrel domain protein\|g__Akkermansia.s__Akkermansia_muciniphila | B2UQY0 | 5.57 | 0.009698 | Low-fibre |
| Pyruvate-flavodoxin oxidoreductase | B2UQ98 | 5.31 | 0.009698 | Low-fibre |
| Pyruvate-flavodoxin oxidoreductase\|g__Akkermansia.s__Akkermansia_muciniphila | B2UQ98 | 5.31 | 0.009698 | Low-fibre |
| Glycoside hydrolase family 2 TIM barrel | B2UQC2 | 5.25 | 0.009698 | Low-fibre |
| Glycoside hydrolase family 2 TIM barrel\|g__Akkermansia.s__Akkermansia_muciniphila | B2UQC2 | 5.25 | 0.009698 | Low-fibre |
| Excinuclease ABC A subunit | R7E5C2 | 5.16 | 0.009698 | Low-fibre |
| Excinuclease ABC A subunit\|g__Akkermansia.s__Akkermansia_muciniphila | R7E5C2 | 5.16 | 0.009698 | Low-fibre |
| NO_NAME | B2UMW7 | 4.98 | 0.009698 | Low-fibre |
| NO_NAME\|g__Akkermansia.s__Akkermansia_muciniphila | B2UMW7 | 4.98 | 0.009698 | Low-fibre |
| UniRef90_unknown\|g__Akkermansia.s__Akkermansia_muciniphila | unknown\|g__Akkermansia.s__Akkermansia_muciniphila | 5.07 | 0.009698 | Low-fibre |
| ATP-dependent zinc metalloprotease FtsH | B2UMY1 | 4.89 | 0.009698 | Low-fibre |
| ATP-dependent zinc metalloprotease FtsH\|g__Akkermansia.s__Akkermansia_muciniphila | B2UMY1 | 4.89 | 0.009698 | Low-fibre |
| ABC transporter related | B2UMZ4 | 4.96 | 0.009698 | Low-fibre |
| ABC transporter related\|g__Akkermansia.s__Akkermansia_muciniphila | B2UMZ4 | 4.96 | 0.009698 | Low-fibre |
| UvrD/REP helicase | B2ULJ5 | 5.27 | 0.009698 | Low-fibre |
| UvrD/REP helicase\|g__Akkermansia.s__Akkermansia_muciniphila | B2ULJ5 | 5.27 | 0.009698 | Low-fibre |
| NO_NAME | B2UNB4 | 5.22 | 0.009698 | Low-fibre |
| NO_NAME\|g__Akkermansia.s__Akkermansia_muciniphila | B2UNB4 | 5.22 | 0.009698 | Low-fibre |
| Polyphosphate kinase | B2UQN5 | 4.74 | 0.009698 | Low-fibre |
| Polyphosphate kinase\|g__Akkermansia.s__Akkermansia_muciniphila | B2UQN5 | 4.74 | 0.009698 | Low-fibre |
| Sulfatase | B2UPC5 | 4.73 | 0.009698 | Low-fibre |
| Sulfatase\|g__Akkermansia.s__Akkermansia_muciniphila | B2UPC5 | 4.73 | 0.009698 | Low-fibre |
| NO_NAME | Q8A5E8 | 3.84 | 0.009698 | Low-fibre |
| Leucine--tRNA ligase | B2UM29 | 4.63 | 0.009698 | Low-fibre |
| Leucine--tRNA ligase\|g__Akkermansia.s__Akkermansia_muciniphila | B2UM29 | 4.63 | 0.009698 | Low-fibre |
| Acyltransferase 3 | B2UQ62 | 4.72 | 0.009698 | Low-fibre |
| Acyltransferase 3\|g__Akkermansia.s__Akkermansia_muciniphila | B2UQ62 | 4.72 | 0.009698 | Low-fibre |
| (NiFe) hydrogenase maturation protein HypF | B2UKV0 | 4.65 | 0.009698 | Low-fibre |
| (NiFe) hydrogenase maturation protein HypF\|g__Akkermansia.s__Akkermansia_muciniphila | B2UKV0 | 4.65 | 0.009698 | Low-fibre |
| Organic solvent tolerance protein OstA-like protein | R7DZY8 | 4.64 | 0.009698 | Low-fibre |
| Organic solvent tolerance protein OstA-like protein\|g__Akkermansia.s__Akkermansia_muciniphila | R7DZY8 | 4.64 | 0.009698 | Low-fibre |
| Glycoside hydrolase family 2 sugar binding | R7DVT5 | 4.48 | 0.009698 | Low-fibre |
| Glycoside hydrolase family 2 sugar binding\|g__Akkermansia.s__Akkermansia_muciniphila | R7DVT5 | 4.48 | 0.009698 | Low-fibre |
| Raffinose synthase | B2UNY5 | 4.57 | 0.009698 | Low-fibre |
| Raffinose synthase\|g__Akkermansia.s__Akkermansia_muciniphila | B2UNY5 | 4.57 | 0.009698 | Low-fibre |
| Non-specific serine/threonine protein kinase | B2UQL8 | 4.71 | 0.009698 | Low-fibre |
| Non-specific serine/threonine protein kinase\|g__Akkermansia.s__Akkermansia_muciniphila | B2UQL8 | 4.71 | 0.009698 | Low-fibre |
| Cytochrome C family protein | B2UNA6 | 4.56 | 0.009698 | Low-fibre |
| Cytochrome C family protein\|g__Akkermansia.s__Akkermansia_muciniphila | B2UNA6 | 4.56 | 0.009698 | Low-fibre |
| Glycosyl transferase group 1 | B2UKV5 | 4.59 | 0.009698 | Low-fibre |
| Glycosyl transferase group 1\|g__Akkermansia.s__Akkermansia_muciniphila | B2UKV5 | 4.59 | 0.009698 | Low-fibre |
| Transporter, hydrophobe/amphiphile efflux-1 (HAE1) family | B2ULQ8 | 4.61 | 0.009698 | Low-fibre |
| Transporter, hydrophobe/amphiphile efflux-1 (HAE1) family\|g__Akkermansia.s__Akkermansia_muciniphila | B2ULQ8 | 4.61 | 0.009698 | Low-fibre |
| Alanine--tRNA ligase | B2UKT4 | 4.40 | 0.009698 | Low-fibre |
| Alanine--tRNA ligase\|g__Akkermansia.s__Akkermansia_muciniphila | B2UKT4 | 4.40 | 0.009698 | Low-fibre |
| NO_NAME | B2UNT4 | 4.54 | 0.009698 | Low-fibre |
| NO_NAME\|g__Akkermansia.s__Akkermansia_muciniphila | B2UNT4 | 4.54 | 0.009698 | Low-fibre |
| Alpha-2-macroglobulin domain protein | B2ULJ9 | 4.91 | 0.009698 | Low-fibre |
| Alpha-2-macroglobulin domain protein\|g__Akkermansia.s__Akkermansia_muciniphila | B2ULJ9 | 4.91 | 0.009698 | Low-fibre |
| GMP synthase, large subunit | B2UQ39 | 4.70 | 0.009698 | Low-fibre |
| GMP synthase, large subunit\|g__Akkermansia.s__Akkermansia_muciniphila | B2UQ39 | 4.70 | 0.009698 | Low-fibre |
| Protein-export membrane protein SecD | R6J9D8 | 4.81 | 0.009698 | Low-fibre |
| Protein-export membrane protein SecD\|g__Akkermansia.s__Akkermansia_muciniphila | R6J9D8 | 4.81 | 0.009698 | Low-fibre |
| 2-succinyl-5-enolpyruvyl-6-hydroxy-3-cyclohexene-1-carboxylate synthase | B2UPN5 | 4.24 | 0.009698 | Low-fibre |
| 2-succinyl-5-enolpyruvyl-6-hydroxy-3-cyclohexene-1-carboxylate synthase\|g__Akkermansia.s__Akkermansia_muciniphila | B2UPN5 | 4.24 | 0.009698 | Low-fibre |
| Type II and III secretion system protein | R7E1P2 | 4.44 | 0.009698 | Low-fibre |
| Type II and III secretion system protein\|g__Akkermansia.s__Akkermansia_muciniphila | R7E1P2 | 4.44 | 0.009698 | Low-fibre |
| UDP-N-acetylmuramoyl-tripeptide--D-alanyl-D-alanine ligase | B2UPL4 | 4.37 | 0.009698 | Low-fibre |
| UDP-N-acetylmuramoyl-tripeptide--D-alanyl-D-alanine ligase\|g__Akkermansia.s__Akkermansia_muciniphila | B2UPL4 | 4.37 | 0.009698 | Low-fibre |
| Pseudouridine synthase | R6IWU5 | 4.22 | 0.009698 | Low-fibre |
| Pseudouridine synthase\|g__Akkermansia.s__Akkermansia_muciniphila | R6IWU5 | 4.22 | 0.009698 | Low-fibre |
| Polysaccharide deacetylase | B2ULZ2 | 4.13 | 0.009698 | Low-fibre |
| Polysaccharide deacetylase\|g__Akkermansia.s__Akkermansia_muciniphila | B2ULZ2 | 4.13 | 0.009698 | Low-fibre |
| GTPase Der | B2UMV5 | 4.54 | 0.009698 | Low-fibre |
| GTPase Der\|g__Akkermansia.s__Akkermansia_muciniphila | B2UMV5 | 4.54 | 0.009698 | Low-fibre |
| Peptidase M16 domain protein | R6JAL6 | 4.47 | 0.009698 | Low-fibre |
| Peptidase M16 domain protein\|g__Akkermansia.s__Akkermansia_muciniphila | R6JAL6 | 4.47 | 0.009698 | Low-fibre |
| NADH-quinone oxidoreductase subunit N | B2ULN7 | 4.36 | 0.009698 | Low-fibre |
| NADH-quinone oxidoreductase subunit N\|g__Akkermansia.s__Akkermansia_muciniphila | B2ULN7 | 4.36 | 0.009698 | Low-fibre |
| Glycine dehydrogenase (decarboxylating) | B2UNH4 | 4.66 | 0.009698 | Low-fibre |
| Glycine dehydrogenase (decarboxylating)\|g__Akkermansia.s__Akkermansia_muciniphila | B2UNH4 | 4.66 | 0.009698 | Low-fibre |
| Putative K(+)-stimulated pyrophosphate-energized sodium pump | B2ULG2 | 4.81 | 0.009698 | Low-fibre |
| Putative K(+)-stimulated pyrophosphate-energized sodium pump\|g__Akkermansia.s__Akkermansia_muciniphila | B2ULG2 | 4.81 | 0.009698 | Low-fibre |
| NO_NAME | B2UNI0 | 4.38 | 0.009698 | Low-fibre |
| NO_NAME\|g__Akkermansia.s__Akkermansia_muciniphila | B2UNI0 | 4.38 | 0.009698 | Low-fibre |
| NO_NAME | B2UR19 | 4.11 | 0.009698 | Low-fibre |
| NO_NAME\|g__Akkermansia.s__Akkermansia_muciniphila | B2UR19 | 4.11 | 0.009698 | Low-fibre |
| Glycosyl transferase family 51 | B2UPQ3 | 4.54 | 0.009698 | Low-fibre |
| Glycosyl transferase family 51\|g__Akkermansia.s__Akkermansia_muciniphila | B2UPQ3 | 4.54 | 0.009698 | Low-fibre |
| NO_NAME | B2ULV8 | 4.59 | 0.009698 | Low-fibre |
| NO_NAME\|g__Akkermansia.s__Akkermansia_muciniphila | B2ULV8 | 4.59 | 0.009698 | Low-fibre |
| N-acylglucosamine 2-epimerase | B2UNP4 | 4.22 | 0.009698 | Low-fibre |
| N-acylglucosamine 2-epimerase\|g__Akkermansia.s__Akkermansia_muciniphila | B2UNP4 | 4.22 | 0.009698 | Low-fibre |
| Sulfatase | R6JDP7 | 4.33 | 0.009698 | Low-fibre |
| Sulfatase\|g__Akkermansia.s__Akkermansia_muciniphila | R6JDP7 | 4.33 | 0.009698 | Low-fibre |
| Mg chelatase, subunit ChlI | B2UQ87 | 4.15 | 0.009703 | Low-fibre |
| Mg chelatase, subunit ChlI\|g__Akkermansia.s__Akkermansia_muciniphila | B2UQ87 | 4.15 | 0.009703 | Low-fibre |
| Sodium/hydrogen exchanger | B2URL8 | 4.24 | 0.010462 | Low-fibre |
| Sodium/hydrogen exchanger\|g__Akkermansia.s__Akkermansia_muciniphila | B2URL8 | 4.24 | 0.010462 | Low-fibre |
| Outer membrane autotransporter barrel domain protein | B2UMI4 | 6.08 | 0.011325 | Low-fibre |
| Outer membrane autotransporter barrel domain protein\|g__Akkermansia.s__Akkermansia_muciniphila | B2UMI4 | 6.08 | 0.011325 | Low-fibre |
| Glycosyl hydrolase family 109 protein 2 | B2UQL7 | 4.72 | 0.011343 | Low-fibre |
| Glycosyl hydrolase family 109 protein 2\|g__Akkermansia.s__Akkermansia_muciniphila | B2UQL7 | 4.72 | 0.011343 | Low-fibre |
| Sec-independent protein translocase protein TatC | B2UN92 | 4.01 | 0.011343 | Low-fibre |
| Sec-independent protein translocase protein TatC\|g__Akkermansia.s__Akkermansia_muciniphila | B2UN92 | 4.01 | 0.011343 | Low-fibre |
| NO_NAME | B2ULX3 | 4.41 | 0.011343 | Low-fibre |
| NO_NAME\|g__Akkermansia.s__Akkermansia_muciniphila | B2ULX3 | 4.41 | 0.011343 | Low-fibre |
| Pyruvate kinase | B2UNF7 | 4.23 | 0.011551 | Low-fibre |
| Pyruvate kinase\|g__Akkermansia.s__Akkermansia_muciniphila | B2UNF7 | 4.23 | 0.011551 | Low-fibre |
| Mobilization protein\|g__Bacteroides.s__Bacteroides_xylanisolvens | W0ETG2 | -5.14 | 0.011623 | High-fibre |
| NO_NAME | B2UQ77 | 4.11 | 0.011623 | Low-fibre |
| NO_NAME\|g__Akkermansia.s__Akkermansia_muciniphila | B2UQ77 | 4.11 | 0.011623 | Low-fibre |
| Aldo/keto reductase | B2UN09 | 4.29 | 0.011623 | Low-fibre |
| Aldo/keto reductase\|g__Akkermansia.s__Akkermansia_muciniphila | B2UN09 | 4.29 | 0.011623 | Low-fibre |
| Bifunctional protein HldE | B2UNE1 | 4.46 | 0.011623 | Low-fibre |
| Bifunctional protein HldE\|g__Akkermansia.s__Akkermansia_muciniphila | B2UNE1 | 4.46 | 0.011623 | Low-fibre |
| Adenylosuccinate synthetase | B2UPQ6 | 3.96 | 0.011623 | Low-fibre |
| Adenylosuccinate synthetase\|g__Akkermansia.s__Akkermansia_muciniphila | B2UPQ6 | 3.96 | 0.011623 | Low-fibre |
| Helicase ATP-dependent domain protein | B2UM09 | 4.38 | 0.011623 | Low-fibre |
| Helicase ATP-dependent domain protein\|g__Akkermansia.s__Akkermansia_muciniphila | B2UM09 | 4.38 | 0.011623 | Low-fibre |
| Excinuclease ABC C subunit domain protein | B2URH7 | 4.17 | 0.011623 | Low-fibre |
| Excinuclease ABC C subunit domain protein\|g__Akkermansia.s__Akkermansia_muciniphila | B2URH7 | 4.17 | 0.011623 | Low-fibre |
| Major facilitator superfamily MFS_1 | B2ULI6 | 3.82 | 0.011623 | Low-fibre |
| Major facilitator superfamily MFS_1\|g__Akkermansia.s__Akkermansia_muciniphila | B2ULI6 | 3.82 | 0.011623 | Low-fibre |
| Inosine-5'-monophosphate dehydrogenase | B2UQ38 | 4.32 | 0.011629 | Low-fibre |
| Inosine-5'-monophosphate dehydrogenase\|g__Akkermansia.s__Akkermansia_muciniphila | B2UQ38 | 4.32 | 0.011629 | Low-fibre |
| Glycosyl transferase group 1 | B2UPJ8 | 3.90 | 0.011629 | Low-fibre |
| Glycosyl transferase group 1\|g__Akkermansia.s__Akkermansia_muciniphila | B2UPJ8 | 3.90 | 0.011629 | Low-fibre |
| Three-deoxy-D-manno-octulosonic-acid transferase domain protein | B2UNU2 | 4.18 | 0.011942 | Low-fibre |
| Three-deoxy-D-manno-octulosonic-acid transferase domain protein\|g__Akkermansia.s__Akkermansia_muciniphila | B2UNU2 | 4.18 | 0.011942 | Low-fibre |
| Chloride channel core | R7DW73 | 3.99 | 0.011942 | Low-fibre |
| Chloride channel core\|g__Akkermansia.s__Akkermansia_muciniphila | R7DW73 | 3.99 | 0.011942 | Low-fibre |
| NO_NAME | B2UQ63 | 3.79 | 0.011942 | Low-fibre |
| NO_NAME\|g__Akkermansia.s__Akkermansia_muciniphila | B2UQ63 | 3.79 | 0.011942 | Low-fibre |
| Efflux transporter, RND family, MFP subunit | B2ULQ9 | 4.17 | 0.011942 | Low-fibre |
| Efflux transporter, RND family, MFP subunit\|g__Akkermansia.s__Akkermansia_muciniphila | B2ULQ9 | 4.17 | 0.011942 | Low-fibre |
| Sodium ion-translocating decarboxylase, beta subunit | B2UM99 | 4.57 | 0.011942 | Low-fibre |
| Sodium ion-translocating decarboxylase, beta subunit\|g__Akkermansia.s__Akkermansia_muciniphila | B2UM99 | 4.57 | 0.011942 | Low-fibre |
| Non-specific serine/threonine protein kinase | B2UMN1 | 4.06 | 0.011942 | Low-fibre |
| Non-specific serine/threonine protein kinase\|g__Akkermansia.s__Akkermansia_muciniphila | B2UMN1 | 4.06 | 0.011942 | Low-fibre |
| Peptidoglycan glycosyltransferase | R6J1N9 | 4.43 | 0.011942 | Low-fibre |
| Peptidoglycan glycosyltransferase\|g__Akkermansia.s__Akkermansia_muciniphila | R6J1N9 | 4.43 | 0.011942 | Low-fibre |
| Uncharacterized conserved protein\|g__Bacteroides.s__Bacteroides_xylanisolvens | D6D3K7 | -4.24 | 0.012632 | High-fibre |
| NADH-quinone oxidoreductase subunit D | B2ULZ0 | 4.30 | 0.012632 | Low-fibre |
| NADH-quinone oxidoreductase subunit D\|g__Akkermansia.s__Akkermansia_muciniphila | B2ULZ0 | 4.30 | 0.012632 | Low-fibre |
| Polynucleotide adenylyltransferase/metal dependent phosphohydrolase | B2UM12 | 4.41 | 0.012898 | Low-fibre |
| Polynucleotide adenylyltransferase/metal dependent phosphohydrolase\|g__Akkermansia.s__Akkermansia_muciniphila | B2UM12 | 4.41 | 0.012898 | Low-fibre |
| DNA mismatch repair protein MutL | B2ULE2 | 3.73 | 0.012898 | Low-fibre |
| DNA mismatch repair protein MutL\|g__Akkermansia.s__Akkermansia_muciniphila | B2ULE2 | 3.73 | 0.012898 | Low-fibre |
| ATP-dependent nuclease subunit B-like protein | R6J3K8 | 4.31 | 0.013035 | Low-fibre |
| ATP-dependent nuclease subunit B-like protein\|g__Akkermansia.s__Akkermansia_muciniphila | R6J3K8 | 4.31 | 0.013035 | Low-fibre |
| NO_NAME | B2UR86 | 4.48 | 0.013035 | Low-fibre |
| NO_NAME\|g__Akkermansia.s__Akkermansia_muciniphila | B2UR86 | 4.48 | 0.013035 | Low-fibre |
| NO_NAME | B2UMZ2 | 3.90 | 0.013035 | Low-fibre |
| NO_NAME\|g__Akkermansia.s__Akkermansia_muciniphila | B2UMZ2 | 3.90 | 0.013035 | Low-fibre |
| Putative lipoprotein | B2UP41 | 3.90 | 0.013035 | Low-fibre |
| Putative lipoprotein\|g__Akkermansia.s__Akkermansia_muciniphila | B2UP41 | 3.90 | 0.013035 | Low-fibre |
| Phosphofructokinase | B2UM45 | 4.18 | 0.013035 | Low-fibre |
| Phosphofructokinase\|g__Akkermansia.s__Akkermansia_muciniphila | B2UM45 | 4.18 | 0.013035 | Low-fibre |
| NO_NAME | I8YQF4 | -4.20 | 0.013451 | High-fibre |
| ABC transporter related | B2ULV7 | 4.57 | 0.01364 | Low-fibre |
| ABC transporter related\|g__Akkermansia.s__Akkermansia_muciniphila | B2ULV7 | 4.57 | 0.01364 | Low-fibre |
| Chaperone protein DnaJ | B2UL09 | 4.14 | 0.01364 | Low-fibre |
| Chaperone protein DnaJ\|g__Akkermansia.s__Akkermansia_muciniphila | B2UL09 | 4.14 | 0.01364 | Low-fibre |
| Radical SAM domain protein | B2UQ90 | 3.83 | 0.01364 | Low-fibre |
| Radical SAM domain protein\|g__Akkermansia.s__Akkermansia_muciniphila | B2UQ90 | 3.83 | 0.01364 | Low-fibre |
| Histidine--tRNA ligase | B2UKU8 | 4.21 | 0.014767 | Low-fibre |
| Histidine--tRNA ligase\|g__Akkermansia.s__Akkermansia_muciniphila | B2UKU8 | 4.21 | 0.014767 | Low-fibre |
| NO_NAME | R6IWH8 | 4.14 | 0.015499 | Low-fibre |
| NO_NAME\|g__Akkermansia.s__Akkermansia_muciniphila | R6IWH8 | 4.14 | 0.015499 | Low-fibre |
| Asparagine--tRNA ligase | R6J148 | 4.15 | 0.016303 | Low-fibre |
| Asparagine--tRNA ligase\|g__Akkermansia.s__Akkermansia_muciniphila | R6J148 | 4.15 | 0.016303 | Low-fibre |
| Transporter, hydrophobe/amphiphile efflux-1 (HAE1) family | B2UP81 | 4.44 | 0.016303 | Low-fibre |
| Transporter, hydrophobe/amphiphile efflux-1 (HAE1) family\|g__Akkermansia.s__Akkermansia_muciniphila | B2UP81 | 4.44 | 0.016303 | Low-fibre |
| Methylmalonyl-CoA mutase | B2UNS9 | 5.37 | 0.016776 | Low-fibre |
| Methylmalonyl-CoA mutase\|g__Akkermansia.s__Akkermansia_muciniphila | B2UNS9 | 5.37 | 0.016776 | Low-fibre |
| NO_NAME | B2UPL7 | 3.71 | 0.016851 | Low-fibre |
| NO_NAME\|g__Akkermansia.s__Akkermansia_muciniphila | B2UPL7 | 3.71 | 0.016851 | Low-fibre |
| Ankyrin | B2UNE4 | 4.13 | 0.017029 | Low-fibre |
| Ankyrin\|g__Akkermansia.s__Akkermansia_muciniphila | B2UNE4 | 4.13 | 0.017029 | Low-fibre |
| GTP-binding protein TypA | B2UQ24 | 4.38 | 0.017035 | Low-fibre |
| GTP-binding protein TypA\|g__Akkermansia.s__Akkermansia_muciniphila | B2UQ24 | 4.38 | 0.017035 | Low-fibre |
| NO_NAME | B2ULL2 | 3.43 | 0.017509 | Low-fibre |
| NO_NAME\|g__Akkermansia.s__Akkermansia_muciniphila | B2ULL2 | 3.43 | 0.017509 | Low-fibre |
| ABC transporter related | B2ULW0 | 3.42 | 0.017509 | Low-fibre |
| ABC transporter related\|g__Akkermansia.s__Akkermansia_muciniphila | B2ULW0 | 3.42 | 0.017509 | Low-fibre |
| Exo-alpha-sialidase | R7E0G8 | 4.38 | 0.017509 | Low-fibre |
| Exo-alpha-sialidase\|g__Akkermansia.s__Akkermansia_muciniphila | R7E0G8 | 4.38 | 0.017509 | Low-fibre |
| Putative PTS IIA-like nitrogen-regulatory protein PtsN | B2UP87 | 3.59 | 0.017509 | Low-fibre |
| Putative PTS IIA-like nitrogen-regulatory protein PtsN\|g__Akkermansia.s__Akkermansia_muciniphila | B2UP87 | 3.59 | 0.017509 | Low-fibre |
| Thiazole synthase | R6IWR9 | 3.69 | 0.017509 | Low-fibre |
| Thiazole synthase\|g__Akkermansia.s__Akkermansia_muciniphila | R6IWR9 | 3.69 | 0.017509 | Low-fibre |
| NO_NAME | R6K002 | 3.80 | 0.017509 | Low-fibre |
| NO_NAME\|g__Akkermansia.s__Akkermansia_muciniphila | R6K002 | 3.80 | 0.017509 | Low-fibre |
| NO_NAME | B2ULT6 | 5.35 | 0.017509 | Low-fibre |
| NO_NAME\|g__Akkermansia.s__Akkermansia_muciniphila | B2ULT6 | 5.35 | 0.017509 | Low-fibre |
| Ribosomal L11 methyltransferase | B2ULZ5 | 3.53 | 0.017945 | Low-fibre |
| Ribosomal L11 methyltransferase\|g__Akkermansia.s__Akkermansia_muciniphila | B2ULZ5 | 3.53 | 0.017945 | Low-fibre |
| Beta-N-acetylhexosaminidase | R6J9B1 | 3.95 | 0.017945 | Low-fibre |
| Beta-N-acetylhexosaminidase\|g__Akkermansia.s__Akkermansia_muciniphila | R6J9B1 | 3.95 | 0.017945 | Low-fibre |
| Major facilitator superfamily MFS_1 | B2UKX3 | 4.14 | 0.017945 | Low-fibre |
| Major facilitator superfamily MFS_1\|g__Akkermansia.s__Akkermansia_muciniphila | B2UKX3 | 4.14 | 0.017945 | Low-fibre |
| NO_NAME | C6IQ45 | -4.06 | 0.018325 | High-fibre |
| NO_NAME | B2ULX2 | 5.23 | 0.018325 | Low-fibre |
| NO_NAME\|g__Akkermansia.s__Akkermansia_muciniphila | B2ULX2 | 5.23 | 0.018325 | Low-fibre |
| NO_NAME | N9ZZP4 | 4.36 | 0.018584 | Low-fibre |
| DNA topoisomerase III | F0R8X1 | 2.85 | 0.019214 | Low-fibre |
| Aldehyde Dehydrogenase | B2UPA5 | 5.13 | 0.019214 | Low-fibre |
| Aldehyde Dehydrogenase\|g__Akkermansia.s__Akkermansia_muciniphila | B2UPA5 | 5.13 | 0.019214 | Low-fibre |
| Beta-N-acetylhexosaminidase | B2UQG6 | 4.98 | 0.019214 | Low-fibre |
| Beta-N-acetylhexosaminidase\|g__Akkermansia.s__Akkermansia_muciniphila | B2UQG6 | 4.98 | 0.019214 | Low-fibre |
| NO_NAME | B2UL64 | 3.80 | 0.019262 | Low-fibre |
| NO_NAME\|g__Akkermansia.s__Akkermansia_muciniphila | B2UL64 | 3.80 | 0.019262 | Low-fibre |
| Lipolytic protein G-D-S-L family | B2URJ9 | 3.73 | 0.019262 | Low-fibre |
| Lipolytic protein G-D-S-L family\|g__Akkermansia.s__Akkermansia_muciniphila | B2URJ9 | 3.73 | 0.019262 | Low-fibre |
| von Willebrand factor type A | R6JWH1 | 4.96 | 0.019406 | Low-fibre |
| von Willebrand factor type A\|g__Akkermansia.s__Akkermansia_muciniphila | R6JWH1 | 4.96 | 0.019406 | Low-fibre |
| NO_NAME | D4WH80 | -4.76 | 0.019501 | High-fibre |
| NO_NAME\|g__Bacteroides.s__Bacteroides_ovatus | D4WH80 | -4.76 | 0.019501 | High-fibre |
| Uncharacterized conserved protein | D6D3K7 | -3.91 | 0.019501 | High-fibre |
| Succinate dehydrogenase (Or fumarate reductase) cytochrome b subunit, b558 family | B2UQT0 | 3.28 | 0.019501 | Low-fibre |
| Succinate dehydrogenase (Or fumarate reductase) cytochrome b subunit, b558 family\|g__Akkermansia.s__Akkermansia_muciniphila | B2UQT0 | 3.28 | 0.019501 | Low-fibre |
| H(+)-transporting two-sector ATPase | B2UQA4 | 3.98 | 0.019501 | Low-fibre |
| H(+)-transporting two-sector ATPase\|g__Akkermansia.s__Akkermansia_muciniphila | B2UQA4 | 3.98 | 0.019501 | Low-fibre |
| Glycoside hydrolase family 95 | B2UM81 | 3.95 | 0.019501 | Low-fibre |
| Glycoside hydrolase family 95\|g__Akkermansia.s__Akkermansia_muciniphila | B2UM81 | 3.95 | 0.019501 | Low-fibre |
| Chorismate mutase | B2UL67 | 4.09 | 0.019501 | Low-fibre |
| Chorismate mutase\|g__Akkermansia.s__Akkermansia_muciniphila | B2UL67 | 4.09 | 0.019501 | Low-fibre |
| NO_NAME | B2UR77 | 3.50 | 0.019501 | Low-fibre |
| NO_NAME\|g__Akkermansia.s__Akkermansia_muciniphila | B2UR77 | 3.50 | 0.019501 | Low-fibre |
| NO_NAME | B3C522 | 3.28 | 0.019501 | Low-fibre |
| Glycoside hydrolase family 2 TIM barrel | B2UM40 | 5.19 | 0.019501 | Low-fibre |
| Glycoside hydrolase family 2 TIM barrel\|g__Akkermansia.s__Akkermansia_muciniphila | B2UM40 | 5.19 | 0.019501 | Low-fibre |
| Acriflavin resistance protein | B2UR00 | 5.08 | 0.019501 | Low-fibre |
| Acriflavin resistance protein\|g__Akkermansia.s__Akkermansia_muciniphila | B2UR00 | 5.08 | 0.019501 | Low-fibre |
| Protein translocase subunit SecY | B2UQN0 | 5.04 | 0.019501 | Low-fibre |
| Protein translocase subunit SecY\|g__Akkermansia.s__Akkermansia_muciniphila | B2UQN0 | 5.04 | 0.019501 | Low-fibre |
| Transporter, hydrophobe/amphiphile efflux-1 (HAE1) family | B2UQI8 | 5.32 | 0.019501 | Low-fibre |
| Transporter, hydrophobe/amphiphile efflux-1 (HAE1) family\|g__Akkermansia.s__Akkermansia_muciniphila | B2UQI8 | 5.32 | 0.019501 | Low-fibre |
| Transcription termination factor Rho | B2UQ10 | 4.88 | 0.019501 | Low-fibre |
| Transcription termination factor Rho\|g__Akkermansia.s__Akkermansia_muciniphila | B2UQ10 | 4.88 | 0.019501 | Low-fibre |
| Aconitate hydratase 1 | R7E343 | 5.03 | 0.019501 | Low-fibre |
| Aconitate hydratase 1\|g__Akkermansia.s__Akkermansia_muciniphila | R7E343 | 5.03 | 0.019501 | Low-fibre |
| 60 kDa chaperonin | B2UKV8 | 5.16 | 0.019501 | Low-fibre |
| 60 kDa chaperonin\|g__Akkermansia.s__Akkermansia_muciniphila | B2UKV8 | 5.16 | 0.019501 | Low-fibre |
| Methylmalonyl-CoA mutase, large subunit | B2UNS8 | 5.30 | 0.019501 | Low-fibre |
| Methylmalonyl-CoA mutase, large subunit\|g__Akkermansia.s__Akkermansia_muciniphila | B2UNS8 | 5.30 | 0.019501 | Low-fibre |
| NO_NAME | B2UMU9 | 4.87 | 0.019501 | Low-fibre |
| NO_NAME\|g__Akkermansia.s__Akkermansia_muciniphila | B2UMU9 | 4.87 | 0.019501 | Low-fibre |
| NO_NAME | B2UNV8 | 5.09 | 0.020202 | Low-fibre |
| NO_NAME\|g__Akkermansia.s__Akkermansia_muciniphila | B2UNV8 | 5.09 | 0.020202 | Low-fibre |
| Thiamine pyrophosphate protein TPP binding domain protein | B2UMA8 | 4.89 | 0.020202 | Low-fibre |
| Thiamine pyrophosphate protein TPP binding domain protein\|g__Akkermansia.s__Akkermansia_muciniphila | B2UMA8 | 4.89 | 0.020202 | Low-fibre |
| Anion transporter | B2UMB5 | 4.98 | 0.020732 | Low-fibre |
| Anion transporter\|g__Akkermansia.s__Akkermansia_muciniphila | B2UMB5 | 4.98 | 0.020732 | Low-fibre |
| NO_NAME | R7E3W7 | 4.85 | 0.021002 | Low-fibre |
| NO_NAME\|g__Akkermansia.s__Akkermansia_muciniphila | R7E3W7 | 4.85 | 0.021002 | Low-fibre |
| Glycoside hydrolase family 18 | B2UPU3 | 5.08 | 0.021002 | Low-fibre |
| Glycoside hydrolase family 18\|g__Akkermansia.s__Akkermansia_muciniphila | B2UPU3 | 5.08 | 0.021002 | Low-fibre |
| NO_NAME | B2UL84 | 4.71 | 0.021056 | Low-fibre |
| NO_NAME\|g__Akkermansia.s__Akkermansia_muciniphila | B2UL84 | 4.71 | 0.021056 | Low-fibre |
| AAA domain protein | A0A015VDF0 | -3.33 | 0.022209 | High-fibre |
| Amino acid permease-associated region | R6JXK6 | 4.73 | 0.022645 | Low-fibre |
| Amino acid permease-associated region\|g__Akkermansia.s__Akkermansia_muciniphila | R6JXK6 | 4.73 | 0.022645 | Low-fibre |
| tRNA-specific 2-thiouridylase MnmA | B2UPK4 | 3.91 | 0.022801 | Low-fibre |
| tRNA-specific 2-thiouridylase MnmA\|g__Akkermansia.s__Akkermansia_muciniphila | B2UPK4 | 3.91 | 0.022801 | Low-fibre |
| Dihydroxy-acid dehydratase | B2UR52 | 4.80 | 0.022801 | Low-fibre |
| Dihydroxy-acid dehydratase\|g__Akkermansia.s__Akkermansia_muciniphila | B2UR52 | 4.80 | 0.022801 | Low-fibre |
| Tetratricopeptide TPR_2 repeat protein | R6JX86 | 4.76 | 0.022965 | Low-fibre |
| Tetratricopeptide TPR_2 repeat protein\|g__Akkermansia.s__Akkermansia_muciniphila | R6JX86 | 4.76 | 0.022965 | Low-fibre |
| NO_NAME | B2UQ69 | 4.95 | 0.02304 | Low-fibre |
| NO_NAME\|g__Akkermansia.s__Akkermansia_muciniphila | B2UQ69 | 4.95 | 0.02304 | Low-fibre |
| Molybdopterin oxidoreductase, iron-sulfur binding subunit | B2URF4 | 4.77 | 0.02304 | Low-fibre |
| Molybdopterin oxidoreductase, iron-sulfur binding subunit\|g__Akkermansia.s__Akkermansia_muciniphila | B2URF4 | 4.77 | 0.02304 | Low-fibre |
| Elongation factor G | B2UMU2 | 5.19 | 0.02304 | Low-fibre |
| Elongation factor G\|g__Akkermansia.s__Akkermansia_muciniphila | B2UMU2 | 5.19 | 0.02304 | Low-fibre |
| Ferredoxin | R6JB51 | 5.07 | 0.02304 | Low-fibre |
| Ferredoxin\|g__Akkermansia.s__Akkermansia_muciniphila | R6JB51 | 5.07 | 0.02304 | Low-fibre |
| NO_NAME | B2UMZ7 | 4.74 | 0.02304 | Low-fibre |
| NO_NAME\|g__Akkermansia.s__Akkermansia_muciniphila | B2UMZ7 | 4.74 | 0.02304 | Low-fibre |
| DNA primase | G2T2N3 | 2.48 | 0.023392 | Low-fibre |
| Formate--tetrahydrofolate ligase | B2UN63 | 4.79 | 0.023392 | Low-fibre |
| Formate--tetrahydrofolate ligase\|g__Akkermansia.s__Akkermansia_muciniphila | B2UN63 | 4.79 | 0.023392 | Low-fibre |
| NO_NAME | B2UQT6 | 4.82 | 0.023628 | Low-fibre |
| NO_NAME\|g__Akkermansia.s__Akkermansia_muciniphila | B2UQT6 | 4.82 | 0.023628 | Low-fibre |
| NO_NAME | R6VTJ6 | 2.68 | 0.023747 | Low-fibre |
| NO_NAME\|g__Bacteroides.s__Bacteroides_ovatus | R6VTJ6 | 2.68 | 0.023747 | Low-fibre |
| Cytochrome c assembly protein | B2UPA9 | 4.74 | 0.023747 | Low-fibre |
| Cytochrome c assembly protein\|g__Akkermansia.s__Akkermansia_muciniphila | B2UPA9 | 4.74 | 0.023747 | Low-fibre |
| tRNA uridine 5-carboxymethylaminomethyl modification enzyme MnmG | B2UQB0 | 5.02 | 0.023747 | Low-fibre |
| tRNA uridine 5-carboxymethylaminomethyl modification enzyme MnmG\|g__Akkermansia.s__Akkermansia_muciniphila | B2UQB0 | 5.02 | 0.023747 | Low-fibre |
| Glutamine synthetase catalytic region | B2URJ2 | 4.71 | 0.023747 | Low-fibre |
| Glutamine synthetase catalytic region\|g__Akkermansia.s__Akkermansia_muciniphila | B2URJ2 | 4.71 | 0.023747 | Low-fibre |
| DNA polymerase IV | B2UKN1 | 4.78 | 0.023803 | Low-fibre |
| DNA polymerase IV\|g__Akkermansia.s__Akkermansia_muciniphila | B2UKN1 | 4.78 | 0.023803 | Low-fibre |
| UDP-N-acetylglucosamine--N-acetylmuramyl-(pentapeptide) pyrophosphoryl-undecaprenol N-acetylglucosamine transferase | B2UPW1 | 4.73 | 0.023849 | Low-fibre |
| UDP-N-acetylglucosamine--N-acetylmuramyl-(pentapeptide) pyrophosphoryl-undecaprenol N-acetylglucosamine transferase\|g__Akkermansia.s__Akkermansia_muciniphila | B2UPW1 | 4.73 | 0.023849 | Low-fibre |
| NO_NAME\|g__Bacteroides.s__Bacteroides_ovatus | R5PHV4 | -2.58 | 0.023853 | High-fibre |
| 5-oxopent-3-ene-1,2,5-tricarboxylate decarboxylase | B2ULC0 | 3.07 | 0.023853 | Low-fibre |
| 5-oxopent-3-ene-1,2,5-tricarboxylate decarboxylase\|g__Akkermansia.s__Akkermansia_muciniphila | B2ULC0 | 3.07 | 0.023853 | Low-fibre |
| Bifunctional purine biosynthesis protein PurH | B2URE6 | 3.71 | 0.023853 | Low-fibre |
| Bifunctional purine biosynthesis protein PurH\|g__Akkermansia.s__Akkermansia_muciniphila | B2URE6 | 3.71 | 0.023853 | Low-fibre |
| NO_NAME\|g__Bacteroides.s__Bacteroides_ovatus | R6VKY9 | 2.57 | 0.023853 | Low-fibre |
| RND efflux system, outer membrane lipoprotein, NodT family | B2ULQ6 | 3.26 | 0.023853 | Low-fibre |
| RND efflux system, outer membrane lipoprotein, NodT family\|g__Akkermansia.s__Akkermansia_muciniphila | B2ULQ6 | 3.26 | 0.023853 | Low-fibre |
| Heavy metal translocating P-type ATPase | B2UQH4 | 4.77 | 0.023853 | Low-fibre |
| Heavy metal translocating P-type ATPase\|g__Akkermansia.s__Akkermansia_muciniphila | B2UQH4 | 4.77 | 0.023853 | Low-fibre |
| Sugar-phosphate isomerase, RpiB/LacA/LacB family | B2UM06 | 4.89 | 0.023853 | Low-fibre |
| Sugar-phosphate isomerase, RpiB/LacA/LacB family\|g__Akkermansia.s__Akkermansia_muciniphila | B2UM06 | 4.89 | 0.023853 | Low-fibre |
| Toprim domain family protein | C9KXZ1 | 2.70 | 0.023853 | Low-fibre |
| Membrane protein-like protein | B2UNQ4 | 4.81 | 0.023853 | Low-fibre |
| Membrane protein-like protein\|g__Akkermansia.s__Akkermansia_muciniphila | B2UNQ4 | 4.81 | 0.023853 | Low-fibre |
| Putative phosphate/sulphate permease | B2UKW0 | 4.64 | 0.023853 | Low-fibre |
| Putative phosphate/sulphate permease\|g__Akkermansia.s__Akkermansia_muciniphila | B2UKW0 | 4.64 | 0.023853 | Low-fibre |
| Glycosyl transferase family 11 | R6JEE0 | 4.56 | 0.023853 | Low-fibre |
| Glycosyl transferase family 11\|g__Akkermansia.s__Akkermansia_muciniphila | R6JEE0 | 4.56 | 0.023853 | Low-fibre |
| ATP-dependent DNA helicase, RecQ family | B2UR16 | 4.74 | 0.023853 | Low-fibre |
| ATP-dependent DNA helicase, RecQ family\|g__Akkermansia.s__Akkermansia_muciniphila | B2UR16 | 4.74 | 0.023853 | Low-fibre |
| CinA-like protein | B2UNF6 | 4.75 | 0.023853 | Low-fibre |
| CinA-like protein\|g__Akkermansia.s__Akkermansia_muciniphila | B2UNF6 | 4.75 | 0.023853 | Low-fibre |
| NO_NAME | B2ULI1 | 4.72 | 0.023853 | Low-fibre |
| NO_NAME\|g__Akkermansia.s__Akkermansia_muciniphila | B2ULI1 | 4.72 | 0.023853 | Low-fibre |
| NO_NAME | R6J0H2 | 4.47 | 0.023853 | Low-fibre |
| NO_NAME\|g__Akkermansia.s__Akkermansia_muciniphila | R6J0H2 | 4.47 | 0.023853 | Low-fibre |
| NO_NAME | B2URA5 | 4.59 | 0.023853 | Low-fibre |
| NO_NAME\|g__Akkermansia.s__Akkermansia_muciniphila | B2URA5 | 4.59 | 0.023853 | Low-fibre |
| Glycoside hydrolase family 2 sugar binding | B2UM41 | 4.65 | 0.023853 | Low-fibre |
| Glycoside hydrolase family 2 sugar binding\|g__Akkermansia.s__Akkermansia_muciniphila | B2UM41 | 4.65 | 0.023853 | Low-fibre |
| Sulfatase | B2UNH7 | 4.52 | 0.023853 | Low-fibre |
| Sulfatase\|g__Akkermansia.s__Akkermansia_muciniphila | B2UNH7 | 4.52 | 0.023853 | Low-fibre |
| Two component regulator propeller domain protein | B2UN98 | 4.78 | 0.023853 | Low-fibre |
| Two component regulator propeller domain protein\|g__Akkermansia.s__Akkermansia_muciniphila | B2UN98 | 4.78 | 0.023853 | Low-fibre |
| Alpha-glucan phosphorylase | B2UMC9 | 5.01 | 0.023853 | Low-fibre |
| Alpha-glucan phosphorylase\|g__Akkermansia.s__Akkermansia_muciniphila | B2UMC9 | 5.01 | 0.023853 | Low-fibre |
| ABC transporter related | R6IYB4 | 4.87 | 0.023853 | Low-fibre |
| ABC transporter related\|g__Akkermansia.s__Akkermansia_muciniphila | R6IYB4 | 4.87 | 0.023853 | Low-fibre |
| UDP-3-O-acylglucosamine N-acyltransferase | B2UND2 | 4.57 | 0.023853 | Low-fibre |
| UDP-3-O-acylglucosamine N-acyltransferase\|g__Akkermansia.s__Akkermansia_muciniphila | B2UND2 | 4.57 | 0.023853 | Low-fibre |
| Serine--tRNA ligase | B2UPP6 | 4.63 | 0.023853 | Low-fibre |
| Serine--tRNA ligase\|g__Akkermansia.s__Akkermansia_muciniphila | B2UPP6 | 4.63 | 0.023853 | Low-fibre |
| Outer membrane autotransporter barrel domain protein | B2ULZ6 | 4.54 | 0.023853 | Low-fibre |
| Outer membrane autotransporter barrel domain protein\|g__Akkermansia.s__Akkermansia_muciniphila | B2ULZ6 | 4.54 | 0.023853 | Low-fibre |
| Beta-N-acetylhexosaminidase | B2UP57 | 4.83 | 0.023853 | Low-fibre |
| Beta-N-acetylhexosaminidase\|g__Akkermansia.s__Akkermansia_muciniphila | B2UP57 | 4.83 | 0.023853 | Low-fibre |
| Cytochrome-c peroxidase | B2UR32 | 4.76 | 0.023853 | Low-fibre |
| Cytochrome-c peroxidase\|g__Akkermansia.s__Akkermansia_muciniphila | B2UR32 | 4.76 | 0.023853 | Low-fibre |
| Peptidase M42 family protein | R6JAH8 | 4.65 | 0.023853 | Low-fibre |
| Peptidase M42 family protein\|g__Akkermansia.s__Akkermansia_muciniphila | R6JAH8 | 4.65 | 0.023853 | Low-fibre |
| Tetratricopeptide TPR_2 repeat protein | B2UL83 | 4.54 | 0.023853 | Low-fibre |
| Tetratricopeptide TPR_2 repeat protein\|g__Akkermansia.s__Akkermansia_muciniphila | B2UL83 | 4.54 | 0.023853 | Low-fibre |
| L-fuculokinase | B2UN37 | 4.60 | 0.023853 | Low-fibre |
| L-fuculokinase\|g__Akkermansia.s__Akkermansia_muciniphila | B2UN37 | 4.60 | 0.023853 | Low-fibre |
| Capsular exopolysaccharide family | B2UPB6 | 4.54 | 0.023853 | Low-fibre |
| Capsular exopolysaccharide family\|g__Akkermansia.s__Akkermansia_muciniphila | B2UPB6 | 4.54 | 0.023853 | Low-fibre |
| Trypsin-like protein serine protease typically periplasmic contain C-terminal PDZ domain-like protein | B2UPX2 | 4.45 | 0.023853 | Low-fibre |
| Trypsin-like protein serine protease typically periplasmic contain C-terminal PDZ domain-like protein\|g__Akkermansia.s__Akkermansia_muciniphila | B2UPX2 | 4.45 | 0.023853 | Low-fibre |
| NO_NAME | R7E580 | 4.86 | 0.023853 | Low-fibre |
| NO_NAME\|g__Akkermansia.s__Akkermansia_muciniphila | R7E580 | 4.86 | 0.023853 | Low-fibre |
| Glycosyl transferase family 2 | B2UQ55 | 4.67 | 0.023853 | Low-fibre |
| Glycosyl transferase family 2\|g__Akkermansia.s__Akkermansia_muciniphila | B2UQ55 | 4.67 | 0.023853 | Low-fibre |
| NO_NAME | B2UNM2 | 4.43 | 0.023853 | Low-fibre |
| NO_NAME\|g__Akkermansia.s__Akkermansia_muciniphila | B2UNM2 | 4.43 | 0.023853 | Low-fibre |
| Glycosyl hydrolase family 98 putative carbohydrate binding module | B2UKY7 | 4.47 | 0.023853 | Low-fibre |
| Glycosyl hydrolase family 98 putative carbohydrate binding module\|g__Akkermansia.s__Akkermansia_muciniphila | B2UKY7 | 4.47 | 0.023853 | Low-fibre |
| NO_NAME | R7E588 | 4.54 | 0.023853 | Low-fibre |
| NO_NAME\|g__Akkermansia.s__Akkermansia_muciniphila | R7E588 | 4.54 | 0.023853 | Low-fibre |
| Outer membrane protein assembly complex, YaeT protein | B2UQZ4 | 3.67 | 0.024098 | Low-fibre |
| Outer membrane protein assembly complex, YaeT protein\|g__Akkermansia.s__Akkermansia_muciniphila | B2UQZ4 | 3.67 | 0.024098 | Low-fibre |
| Polypeptide-transport-associated domain protein FtsQ-type | B2UPW4 | 4.51 | 0.024174 | Low-fibre |
| Polypeptide-transport-associated domain protein FtsQ-type\|g__Akkermansia.s__Akkermansia_muciniphila | B2UPW4 | 4.51 | 0.024174 | Low-fibre |
| Lipid A biosynthesis acyltransferase | R6JEQ5 | 4.46 | 0.024174 | Low-fibre |
| Lipid A biosynthesis acyltransferase\|g__Akkermansia.s__Akkermansia_muciniphila | R6JEQ5 | 4.46 | 0.024174 | Low-fibre |
| NO_NAME | B2UR81 | 3.96 | 0.024185 | Low-fibre |
| NO_NAME\|g__Akkermansia.s__Akkermansia_muciniphila | B2UR81 | 3.96 | 0.024185 | Low-fibre |
| NO_NAME | C5EF02 | 3.11 | 0.024185 | Low-fibre |
| NADH:flavin oxidoreductase/NADH oxidase | R6J5P5 | 3.47 | 0.024185 | Low-fibre |
| NADH:flavin oxidoreductase/NADH oxidase\|g__Akkermansia.s__Akkermansia_muciniphila | R6J5P5 | 3.47 | 0.024185 | Low-fibre |
| NO_NAME | UPI00046A43F1 | 2.99 | 0.024185 | Low-fibre |
| Beta-N-acetylhexosaminidase | B2UPS8 | 4.41 | 0.024185 | Low-fibre |
| Beta-N-acetylhexosaminidase\|g__Akkermansia.s__Akkermansia_muciniphila | B2UPS8 | 4.41 | 0.024185 | Low-fibre |
| Phosphoglycerate mutase 1 family | B2UQ72 | 4.53 | 0.024185 | Low-fibre |
| Phosphoglycerate mutase 1 family\|g__Akkermansia.s__Akkermansia_muciniphila | B2UQ72 | 4.53 | 0.024185 | Low-fibre |
| Metal dependent phosphohydrolase | B2UQN4 | 4.54 | 0.024185 | Low-fibre |
| Metal dependent phosphohydrolase\|g__Akkermansia.s__Akkermansia_muciniphila | B2UQN4 | 4.54 | 0.024185 | Low-fibre |
| YD repeat protein | B2UR84 | 4.56 | 0.024185 | Low-fibre |
| YD repeat protein\|g__Akkermansia.s__Akkermansia_muciniphila | B2UR84 | 4.56 | 0.024185 | Low-fibre |
| NO_NAME | B2ULN2 | 4.38 | 0.024185 | Low-fibre |
| NO_NAME\|g__Akkermansia.s__Akkermansia_muciniphila | B2ULN2 | 4.38 | 0.024185 | Low-fibre |
| DEAD/DEAH box helicase domain protein | B2URF2 | 4.67 | 0.024185 | Low-fibre |
| DEAD/DEAH box helicase domain protein\|g__Akkermansia.s__Akkermansia_muciniphila | B2URF2 | 4.67 | 0.024185 | Low-fibre |
| NO_NAME | R6JA57 | 4.46 | 0.024185 | Low-fibre |
| NO_NAME\|g__Akkermansia.s__Akkermansia_muciniphila | R6JA57 | 4.46 | 0.024185 | Low-fibre |
| Beta-galactosidase | B2UQ71 | 4.54 | 0.024185 | Low-fibre |
| Beta-galactosidase\|g__Akkermansia.s__Akkermansia_muciniphila | B2UQ71 | 4.54 | 0.024185 | Low-fibre |
| UDP-N-acetylmuramoylalanine--D-glutamate ligase | B2UPV8 | 4.53 | 0.024185 | Low-fibre |
| UDP-N-acetylmuramoylalanine--D-glutamate ligase\|g__Akkermansia.s__Akkermansia_muciniphila | B2UPV8 | 4.53 | 0.024185 | Low-fibre |
| NO_NAME | R6J5D7 | 4.68 | 0.024185 | Low-fibre |
| NO_NAME\|g__Akkermansia.s__Akkermansia_muciniphila | R6J5D7 | 4.68 | 0.024185 | Low-fibre |
| Prolipoprotein diacylglyceryl transferase | B2UR45 | 4.48 | 0.024185 | Low-fibre |
| Prolipoprotein diacylglyceryl transferase\|g__Akkermansia.s__Akkermansia_muciniphila | B2UR45 | 4.48 | 0.024185 | Low-fibre |
| NO_NAME\|g__Bacteroides.s__Bacteroides_ovatus | I8YQF4 | -3.80 | 0.024204 | High-fibre |
| NO_NAME | W0EUY5 | -4.47 | 0.024204 | High-fibre |
| Putative PAS/PAC sensor protein | B2ULF0 | 3.21 | 0.024204 | Low-fibre |
| Putative PAS/PAC sensor protein\|g__Akkermansia.s__Akkermansia_muciniphila | B2ULF0 | 3.21 | 0.024204 | Low-fibre |
| Glycosyl transferase family 8 | B2UQ54 | 3.46 | 0.024204 | Low-fibre |
| Glycosyl transferase family 8\|g__Akkermansia.s__Akkermansia_muciniphila | B2UQ54 | 3.46 | 0.024204 | Low-fibre |
| NO_NAME | K5YMU9 | 4.06 | 0.024204 | Low-fibre |
| NO_NAME | R7E5M4 | 3.05 | 0.024204 | Low-fibre |
| NO_NAME\|g__Akkermansia.s__Akkermansia_muciniphila | R7E5M4 | 3.05 | 0.024204 | Low-fibre |
| DNA primase | UPI00016C66A6 | 2.31 | 0.024204 | Low-fibre |
| Elongation factor 4 | B2UQX7 | 4.48 | 0.024204 | Low-fibre |
| Elongation factor 4\|g__Akkermansia.s__Akkermansia_muciniphila | B2UQX7 | 4.48 | 0.024204 | Low-fibre |
| MscS Mechanosensitive ion channel | R7E058 | 4.40 | 0.024204 | Low-fibre |
| MscS Mechanosensitive ion channel\|g__Akkermansia.s__Akkermansia_muciniphila | R7E058 | 4.40 | 0.024204 | Low-fibre |
| Conserved hypothetical membrane protein | B2ULT5 | 4.50 | 0.024204 | Low-fibre |
| Conserved hypothetical membrane protein\|g__Akkermansia.s__Akkermansia_muciniphila | B2ULT5 | 4.50 | 0.024204 | Low-fibre |
| Uroporphyrin-III C-methyltransferase | R6JD31 | 4.51 | 0.024204 | Low-fibre |
| Uroporphyrin-III C-methyltransferase\|g__Akkermansia.s__Akkermansia_muciniphila | R6JD31 | 4.51 | 0.024204 | Low-fibre |
| NO_NAME | B2UPI7 | 4.42 | 0.024204 | Low-fibre |
| NO_NAME\|g__Akkermansia.s__Akkermansia_muciniphila | B2UPI7 | 4.42 | 0.024204 | Low-fibre |
| Lipoyl synthase | B2UR33 | 4.55 | 0.024204 | Low-fibre |
| Lipoyl synthase\|g__Akkermansia.s__Akkermansia_muciniphila | B2UR33 | 4.55 | 0.024204 | Low-fibre |
| NO_NAME | B2UM15 | 4.52 | 0.024204 | Low-fibre |
| NO_NAME\|g__Akkermansia.s__Akkermansia_muciniphila | B2UM15 | 4.52 | 0.024204 | Low-fibre |
| Glutamate dehydrogenase | B2UP90 | 4.44 | 0.024204 | Low-fibre |
| Glutamate dehydrogenase\|g__Akkermansia.s__Akkermansia_muciniphila | B2UP90 | 4.44 | 0.024204 | Low-fibre |
| Tetratricopeptide TPR_2 repeat protein | B2UPU7 | 4.67 | 0.024204 | Low-fibre |
| Tetratricopeptide TPR_2 repeat protein\|g__Akkermansia.s__Akkermansia_muciniphila | B2UPU7 | 4.67 | 0.024204 | Low-fibre |
| ABC transporter related | B2UQW8 | 4.43 | 0.024204 | Low-fibre |
| ABC transporter related\|g__Akkermansia.s__Akkermansia_muciniphila | B2UQW8 | 4.43 | 0.024204 | Low-fibre |
| Isocitrate dehydrogenase [NADP] | B2UN83 | 4.62 | 0.024204 | Low-fibre |
| Isocitrate dehydrogenase [NADP]\|g__Akkermansia.s__Akkermansia_muciniphila | B2UN83 | 4.62 | 0.024204 | Low-fibre |
| DNA polymerase I | B2UNN5 | 4.55 | 0.024204 | Low-fibre |
| DNA polymerase I\|g__Akkermansia.s__Akkermansia_muciniphila | B2UNN5 | 4.55 | 0.024204 | Low-fibre |
| NO_NAME | B2UNE6 | 4.38 | 0.024204 | Low-fibre |
| NO_NAME\|g__Akkermansia.s__Akkermansia_muciniphila | B2UNE6 | 4.38 | 0.024204 | Low-fibre |
| Sodium/hydrogen exchanger | B2UQ13 | 4.51 | 0.024204 | Low-fibre |
| Sodium/hydrogen exchanger\|g__Akkermansia.s__Akkermansia_muciniphila | B2UQ13 | 4.51 | 0.024204 | Low-fibre |
| Glycoside hydrolase family 2 sugar binding | R6JCV4 | 4.77 | 0.024204 | Low-fibre |
| Glycoside hydrolase family 2 sugar binding\|g__Akkermansia.s__Akkermansia_muciniphila | R6JCV4 | 4.77 | 0.024204 | Low-fibre |
| Pyridoxal-5'-phosphate-dependent protein beta subunit | B2UPE5 | 4.49 | 0.024204 | Low-fibre |
| Pyridoxal-5'-phosphate-dependent protein beta subunit\|g__Akkermansia.s__Akkermansia_muciniphila | B2UPE5 | 4.49 | 0.024204 | Low-fibre |
| NO_NAME | B2UKY3 | 4.56 | 0.024204 | Low-fibre |
| NO_NAME\|g__Akkermansia.s__Akkermansia_muciniphila | B2UKY3 | 4.56 | 0.024204 | Low-fibre |
| NO_NAME | B2UQJ7 | 4.51 | 0.024204 | Low-fibre |
| NO_NAME\|g__Akkermansia.s__Akkermansia_muciniphila | B2UQJ7 | 4.51 | 0.024204 | Low-fibre |
| Putative transmembrane protein | B2UNX0 | 4.73 | 0.024204 | Low-fibre |
| Putative transmembrane protein\|g__Akkermansia.s__Akkermansia_muciniphila | B2UNX0 | 4.73 | 0.024204 | Low-fibre |
| dTDP-glucose 4,6-dehydratase | B2UMA4 | 4.51 | 0.024204 | Low-fibre |
| dTDP-glucose 4,6-dehydratase\|g__Akkermansia.s__Akkermansia_muciniphila | B2UMA4 | 4.51 | 0.024204 | Low-fibre |
| Membrane protein insertase YidC | R7DZB1 | 4.41 | 0.024204 | Low-fibre |
| Membrane protein insertase YidC\|g__Akkermansia.s__Akkermansia_muciniphila | R7DZB1 | 4.41 | 0.024204 | Low-fibre |
| Sulfatase | B2UR15 | 4.33 | 0.024204 | Low-fibre |
| Sulfatase\|g__Akkermansia.s__Akkermansia_muciniphila | B2UR15 | 4.33 | 0.024204 | Low-fibre |
| Aminotransferase class I and II | B2UL36 | 4.59 | 0.024204 | Low-fibre |
| Aminotransferase class I and II\|g__Akkermansia.s__Akkermansia_muciniphila | B2UL36 | 4.59 | 0.024204 | Low-fibre |
| Anthranilate phosphoribosyltransferase | B2ULG6 | 4.33 | 0.024204 | Low-fibre |
| Anthranilate phosphoribosyltransferase\|g__Akkermansia.s__Akkermansia_muciniphila | B2ULG6 | 4.33 | 0.024204 | Low-fibre |
| Ketol-acid reductoisomerase | B2URB8 | 4.59 | 0.024204 | Low-fibre |
| Ketol-acid reductoisomerase\|g__Akkermansia.s__Akkermansia_muciniphila | B2URB8 | 4.59 | 0.024204 | Low-fibre |
| NO_NAME | R7E0M5 | 4.59 | 0.024204 | Low-fibre |
| NO_NAME\|g__Akkermansia.s__Akkermansia_muciniphila | R7E0M5 | 4.59 | 0.024204 | Low-fibre |
| Alpha-L-fucosidase | B2UL68 | 4.36 | 0.024204 | Low-fibre |
| Alpha-L-fucosidase\|g__Akkermansia.s__Akkermansia_muciniphila | B2UL68 | 4.36 | 0.024204 | Low-fibre |
| Alkyl sulfatase and related hydrolase-like protein | B2UP61 | 4.66 | 0.024204 | Low-fibre |
| Alkyl sulfatase and related hydrolase-like protein\|g__Akkermansia.s__Akkermansia_muciniphila | B2UP61 | 4.66 | 0.024204 | Low-fibre |
| NO_NAME | B2UL46 | 4.25 | 0.024204 | Low-fibre |
| NO_NAME\|g__Akkermansia.s__Akkermansia_muciniphila | B2UL46 | 4.25 | 0.024204 | Low-fibre |
| NO_NAME | B2UNC7 | 4.23 | 0.024204 | Low-fibre |
| NO_NAME\|g__Akkermansia.s__Akkermansia_muciniphila | B2UNC7 | 4.23 | 0.024204 | Low-fibre |
| Ornithine carbamoyltransferase | B2UML6 | 4.24 | 0.024204 | Low-fibre |
| Ornithine carbamoyltransferase\|g__Akkermansia.s__Akkermansia_muciniphila | B2UML6 | 4.24 | 0.024204 | Low-fibre |
| L-fucose transporter | B2UN40 | 4.33 | 0.024204 | Low-fibre |
| L-fucose transporter\|g__Akkermansia.s__Akkermansia_muciniphila | B2UN40 | 4.33 | 0.024204 | Low-fibre |
| Sel1 domain protein repeat-containing protein | B2UNS3 | 4.30 | 0.024204 | Low-fibre |
| Sel1 domain protein repeat-containing protein\|g__Akkermansia.s__Akkermansia_muciniphila | B2UNS3 | 4.30 | 0.024204 | Low-fibre |
| NO_NAME | B2UQL6 | 4.29 | 0.024404 | Low-fibre |
| NO_NAME\|g__Akkermansia.s__Akkermansia_muciniphila | B2UQL6 | 4.29 | 0.024404 | Low-fibre |
| 4Fe-4S ferredoxin iron-sulfur binding domain protein | B2UQQ9 | 4.43 | 0.024404 | Low-fibre |
| 4Fe-4S ferredoxin iron-sulfur binding domain protein\|g__Akkermansia.s__Akkermansia_muciniphila | B2UQQ9 | 4.43 | 0.024404 | Low-fibre |
| NO_NAME | R6VKY9 | 2.41 | 0.024474 | Low-fibre |
| NO_NAME | B2UMS8 | 4.19 | 0.024474 | Low-fibre |
| NO_NAME\|g__Akkermansia.s__Akkermansia_muciniphila | B2UMS8 | 4.19 | 0.024474 | Low-fibre |
| Cell division protein FtsZ | B2ULV2 | 4.20 | 0.024474 | Low-fibre |
| Cell division protein FtsZ\|g__Akkermansia.s__Akkermansia_muciniphila | B2ULV2 | 4.20 | 0.024474 | Low-fibre |
| Citrate transporter | B2UQ66 | 4.49 | 0.024474 | Low-fibre |
| Citrate transporter\|g__Akkermansia.s__Akkermansia_muciniphila | B2UQ66 | 4.49 | 0.024474 | Low-fibre |
| Efflux transporter, RND family, MFP subunit | B2UQI7 | 4.29 | 0.024474 | Low-fibre |
| Efflux transporter, RND family, MFP subunit\|g__Akkermansia.s__Akkermansia_muciniphila | B2UQI7 | 4.29 | 0.024474 | Low-fibre |
| NO_NAME | B2UQD5 | 4.25 | 0.024474 | Low-fibre |
| NO_NAME\|g__Akkermansia.s__Akkermansia_muciniphila | B2UQD5 | 4.25 | 0.024474 | Low-fibre |
| Oxygen-independent coproporphyrinogen III oxidase | B2UPP4 | 4.42 | 0.024835 | Low-fibre |
| Oxygen-independent coproporphyrinogen III oxidase\|g__Akkermansia.s__Akkermansia_muciniphila | B2UPP4 | 4.42 | 0.024835 | Low-fibre |
| Lipolytic protein G-D-S-L family | B2ULI0 | 4.36 | 0.024835 | Low-fibre |
| Lipolytic protein G-D-S-L family\|g__Akkermansia.s__Akkermansia_muciniphila | B2ULI0 | 4.36 | 0.024835 | Low-fibre |
| Cytidyltransferase-related domain protein | B2UMK9 | 4.38 | 0.024835 | Low-fibre |
| Cytidyltransferase-related domain protein\|g__Akkermansia.s__Akkermansia_muciniphila | B2UMK9 | 4.38 | 0.024835 | Low-fibre |
| Capsular exopolysaccharide family | B2UKW3 | 4.51 | 0.024835 | Low-fibre |
| Capsular exopolysaccharide family\|g__Akkermansia.s__Akkermansia_muciniphila | B2UKW3 | 4.51 | 0.024835 | Low-fibre |
| Glycosyl transferase family 2 | B2UNW7 | 4.51 | 0.024835 | Low-fibre |
| Glycosyl transferase family 2\|g__Akkermansia.s__Akkermansia_muciniphila | B2UNW7 | 4.51 | 0.024835 | Low-fibre |
| RteR protein | Q8G9D9 | -2.64 | 0.024903 | High-fibre |
| ABC transporter related | B2UND4 | 3.58 | 0.024903 | Low-fibre |
| ABC transporter related\|g__Akkermansia.s__Akkermansia_muciniphila | B2UND4 | 3.58 | 0.024903 | Low-fibre |
| TPR repeat-containing protein | B2UMV2 | 4.23 | 0.024903 | Low-fibre |
| TPR repeat-containing protein\|g__Akkermansia.s__Akkermansia_muciniphila | B2UMV2 | 4.23 | 0.024903 | Low-fibre |
| NO_NAME | B2UNA7 | 4.36 | 0.024903 | Low-fibre |
| NO_NAME\|g__Akkermansia.s__Akkermansia_muciniphila | B2UNA7 | 4.36 | 0.024903 | Low-fibre |
| NO_NAME | B2UQE1 | 4.60 | 0.024903 | Low-fibre |
| NO_NAME\|g__Akkermansia.s__Akkermansia_muciniphila | B2UQE1 | 4.60 | 0.024903 | Low-fibre |
| Coagulation factor 5/8 type domain protein | B2UNB9 | 4.41 | 0.024903 | Low-fibre |
| Coagulation factor 5/8 type domain protein\|g__Akkermansia.s__Akkermansia_muciniphila | B2UNB9 | 4.41 | 0.024903 | Low-fibre |
| NO_NAME | B2UPZ2 | 4.16 | 0.024903 | Low-fibre |
| NO_NAME\|g__Akkermansia.s__Akkermansia_muciniphila | B2UPZ2 | 4.16 | 0.024903 | Low-fibre |
| NO_NAME | B2UN04 | 4.37 | 0.024903 | Low-fibre |
| NO_NAME\|g__Akkermansia.s__Akkermansia_muciniphila | B2UN04 | 4.37 | 0.024903 | Low-fibre |
| DEAD/DEAH box helicase domain protein | B2UNH8 | 4.51 | 0.024903 | Low-fibre |
| DEAD/DEAH box helicase domain protein\|g__Akkermansia.s__Akkermansia_muciniphila | B2UNH8 | 4.51 | 0.024903 | Low-fibre |
| NO_NAME | B2UMH9 | 4.24 | 0.024903 | Low-fibre |
| NO_NAME\|g__Akkermansia.s__Akkermansia_muciniphila | B2UMH9 | 4.24 | 0.024903 | Low-fibre |
| Serine/threonine protein kinase | B2UR99 | 4.59 | 0.024903 | Low-fibre |
| Serine/threonine protein kinase\|g__Akkermansia.s__Akkermansia_muciniphila | B2UR99 | 4.59 | 0.024903 | Low-fibre |
| NO_NAME | B2UQK5 | 4.11 | 0.0256 | Low-fibre |
| NO_NAME\|g__Akkermansia.s__Akkermansia_muciniphila | B2UQK5 | 4.11 | 0.0256 | Low-fibre |
| NO_NAME | B2ULB1 | 3.00 | 0.025778 | Low-fibre |
| Aldose 1-epimerase | B2URK1 | 4.32 | 0.025961 | Low-fibre |
| Aldose 1-epimerase\|g__Akkermansia.s__Akkermansia_muciniphila | B2URK1 | 4.32 | 0.025961 | Low-fibre |
| NO_NAME | R9IH41 | 3.08 | 0.02611 | Low-fibre |
| Cell cycle protein | B2UPW0 | 3.85 | 0.02611 | Low-fibre |
| Cell cycle protein\|g__Akkermansia.s__Akkermansia_muciniphila | B2UPW0 | 3.85 | 0.02611 | Low-fibre |
| Succinate CoA transferase\|g__Akkermansia.s__Akkermansia_muciniphila | R6K3X5 | 4.36 | 0.02611 | Low-fibre |
| SSS sodium solute transporter superfamily | B2ULX1 | 4.13 | 0.02611 | Low-fibre |
| SSS sodium solute transporter superfamily\|g__Akkermansia.s__Akkermansia_muciniphila | B2ULX1 | 4.13 | 0.02611 | Low-fibre |
| Lysine--tRNA ligase | B2UQ97 | 4.29 | 0.02611 | Low-fibre |
| Lysine--tRNA ligase\|g__Akkermansia.s__Akkermansia_muciniphila | B2UQ97 | 4.29 | 0.02611 | Low-fibre |
| Metallophosphoesterase | B2ULU9 | 4.30 | 0.02611 | Low-fibre |
| Metallophosphoesterase\|g__Akkermansia.s__Akkermansia_muciniphila | B2ULU9 | 4.30 | 0.02611 | Low-fibre |
| Malate dehydrogenase | B2UKY5 | 4.50 | 0.02611 | Low-fibre |
| Malate dehydrogenase\|g__Akkermansia.s__Akkermansia_muciniphila | B2UKY5 | 4.50 | 0.02611 | Low-fibre |
| Peptidoglycan glycosyltransferase | B2UNY2 | 4.41 | 0.02611 | Low-fibre |
| Peptidoglycan glycosyltransferase\|g__Akkermansia.s__Akkermansia_muciniphila | B2UNY2 | 4.41 | 0.02611 | Low-fibre |
| NO_NAME | R7E4Q0 | 4.20 | 0.02611 | Low-fibre |
| NO_NAME\|g__Akkermansia.s__Akkermansia_muciniphila | R7E4Q0 | 4.20 | 0.02611 | Low-fibre |
| NO_NAME | B2UQN7 | 4.22 | 0.02611 | Low-fibre |
| NO_NAME\|g__Akkermansia.s__Akkermansia_muciniphila | B2UQN7 | 4.22 | 0.02611 | Low-fibre |
| NO_NAME | B2ULM6 | 4.38 | 0.02611 | Low-fibre |
| NO_NAME\|g__Akkermansia.s__Akkermansia_muciniphila | B2ULM6 | 4.38 | 0.02611 | Low-fibre |
| ErfK/YbiS/YcfS/YnhG family protein | B2UQB8 | 4.13 | 0.02611 | Low-fibre |
| ErfK/YbiS/YcfS/YnhG family protein\|g__Akkermansia.s__Akkermansia_muciniphila | B2UQB8 | 4.13 | 0.02611 | Low-fibre |
| Beta-N-acetylhexosaminidase | B2UM43 | 4.32 | 0.02611 | Low-fibre |
| Beta-N-acetylhexosaminidase\|g__Akkermansia.s__Akkermansia_muciniphila | B2UM43 | 4.32 | 0.02611 | Low-fibre |
| NO_NAME | B2UQS6 | 4.29 | 0.026123 | Low-fibre |
| NO_NAME\|g__Akkermansia.s__Akkermansia_muciniphila | B2UQS6 | 4.29 | 0.026123 | Low-fibre |
| UBA/THIF-type NAD/FAD binding protein | B2UNZ0 | 4.22 | 0.026125 | Low-fibre |
| UBA/THIF-type NAD/FAD binding protein\|g__Akkermansia.s__Akkermansia_muciniphila | B2UNZ0 | 4.22 | 0.026125 | Low-fibre |
| Glycosyl hydrolase family 88 | B2UQG1 | 4.32 | 0.026125 | Low-fibre |
| Glycosyl hydrolase family 88\|g__Akkermansia.s__Akkermansia_muciniphila | B2UQG1 | 4.32 | 0.026125 | Low-fibre |
| Aspartyl/glutamyl-tRNA(Asn/Gln) amidotransferase subunit B | B2UMA7 | 4.33 | 0.026191 | Low-fibre |
| Aspartyl/glutamyl-tRNA(Asn/Gln) amidotransferase subunit B\|g__Akkermansia.s__Akkermansia_muciniphila | B2UMA7 | 4.33 | 0.026191 | Low-fibre |
| NO_NAME | E1WW60 | 2.45 | 0.026196 | Low-fibre |
| Cation diffusion facilitator family transporter | B2UQ12 | 4.41 | 0.026196 | Low-fibre |
| Cation diffusion facilitator family transporter\|g__Akkermansia.s__Akkermansia_muciniphila | B2UQ12 | 4.41 | 0.026196 | Low-fibre |
| Excinuclease ABC, A subunit | B2UL01 | 4.65 | 0.026354 | Low-fibre |
| Excinuclease ABC, A subunit\|g__Akkermansia.s__Akkermansia_muciniphila | B2UL01 | 4.65 | 0.026354 | Low-fibre |
| Serine/threonine protein kinase | B2ULD6 | 4.27 | 0.026354 | Low-fibre |
| Serine/threonine protein kinase\|g__Akkermansia.s__Akkermansia_muciniphila | B2ULD6 | 4.27 | 0.026354 | Low-fibre |
| NUDIX hydrolase | B2UMR4 | 4.26 | 0.026354 | Low-fibre |
| NUDIX hydrolase\|g__Akkermansia.s__Akkermansia_muciniphila | B2UMR4 | 4.26 | 0.026354 | Low-fibre |
| NO_NAME | B2ULS0 | 4.18 | 0.026407 | Low-fibre |
| NO_NAME\|g__Akkermansia.s__Akkermansia_muciniphila | B2ULS0 | 4.18 | 0.026407 | Low-fibre |
| Outer membrane autotransporter barrel domain protein | B2UMI5 | 4.31 | 0.026407 | Low-fibre |
| Outer membrane autotransporter barrel domain protein\|g__Akkermansia.s__Akkermansia_muciniphila | B2UMI5 | 4.31 | 0.026407 | Low-fibre |
| DNA topoisomerase (ATP-hydrolyzing) | R7DUI7 | 4.16 | 0.026619 | Low-fibre |
| DNA topoisomerase (ATP-hydrolyzing)\|g__Akkermansia.s__Akkermansia_muciniphila | R7DUI7 | 4.16 | 0.026619 | Low-fibre |
| NO_NAME | B2UR64 | 4.13 | 0.026649 | Low-fibre |
| NO_NAME\|g__Akkermansia.s__Akkermansia_muciniphila | B2UR64 | 4.13 | 0.026649 | Low-fibre |
| Aminoglycoside phosphotransferase | B2UM73 | 4.06 | 0.026722 | Low-fibre |
| Aminoglycoside phosphotransferase\|g__Akkermansia.s__Akkermansia_muciniphila | B2UM73 | 4.06 | 0.026722 | Low-fibre |
| DNA polymerase III, subunits gamma and tau | B2URG8 | 4.24 | 0.026722 | Low-fibre |
| DNA polymerase III, subunits gamma and tau\|g__Akkermansia.s__Akkermansia_muciniphila | B2URG8 | 4.24 | 0.026722 | Low-fibre |
| Type II secretion system protein E | B2ULL8 | 4.33 | 0.026722 | Low-fibre |
| Type II secretion system protein E\|g__Akkermansia.s__Akkermansia_muciniphila | B2ULL8 | 4.33 | 0.026722 | Low-fibre |
| Alpha-galactosidase | R6J0Q6 | 4.19 | 0.026801 | Low-fibre |
| Alpha-galactosidase\|g__Akkermansia.s__Akkermansia_muciniphila | R6J0Q6 | 4.19 | 0.026801 | Low-fibre |
| Succinate dehydrogenase or fumarate reductase, flavoprotein subunit | B2UQS9 | 4.69 | 0.026801 | Low-fibre |
| Succinate dehydrogenase or fumarate reductase, flavoprotein subunit\|g__Akkermansia.s__Akkermansia_muciniphila | B2UQS9 | 4.69 | 0.026801 | Low-fibre |
| NO_NAME | Q8A5F3 | 2.91 | 0.026801 | Low-fibre |
| NO_NAME | B2UP33 | 4.44 | 0.026801 | Low-fibre |
| NO_NAME\|g__Akkermansia.s__Akkermansia_muciniphila | B2UP33 | 4.44 | 0.026801 | Low-fibre |
| GreA/GreB family elongation factor | R6IZ73 | 4.53 | 0.026801 | Low-fibre |
| GreA/GreB family elongation factor\|g__Akkermansia.s__Akkermansia_muciniphila | R6IZ73 | 4.53 | 0.026801 | Low-fibre |
| 4-hydroxy-3-methylbut-2-enyl diphosphate reductase | B2UM22 | 4.37 | 0.026801 | Low-fibre |
| 4-hydroxy-3-methylbut-2-enyl diphosphate reductase\|g__Akkermansia.s__Akkermansia_muciniphila | B2UM22 | 4.37 | 0.026801 | Low-fibre |
| Metallophosphoesterase | B2ULC8 | 4.19 | 0.026801 | Low-fibre |
| Metallophosphoesterase\|g__Akkermansia.s__Akkermansia_muciniphila | B2ULC8 | 4.19 | 0.026801 | Low-fibre |
| Argininosuccinate lyase | R6IZP5 | 4.26 | 0.026801 | Low-fibre |
| Argininosuccinate lyase\|g__Akkermansia.s__Akkermansia_muciniphila | R6IZP5 | 4.26 | 0.026801 | Low-fibre |
| Transport system permease protein | B2UNM8 | 4.47 | 0.026801 | Low-fibre |
| Transport system permease protein\|g__Akkermansia.s__Akkermansia_muciniphila | B2UNM8 | 4.47 | 0.026801 | Low-fibre |
| Integral membrane sensor signal transduction histidine kinase | B2UMI9 | 4.24 | 0.026898 | Low-fibre |
| Integral membrane sensor signal transduction histidine kinase\|g__Akkermansia.s__Akkermansia_muciniphila | B2UMI9 | 4.24 | 0.026898 | Low-fibre |
| NO_NAME | B2UPQ5 | 4.17 | 0.026958 | Low-fibre |
| NO_NAME\|g__Akkermansia.s__Akkermansia_muciniphila | B2UPQ5 | 4.17 | 0.026958 | Low-fibre |
| Dihydrolipoyl dehydrogenase | B2UM63 | 4.03 | 0.027167 | Low-fibre |
| Dihydrolipoyl dehydrogenase\|g__Akkermansia.s__Akkermansia_muciniphila | B2UM63 | 4.03 | 0.027167 | Low-fibre |
| NO_NAME | B2UL63 | 4.16 | 0.027171 | Low-fibre |
| NO_NAME\|g__Akkermansia.s__Akkermansia_muciniphila | B2UL63 | 4.16 | 0.027171 | Low-fibre |
| ATP-dependent Clp protease proteolytic subunit | B2UQZ2 | 4.01 | 0.027171 | Low-fibre |
| ATP-dependent Clp protease proteolytic subunit\|g__Akkermansia.s__Akkermansia_muciniphila | B2UQZ2 | 4.01 | 0.027171 | Low-fibre |
| Trigger factor | B2UQZ3 | 3.99 | 0.027171 | Low-fibre |
| Trigger factor\|g__Akkermansia.s__Akkermansia_muciniphila | B2UQZ3 | 3.99 | 0.027171 | Low-fibre |
| MATE efflux family protein | B2ULA6 | 4.05 | 0.027429 | Low-fibre |
| MATE efflux family protein\|g__Akkermansia.s__Akkermansia_muciniphila | B2ULA6 | 4.05 | 0.027429 | Low-fibre |
| NO_NAME | R5B0Q2 | 2.68 | 0.027491 | Low-fibre |
| Metal dependent phosphohydrolase | B2UQE5 | 4.59 | 0.02754 | Low-fibre |
| Metal dependent phosphohydrolase\|g__Akkermansia.s__Akkermansia_muciniphila | B2UQE5 | 4.59 | 0.02754 | Low-fibre |
| HPr kinase/phosphorylase | R7E120 | 4.35 | 0.027609 | Low-fibre |
| HPr kinase/phosphorylase\|g__Akkermansia.s__Akkermansia_muciniphila | R7E120 | 4.35 | 0.027609 | Low-fibre |
| NO_NAME\|g__Bacteroides.s__Bacteroides_ovatus | R6JCQ7 | -5.13 | 0.027618 | High-fibre |
| GDP-L-fucose synthase | B2URI8 | 4.54 | 0.027724 | Low-fibre |
| GDP-L-fucose synthase\|g__Akkermansia.s__Akkermansia_muciniphila | B2URI8 | 4.54 | 0.027724 | Low-fibre |
| Metallophosphoesterase | B2UQ53 | 4.43 | 0.027963 | Low-fibre |
| Metallophosphoesterase\|g__Akkermansia.s__Akkermansia_muciniphila | B2UQ53 | 4.43 | 0.027963 | Low-fibre |
| TDP-4-keto-6-deoxy-D-glucose transaminase | R6IZ32 | 4.67 | 0.028086 | Low-fibre |
| TDP-4-keto-6-deoxy-D-glucose transaminase\|g__Akkermansia.s__Akkermansia_muciniphila | R6IZ32 | 4.67 | 0.028086 | Low-fibre |
| TatD-related deoxyribonuclease | B2UQT2 | 4.30 | 0.028086 | Low-fibre |
| TatD-related deoxyribonuclease\|g__Akkermansia.s__Akkermansia_muciniphila | B2UQT2 | 4.30 | 0.028086 | Low-fibre |
| Heavy metal translocating P-type ATPase | R6IX25 | 4.61 | 0.028086 | Low-fibre |
| Heavy metal translocating P-type ATPase\|g__Akkermansia.s__Akkermansia_muciniphila | R6IX25 | 4.61 | 0.028086 | Low-fibre |
| TonB-dependent receptor | B2UM58 | 4.36 | 0.028184 | Low-fibre |
| TonB-dependent receptor\|g__Akkermansia.s__Akkermansia_muciniphila | B2UM58 | 4.36 | 0.028184 | Low-fibre |
| Ribonuclease 3 | R6IZB0 | 4.03 | 0.028184 | Low-fibre |
| Ribonuclease 3\|g__Akkermansia.s__Akkermansia_muciniphila | R6IZB0 | 4.03 | 0.028184 | Low-fibre |
| Fe2+-dicitrate sensor, membrane component | D6D782 | 3.50 | 0.028234 | Low-fibre |
| Polar amino acid ABC transporter, inner membrane subunit | B2UKZ5 | 4.22 | 0.028234 | Low-fibre |
| Polar amino acid ABC transporter, inner membrane subunit\|g__Akkermansia.s__Akkermansia_muciniphila | B2UKZ5 | 4.22 | 0.028234 | Low-fibre |
| Glutamate decarboxylase | R6IYN9 | 4.14 | 0.028245 | Low-fibre |
| Glutamate decarboxylase\|g__Akkermansia.s__Akkermansia_muciniphila | R6IYN9 | 4.14 | 0.028245 | Low-fibre |
| Bifunctional protein FolD | B2UM17 | 4.02 | 0.028245 | Low-fibre |
| Bifunctional protein FolD\|g__Akkermansia.s__Akkermansia_muciniphila | B2UM17 | 4.02 | 0.028245 | Low-fibre |
| Citrate synthase | B2ULK6 | 4.00 | 0.0283 | Low-fibre |
| Citrate synthase\|g__Akkermansia.s__Akkermansia_muciniphila | B2ULK6 | 4.00 | 0.0283 | Low-fibre |
| Ribonuclease Rne/Rng family | R6IZ82 | 4.28 | 0.028407 | Low-fibre |
| Ribonuclease Rne/Rng family\|g__Akkermansia.s__Akkermansia_muciniphila | R6IZ82 | 4.28 | 0.028407 | Low-fibre |
| Ribonuclease R | R6J1J8 | 4.03 | 0.028407 | Low-fibre |
| Ribonuclease R\|g__Akkermansia.s__Akkermansia_muciniphila | R6J1J8 | 4.03 | 0.028407 | Low-fibre |
| NO_NAME | B2UR61 | 4.03 | 0.028407 | Low-fibre |
| NO_NAME\|g__Akkermansia.s__Akkermansia_muciniphila | B2UR61 | 4.03 | 0.028407 | Low-fibre |
| NO_NAME | B2UKZ2 | 4.21 | 0.028438 | Low-fibre |
| NO_NAME\|g__Akkermansia.s__Akkermansia_muciniphila | B2UKZ2 | 4.21 | 0.028438 | Low-fibre |
| Maltose O-acetyltransferase | B2UNB1 | 3.38 | 0.028482 | Low-fibre |
| Maltose O-acetyltransferase\|g__Akkermansia.s__Akkermansia_muciniphila | B2UNB1 | 3.38 | 0.028482 | Low-fibre |
| Exopolysaccharide biosynthesis polyprenyl glycosylphosphotransferase | B2UQR5 | 4.17 | 0.028482 | Low-fibre |
| Exopolysaccharide biosynthesis polyprenyl glycosylphosphotransferase\|g__Akkermansia.s__Akkermansia_muciniphila | B2UQR5 | 4.17 | 0.028482 | Low-fibre |
| Cytochrome bd ubiquinol oxidase subunit I | B2UM69 | 4.09 | 0.028482 | Low-fibre |
| Cytochrome bd ubiquinol oxidase subunit I\|g__Akkermansia.s__Akkermansia_muciniphila | B2UM69 | 4.09 | 0.028482 | Low-fibre |
| Beta-N-acetylhexosaminidase | B2UP58 | 4.51 | 0.028482 | Low-fibre |
| Beta-N-acetylhexosaminidase\|g__Akkermansia.s__Akkermansia_muciniphila | B2UP58 | 4.51 | 0.028482 | Low-fibre |
| NO_NAME | B2URA9 | 4.12 | 0.028482 | Low-fibre |
| NO_NAME\|g__Akkermansia.s__Akkermansia_muciniphila | B2URA9 | 4.12 | 0.028482 | Low-fibre |
| NO_NAME | R9N1R7 | -2.85 | 0.028774 | High-fibre |
| Putative SAM-dependent methyltransferase | B2UN28 | 4.02 | 0.028774 | Low-fibre |
| Putative SAM-dependent methyltransferase\|g__Akkermansia.s__Akkermansia_muciniphila | B2UN28 | 4.02 | 0.028774 | Low-fibre |
| GTPase HflX | B2UPE7 | 4.25 | 0.028774 | Low-fibre |
| GTPase HflX\|g__Akkermansia.s__Akkermansia_muciniphila | B2UPE7 | 4.25 | 0.028774 | Low-fibre |
| Glycosyl transferase family 2 | B2UQ58 | 4.01 | 0.028908 | Low-fibre |
| Glycosyl transferase family 2\|g__Akkermansia.s__Akkermansia_muciniphila | B2UQ58 | 4.01 | 0.028908 | Low-fibre |
| NO_NAME | B2URL6 | 3.91 | 0.028969 | Low-fibre |
| NO_NAME\|g__Akkermansia.s__Akkermansia_muciniphila | B2URL6 | 3.91 | 0.028969 | Low-fibre |
| NO_NAME | B2UNP2 | 4.12 | 0.029082 | Low-fibre |
| NO_NAME\|g__Akkermansia.s__Akkermansia_muciniphila | B2UNP2 | 4.12 | 0.029082 | Low-fibre |
| Fructose-bisphosphate aldolase, class II | B2UQ23 | 4.13 | 0.029101 | Low-fibre |
| Fructose-bisphosphate aldolase, class II\|g__Akkermansia.s__Akkermansia_muciniphila | B2UQ23 | 4.13 | 0.029101 | Low-fibre |
| NO_NAME\|g__Bacteroides.s__Bacteroides_ovatus | K5YTB0 | -2.24 | 0.029339 | High-fibre |
| CDP-diacylglycerol/serine O-phosphatidyltransferase | B2UKT5 | 3.21 | 0.029345 | Low-fibre |
| CDP-diacylglycerol/serine O-phosphatidyltransferase\|g__Akkermansia.s__Akkermansia_muciniphila | B2UKT5 | 3.21 | 0.029345 | Low-fibre |
| O-acetylhomoserine/O-acetylserine sulfhydrylase | B2UP52 | 3.99 | 0.029371 | Low-fibre |
| O-acetylhomoserine/O-acetylserine sulfhydrylase\|g__Akkermansia.s__Akkermansia_muciniphila | B2UP52 | 3.99 | 0.029371 | Low-fibre |
| Single-stranded-DNA-specific exonuclease RecJ | B2ULA8 | 4.22 | 0.029371 | Low-fibre |
| Single-stranded-DNA-specific exonuclease RecJ\|g__Akkermansia.s__Akkermansia_muciniphila | B2ULA8 | 4.22 | 0.029371 | Low-fibre |
| Proline--tRNA ligase | B2UL92 | 4.13 | 0.029371 | Low-fibre |
| Proline--tRNA ligase\|g__Akkermansia.s__Akkermansia_muciniphila | B2UL92 | 4.13 | 0.029371 | Low-fibre |
| MATE efflux family protein | B2UR72 | 4.02 | 0.029371 | Low-fibre |
| MATE efflux family protein\|g__Akkermansia.s__Akkermansia_muciniphila | B2UR72 | 4.02 | 0.029371 | Low-fibre |
| NO_NAME | B2UNK7 | 3.89 | 0.029371 | Low-fibre |
| NO_NAME\|g__Akkermansia.s__Akkermansia_muciniphila | B2UNK7 | 3.89 | 0.029371 | Low-fibre |
| Putative ribosome biogenesis GTPase RsgA | B2UNU0 | 3.91 | 0.029371 | Low-fibre |
| Putative ribosome biogenesis GTPase RsgA\|g__Akkermansia.s__Akkermansia_muciniphila | B2UNU0 | 3.91 | 0.029371 | Low-fibre |
| Tryptophanase | B2UP35 | 4.05 | 0.02951 | Low-fibre |
| Tryptophanase\|g__Akkermansia.s__Akkermansia_muciniphila | B2UP35 | 4.05 | 0.02951 | Low-fibre |
| 3-oxoacyl-(Acyl-carrier-protein) reductase | R7DVZ8 | 4.17 | 0.029539 | Low-fibre |
| 3-oxoacyl-(Acyl-carrier-protein) reductase\|g__Akkermansia.s__Akkermansia_muciniphila | R7DVZ8 | 4.17 | 0.029539 | Low-fibre |
| NO_NAME | B2UNU1 | 4.03 | 0.029539 | Low-fibre |
| NO_NAME\|g__Akkermansia.s__Akkermansia_muciniphila | B2UNU1 | 4.03 | 0.029539 | Low-fibre |
| Chloride channel core | B2UP88 | 4.25 | 0.029594 | Low-fibre |
| Chloride channel core\|g__Akkermansia.s__Akkermansia_muciniphila | B2UP88 | 4.25 | 0.029594 | Low-fibre |
| NO_NAME | B2UP01 | 3.00 | 0.029836 | Low-fibre |
| NO_NAME\|g__Akkermansia.s__Akkermansia_muciniphila | B2UP01 | 3.00 | 0.029836 | Low-fibre |
| Riboflavin biosynthesis protein RibD | B2UQB1 | 3.85 | 0.029836 | Low-fibre |
| Riboflavin biosynthesis protein RibD\|g__Akkermansia.s__Akkermansia_muciniphila | B2UQB1 | 3.85 | 0.029836 | Low-fibre |
| Heavy metal translocating P-type ATPase | B2UN45 | 3.89 | 0.029945 | Low-fibre |
| Heavy metal translocating P-type ATPase\|g__Akkermansia.s__Akkermansia_muciniphila | B2UN45 | 3.89 | 0.029945 | Low-fibre |
| OmpA/MotB domain protein | B2UPY4 | 4.02 | 0.030038 | Low-fibre |
| OmpA/MotB domain protein\|g__Akkermansia.s__Akkermansia_muciniphila | B2UPY4 | 4.02 | 0.030038 | Low-fibre |
| FHA domain containing protein | B2UQ28 | 4.02 | 0.030038 | Low-fibre |
| FHA domain containing protein\|g__Akkermansia.s__Akkermansia_muciniphila | B2UQ28 | 4.02 | 0.030038 | Low-fibre |
| NO_NAME | R7DVV9 | 3.86 | 0.030038 | Low-fibre |
| NO_NAME\|g__Akkermansia.s__Akkermansia_muciniphila | R7DVV9 | 3.86 | 0.030038 | Low-fibre |
| NO_NAME | B2UPD4 | 3.98 | 0.030038 | Low-fibre |
| NO_NAME\|g__Akkermansia.s__Akkermansia_muciniphila | B2UPD4 | 3.98 | 0.030038 | Low-fibre |
| NO_NAME | B2UNA8 | 3.95 | 0.030038 | Low-fibre |
| NO_NAME\|g__Akkermansia.s__Akkermansia_muciniphila | B2UNA8 | 3.95 | 0.030038 | Low-fibre |
| CRISPR-associated endonuclease Cas1 | B2UP48 | 3.85 | 0.030038 | Low-fibre |
| CRISPR-associated endonuclease Cas1\|g__Akkermansia.s__Akkermansia_muciniphila | B2UP48 | 3.85 | 0.030038 | Low-fibre |
| Permease YjgP/YjgQ family protein | B2UMV4 | 3.87 | 0.030038 | Low-fibre |
| Permease YjgP/YjgQ family protein\|g__Akkermansia.s__Akkermansia_muciniphila | B2UMV4 | 3.87 | 0.030038 | Low-fibre |
| Sel1 domain protein repeat-containing protein | B2UPU2 | 4.12 | 0.030038 | Low-fibre |
| Sel1 domain protein repeat-containing protein\|g__Akkermansia.s__Akkermansia_muciniphila | B2UPU2 | 4.12 | 0.030038 | Low-fibre |
| NO_NAME | B2UM94 | 4.30 | 0.030049 | Low-fibre |
| NO_NAME\|g__Akkermansia.s__Akkermansia_muciniphila | B2UM94 | 4.30 | 0.030049 | Low-fibre |
| NO_NAME | B2UR95 | 4.44 | 0.030101 | Low-fibre |
| NO_NAME\|g__Akkermansia.s__Akkermansia_muciniphila | B2UR95 | 4.44 | 0.030101 | Low-fibre |
| NO_NAME | E1WW58 | 2.71 | 0.030148 | Low-fibre |
| NO_NAME | N2B7E9 | -4.93 | 0.030281 | High-fibre |
| Lipid-A-disaccharide synthase | B2UP91 | 3.97 | 0.030281 | Low-fibre |
| Lipid-A-disaccharide synthase\|g__Akkermansia.s__Akkermansia_muciniphila | B2UP91 | 3.97 | 0.030281 | Low-fibre |
| Biosynthetic arginine decarboxylase | B2UM33 | 4.24 | 0.030283 | Low-fibre |
| Biosynthetic arginine decarboxylase\|g__Akkermansia.s__Akkermansia_muciniphila | B2UM33 | 4.24 | 0.030283 | Low-fibre |
| NAD-dependent epimerase/dehydratase | B2UL87 | 4.15 | 0.030283 | Low-fibre |
| NAD-dependent epimerase/dehydratase\|g__Akkermansia.s__Akkermansia_muciniphila | B2UL87 | 4.15 | 0.030283 | Low-fibre |
| NO_NAME | B2UP04 | 3.84 | 0.03036 | Low-fibre |
| NO_NAME\|g__Akkermansia.s__Akkermansia_muciniphila | B2UP04 | 3.84 | 0.03036 | Low-fibre |
| Alanine racemase | B2UN53 | 4.10 | 0.030508 | Low-fibre |
| Alanine racemase\|g__Akkermansia.s__Akkermansia_muciniphila | B2UN53 | 4.10 | 0.030508 | Low-fibre |
| NO_NAME | B2UKX2 | 4.02 | 0.030603 | Low-fibre |
| NO_NAME\|g__Akkermansia.s__Akkermansia_muciniphila | B2UKX2 | 4.02 | 0.030603 | Low-fibre |
| Sulfite reductase (NADPH) | B2URN3 | 4.06 | 0.030623 | Low-fibre |
| Sulfite reductase (NADPH)\|g__Akkermansia.s__Akkermansia_muciniphila | B2URN3 | 4.06 | 0.030623 | Low-fibre |
| Integrase family protein | B2UPG0 | 4.02 | 0.031054 | Low-fibre |
| Integrase family protein\|g__Akkermansia.s__Akkermansia_muciniphila | B2UPG0 | 4.02 | 0.031054 | Low-fibre |
| Ctn044 | Q56VD5 | 2.21 | 0.031194 | Low-fibre |
| NO_NAME | R9HU68 | -2.33 | 0.031688 | High-fibre |
| NO_NAME | B2UQ21 | 3.45 | 0.031688 | Low-fibre |
| NO_NAME\|g__Akkermansia.s__Akkermansia_muciniphila | B2UQ21 | 3.45 | 0.031688 | Low-fibre |
| Signal recognition particle protein | R6J218 | 3.99 | 0.031688 | Low-fibre |
| Signal recognition particle protein\|g__Akkermansia.s__Akkermansia_muciniphila | R6J218 | 3.99 | 0.031688 | Low-fibre |
| YidE/YbjL duplication | B2UQF4 | 4.38 | 0.031688 | Low-fibre |
| YidE/YbjL duplication\|g__Akkermansia.s__Akkermansia_muciniphila | B2UQF4 | 4.38 | 0.031688 | Low-fibre |
| Outer membrane autotransporter barrel domain protein | B2UPY9 | 4.01 | 0.031688 | Low-fibre |
| Outer membrane autotransporter barrel domain protein\|g__Akkermansia.s__Akkermansia_muciniphila | B2UPY9 | 4.01 | 0.031688 | Low-fibre |
| Anthranilate synthase | R7DSS0 | 4.19 | 0.031688 | Low-fibre |
| Anthranilate synthase\|g__Akkermansia.s__Akkermansia_muciniphila | R7DSS0 | 4.19 | 0.031688 | Low-fibre |
| Methyltransferase type 11 | B2UMK4 | 4.02 | 0.031688 | Low-fibre |
| Methyltransferase type 11\|g__Akkermansia.s__Akkermansia_muciniphila | B2UMK4 | 4.02 | 0.031688 | Low-fibre |
| NO_NAME | B2ULS4 | 4.04 | 0.031688 | Low-fibre |
| NO_NAME\|g__Akkermansia.s__Akkermansia_muciniphila | B2ULS4 | 4.04 | 0.031688 | Low-fibre |
| Na+/H+ antiporter NhaC | B2URI7 | 4.15 | 0.031688 | Low-fibre |
| Na+/H+ antiporter NhaC\|g__Akkermansia.s__Akkermansia_muciniphila | B2URI7 | 4.15 | 0.031688 | Low-fibre |
| Aminotransferase class I and II | B2ULM2 | 3.92 | 0.031724 | Low-fibre |
| Aminotransferase class I and II\|g__Akkermansia.s__Akkermansia_muciniphila | B2ULM2 | 3.92 | 0.031724 | Low-fibre |
| NO_NAME | B2UNK6 | 3.84 | 0.031724 | Low-fibre |
| NO_NAME\|g__Akkermansia.s__Akkermansia_muciniphila | B2UNK6 | 3.84 | 0.031724 | Low-fibre |
| Haloacid dehalogenase domain protein hydrolase | B2UL11 | 3.83 | 0.031724 | Low-fibre |
| Haloacid dehalogenase domain protein hydrolase\|g__Akkermansia.s__Akkermansia_muciniphila | B2UL11 | 3.83 | 0.031724 | Low-fibre |
| UDP-N-acetylmuramate | B2UPY3 | 4.24 | 0.031803 | Low-fibre |
| UDP-N-acetylmuramate\|g__Akkermansia.s__Akkermansia_muciniphila | B2UPY3 | 4.24 | 0.031803 | Low-fibre |
| NO_NAME | B2UNB5 | 4.02 | 0.031903 | Low-fibre |
| NO_NAME\|g__Akkermansia.s__Akkermansia_muciniphila | B2UNB5 | 4.02 | 0.031903 | Low-fibre |
| Regulatory protein GntR HTH | B2UPI4 | 3.75 | 0.031903 | Low-fibre |
| Regulatory protein GntR HTH\|g__Akkermansia.s__Akkermansia_muciniphila | B2UPI4 | 3.75 | 0.031903 | Low-fibre |
| tRNA N6-adenosine threonylcarbamoyltransferase | B2UQZ0 | 3.93 | 0.031903 | Low-fibre |
| tRNA N6-adenosine threonylcarbamoyltransferase\|g__Akkermansia.s__Akkermansia_muciniphila | B2UQZ0 | 3.93 | 0.031903 | Low-fibre |
| NO_NAME | R7DYU5 | 4.14 | 0.031903 | Low-fibre |
| NO_NAME\|g__Akkermansia.s__Akkermansia_muciniphila | R7DYU5 | 4.14 | 0.031903 | Low-fibre |
| Glycosyl transferase family 51 | B2URI1 | 3.95 | 0.031903 | Low-fibre |
| Glycosyl transferase family 51\|g__Akkermansia.s__Akkermansia_muciniphila | B2URI1 | 3.95 | 0.031903 | Low-fibre |
| Aldo/keto reductase | B2UMN7 | 4.01 | 0.032051 | Low-fibre |
| Aldo/keto reductase\|g__Akkermansia.s__Akkermansia_muciniphila | B2UMN7 | 4.01 | 0.032051 | Low-fibre |
| Phosphoesterase RecJ domain protein | B2UQ51 | 4.39 | 0.032117 | Low-fibre |
| Phosphoesterase RecJ domain protein\|g__Akkermansia.s__Akkermansia_muciniphila | B2UQ51 | 4.39 | 0.032117 | Low-fibre |
| Phosphate import ATP-binding protein PstB | B2URP6 | 4.29 | 0.032229 | Low-fibre |
| Phosphate import ATP-binding protein PstB\|g__Akkermansia.s__Akkermansia_muciniphila | B2URP6 | 4.29 | 0.032229 | Low-fibre |
| Adenylosuccinate lyase | B2UNN8 | 4.01 | 0.032229 | Low-fibre |
| Adenylosuccinate lyase\|g__Akkermansia.s__Akkermansia_muciniphila | B2UNN8 | 4.01 | 0.032229 | Low-fibre |
| NO_NAME | B2UPK0 | 3.78 | 0.032229 | Low-fibre |
| NO_NAME\|g__Akkermansia.s__Akkermansia_muciniphila | B2UPK0 | 3.78 | 0.032229 | Low-fibre |
| Polysaccharide export protein | B2UKW4 | 4.04 | 0.032229 | Low-fibre |
| Polysaccharide export protein\|g__Akkermansia.s__Akkermansia_muciniphila | B2UKW4 | 4.04 | 0.032229 | Low-fibre |
| NO_NAME | R7E2D9 | -3.04 | 0.032399 | High-fibre |
| Potassium-transporting ATPase A chain | B2UR91 | 4.13 | 0.032758 | Low-fibre |
| Potassium-transporting ATPase A chain\|g__Akkermansia.s__Akkermansia_muciniphila | B2UR91 | 4.13 | 0.032758 | Low-fibre |
| Conserved protein found in conjugate transposon | Q8A5D7 | 2.32 | 0.032758 | Low-fibre |
| NO_NAME | B2UR13 | 4.08 | 0.032769 | Low-fibre |
| NO_NAME\|g__Akkermansia.s__Akkermansia_muciniphila | B2UR13 | 4.08 | 0.032769 | Low-fibre |
| Serine/threonine protein kinase | B2UKX1 | 3.76 | 0.032769 | Low-fibre |
| Serine/threonine protein kinase\|g__Akkermansia.s__Akkermansia_muciniphila | B2UKX1 | 3.76 | 0.032769 | Low-fibre |
| Alcohol dehydrogenase zinc-binding domain protein | B2UPP7 | 3.89 | 0.032787 | Low-fibre |
| Alcohol dehydrogenase zinc-binding domain protein\|g__Akkermansia.s__Akkermansia_muciniphila | B2UPP7 | 3.89 | 0.032787 | Low-fibre |
| Alanine--glyoxylate transaminase | B2UQ05 | 3.76 | 0.032787 | Low-fibre |
| Alanine--glyoxylate transaminase\|g__Akkermansia.s__Akkermansia_muciniphila | B2UQ05 | 3.76 | 0.032787 | Low-fibre |
| NO_NAME | B2UPK6 | 4.03 | 0.032787 | Low-fibre |
| NO_NAME\|g__Akkermansia.s__Akkermansia_muciniphila | B2UPK6 | 4.03 | 0.032787 | Low-fibre |
| Amino acid/peptide transporter | R7E3Y9 | 4.04 | 0.032787 | Low-fibre |
| Amino acid/peptide transporter\|g__Akkermansia.s__Akkermansia_muciniphila | R7E3Y9 | 4.04 | 0.032787 | Low-fibre |
| Malate dehydrogenase (Oxaloacetate-decarboxylating) (NADP(+)) | R7DTH7 | 3.99 | 0.032787 | Low-fibre |
| Malate dehydrogenase (Oxaloacetate-decarboxylating) (NADP(+))\|g__Akkermansia.s__Akkermansia_muciniphila | R7DTH7 | 3.99 | 0.032787 | Low-fibre |
| Nucleotidyl transferase | B2UNL6 | 3.04 | 0.032897 | Low-fibre |
| Nucleotidyl transferase\|g__Akkermansia.s__Akkermansia_muciniphila | B2UNL6 | 3.04 | 0.032897 | Low-fibre |
| Outer membrane autotransporter barrel domain protein | B2UQJ0 | 4.21 | 0.032959 | Low-fibre |
| Outer membrane autotransporter barrel domain protein\|g__Akkermansia.s__Akkermansia_muciniphila | B2UQJ0 | 4.21 | 0.032959 | Low-fibre |
| NO_NAME | C3QQ01 | -2.36 | 0.033111 | High-fibre |
| NO_NAME | B2UQZ5 | 3.13 | 0.033111 | Low-fibre |
| NO_NAME\|g__Akkermansia.s__Akkermansia_muciniphila | B2UQZ5 | 3.13 | 0.033111 | Low-fibre |
| Putative substrate-binding protein of aliphatic sulfonate ABC transporter | B2UND5 | 4.10 | 0.033111 | Low-fibre |
| Putative substrate-binding protein of aliphatic sulfonate ABC transporter\|g__Akkermansia.s__Akkermansia_muciniphila | B2UND5 | 4.10 | 0.033111 | Low-fibre |
| NADH dehydrogenase (Quinone) | B2ULY8 | 3.82 | 0.033111 | Low-fibre |
| NADH dehydrogenase (Quinone)\|g__Akkermansia.s__Akkermansia_muciniphila | B2ULY8 | 3.82 | 0.033111 | Low-fibre |
| GDP-mannose 4,6-dehydratase\|g__Akkermansia.s__Akkermansia_muciniphila | B2URI9 | 3.81 | 0.033111 | Low-fibre |
| Peptidase M50 | B2ULJ2 | 3.83 | 0.033111 | Low-fibre |
| Peptidase M50\|g__Akkermansia.s__Akkermansia_muciniphila | B2ULJ2 | 3.83 | 0.033111 | Low-fibre |
| Peptidase A24A domain protein | B2UL51 | 4.32 | 0.03321 | Low-fibre |
| Peptidase A24A domain protein\|g__Akkermansia.s__Akkermansia_muciniphila | B2UL51 | 4.32 | 0.03321 | Low-fibre |
| NO_NAME | B2ULT7 | 4.02 | 0.03321 | Low-fibre |
| NO_NAME\|g__Akkermansia.s__Akkermansia_muciniphila | B2ULT7 | 4.02 | 0.03321 | Low-fibre |
| AcrB/D/F family transporter | D4VN41 | 2.40 | 0.03321 | Low-fibre |
| NO_NAME\|g__Bacteroides.s__Bacteroides_ovatus | C6IQ45 | -3.49 | 0.033756 | High-fibre |
| NO_NAME | E2BKA3 | -4.71 | 0.033759 | High-fibre |
| ErfK/YbiS/YcfS/YnhG family protein | R7E4Y9 | 3.85 | 0.033759 | Low-fibre |
| ErfK/YbiS/YcfS/YnhG family protein\|g__Akkermansia.s__Akkermansia_muciniphila | R7E4Y9 | 3.85 | 0.033759 | Low-fibre |
| NO_NAME | B2URI6 | 3.77 | 0.033759 | Low-fibre |
| NO_NAME\|g__Akkermansia.s__Akkermansia_muciniphila | B2URI6 | 3.77 | 0.033759 | Low-fibre |
| NO_NAME | R5AXR7 | 2.78 | 0.03395 | Low-fibre |
| NO_NAME | C0B7Y7 | -2.42 | 0.034468 | High-fibre |
| Glycosyl transferase family 11 | B2UR82 | 2.91 | 0.034468 | Low-fibre |
| Glycosyl transferase family 11\|g__Akkermansia.s__Akkermansia_muciniphila | B2UR82 | 2.91 | 0.034468 | Low-fibre |
| NO_NAME\|g__Akkermansia.s__Akkermansia_muciniphila | B2UNK1 | 3.40 | 0.034559 | Low-fibre |
| NO_NAME | R9HU25 | -2.27 | 0.03469 | High-fibre |
| NO_NAME | R9HVW0 | 2.13 | 0.03469 | Low-fibre |
| NO_NAME\|g__Bacteroides.s__Bacteroides_ovatus | R6JIX5 | -4.67 | 0.034816 | High-fibre |
| Glucokinase\|g__Bacteroides.s__Bacteroides_ovatus | D6D2R8 | -4.66 | 0.034816 | High-fibre |
| Tetraacyldisaccharide 4'-kinase | B2UPD5 | 4.22 | 0.034816 | Low-fibre |
| Tetraacyldisaccharide 4'-kinase\|g__Akkermansia.s__Akkermansia_muciniphila | B2UPD5 | 4.22 | 0.034816 | Low-fibre |
| Glycosyl transferase family 2 | B2UQP2 | 3.98 | 0.034816 | Low-fibre |
| Glycosyl transferase family 2\|g__Akkermansia.s__Akkermansia_muciniphila | B2UQP2 | 3.98 | 0.034816 | Low-fibre |
| Phosphoribosylformylglycinamidine synthase 2 | B2UP02 | 4.07 | 0.034816 | Low-fibre |
| Phosphoribosylformylglycinamidine synthase 2\|g__Akkermansia.s__Akkermansia_muciniphila | B2UP02 | 4.07 | 0.034816 | Low-fibre |
| NO_NAME | B2UM61 | 3.03 | 0.034816 | Low-fibre |
| NO_NAME\|g__Akkermansia.s__Akkermansia_muciniphila | B2UM61 | 3.03 | 0.034816 | Low-fibre |
| Lipoprotein signal peptidase | B2UM85 | 3.94 | 0.034816 | Low-fibre |
| Lipoprotein signal peptidase\|g__Akkermansia.s__Akkermansia_muciniphila | B2UM85 | 3.94 | 0.034816 | Low-fibre |
| NO_NAME | B2UN26 | 3.69 | 0.034816 | Low-fibre |
| NO_NAME\|g__Akkermansia.s__Akkermansia_muciniphila | B2UN26 | 3.69 | 0.034816 | Low-fibre |
| Saccharopine dehydrogenase | B2ULU8 | 3.76 | 0.034816 | Low-fibre |
| Saccharopine dehydrogenase\|g__Akkermansia.s__Akkermansia_muciniphila | B2ULU8 | 3.76 | 0.034816 | Low-fibre |
| NO_NAME | B2UN67 | 3.65 | 0.034816 | Low-fibre |
| NO_NAME\|g__Akkermansia.s__Akkermansia_muciniphila | B2UN67 | 3.65 | 0.034816 | Low-fibre |
| PpiC-type peptidyl-prolyl cis-trans isomerase | B2ULW7 | 3.82 | 0.034816 | Low-fibre |
| PpiC-type peptidyl-prolyl cis-trans isomerase\|g__Akkermansia.s__Akkermansia_muciniphila | B2ULW7 | 3.82 | 0.034816 | Low-fibre |
| NO_NAME | B2UKX0 | 3.75 | 0.034816 | Low-fibre |
| NO_NAME\|g__Akkermansia.s__Akkermansia_muciniphila | B2UKX0 | 3.75 | 0.034816 | Low-fibre |
| NO_NAME | B2UPI2 | 3.88 | 0.034816 | Low-fibre |
| NO_NAME\|g__Akkermansia.s__Akkermansia_muciniphila | B2UPI2 | 3.88 | 0.034816 | Low-fibre |
| NO_NAME | R7E2M9 | 3.65 | 0.034816 | Low-fibre |
| NO_NAME\|g__Akkermansia.s__Akkermansia_muciniphila | R7E2M9 | 3.65 | 0.034816 | Low-fibre |
| Thiamine-phosphate synthase | B2UP54 | 3.76 | 0.034816 | Low-fibre |
| Thiamine-phosphate synthase\|g__Akkermansia.s__Akkermansia_muciniphila | B2UP54 | 3.76 | 0.034816 | Low-fibre |
| D-alanine--D-alanine ligase | B2UPW3 | 3.86 | 0.034816 | Low-fibre |
| D-alanine--D-alanine ligase\|g__Akkermansia.s__Akkermansia_muciniphila | B2UPW3 | 3.86 | 0.034816 | Low-fibre |
| Serine/threonine protein kinase | B2UM91 | 4.23 | 0.034816 | Low-fibre |
| Serine/threonine protein kinase\|g__Akkermansia.s__Akkermansia_muciniphila | B2UM91 | 4.23 | 0.034816 | Low-fibre |
| NO_NAME | B2URA7 | 3.35 | 0.034816 | Low-fibre |
| NO_NAME\|g__Akkermansia.s__Akkermansia_muciniphila | B2URA7 | 3.35 | 0.034816 | Low-fibre |
| Phospho-N-acetylmuramoyl-pentapeptide-transferase | B2UPL5 | 4.11 | 0.034816 | Low-fibre |
| Phospho-N-acetylmuramoyl-pentapeptide-transferase\|g__Akkermansia.s__Akkermansia_muciniphila | B2UPL5 | 4.11 | 0.034816 | Low-fibre |
| Beta-lactamase domain protein | B2UPK8 | 3.65 | 0.034816 | Low-fibre |
| Beta-lactamase domain protein\|g__Akkermansia.s__Akkermansia_muciniphila | B2UPK8 | 3.65 | 0.034816 | Low-fibre |
| Aminotransferase class I and II | B2UMN3 | 3.66 | 0.034816 | Low-fibre |
| Aminotransferase class I and II\|g__Akkermansia.s__Akkermansia_muciniphila | B2UMN3 | 3.66 | 0.034816 | Low-fibre |
| Peptidase M50 | B2UL91 | 3.78 | 0.034816 | Low-fibre |
| Peptidase M50\|g__Akkermansia.s__Akkermansia_muciniphila | B2UL91 | 3.78 | 0.034816 | Low-fibre |
| Regulatory protein GntR HTH | R7E362 | 3.67 | 0.034816 | Low-fibre |
| Regulatory protein GntR HTH\|g__Akkermansia.s__Akkermansia_muciniphila | R7E362 | 3.67 | 0.034816 | Low-fibre |
| Cyclic nucleotide-binding protein | B2UQF6 | 4.09 | 0.034816 | Low-fibre |
| Cyclic nucleotide-binding protein\|g__Akkermansia.s__Akkermansia_muciniphila | B2UQF6 | 4.09 | 0.034816 | Low-fibre |
| NO_NAME\|g__Parabacteroides.s__Parabacteroides_goldsteinii | R5DEL5 | 2.17 | 0.034816 | Low-fibre |
| Reverse transcriptase (RNA-dependent DNA polymerase) | R6D207 | 2.07 | 0.034816 | Low-fibre |
| CTP synthase | B2ULX0 | 4.07 | 0.035137 | Low-fibre |
| CTP synthase\|g__Akkermansia.s__Akkermansia_muciniphila | B2ULX0 | 4.07 | 0.035137 | Low-fibre |
| Ribonuclease | R6J1U3 | 3.81 | 0.035316 | Low-fibre |
| Ribonuclease\|g__Akkermansia.s__Akkermansia_muciniphila | R6J1U3 | 3.81 | 0.035316 | Low-fibre |
| Oligopeptide/dipeptide ABC transporter, ATPase subunit | B2UN55 | 3.85 | 0.035316 | Low-fibre |
| Oligopeptide/dipeptide ABC transporter, ATPase subunit\|g__Akkermansia.s__Akkermansia_muciniphila | B2UN55 | 3.85 | 0.035316 | Low-fibre |
| Glycosyl transferase family 2 | B2UPJ3 | 3.90 | 0.035316 | Low-fibre |
| Glycosyl transferase family 2\|g__Akkermansia.s__Akkermansia_muciniphila | B2UPJ3 | 3.90 | 0.035316 | Low-fibre |
| UDP-glucose 4-epimerase | R7DY48 | 3.78 | 0.035316 | Low-fibre |
| UDP-glucose 4-epimerase\|g__Akkermansia.s__Akkermansia_muciniphila | R7DY48 | 3.78 | 0.035316 | Low-fibre |
| Glycyl-tRNA synthetase | B2UMN6 | 3.97 | 0.035316 | Low-fibre |
| Glycyl-tRNA synthetase\|g__Akkermansia.s__Akkermansia_muciniphila | B2UMN6 | 3.97 | 0.035316 | Low-fibre |
| Glutamate--tRNA ligase | B2UN91 | 3.64 | 0.035316 | Low-fibre |
| Glutamate--tRNA ligase\|g__Akkermansia.s__Akkermansia_muciniphila | B2UN91 | 3.64 | 0.035316 | Low-fibre |
| TrkA-C domain protein | B2UL39 | 3.78 | 0.035316 | Low-fibre |
| TrkA-C domain protein\|g__Akkermansia.s__Akkermansia_muciniphila | B2UL39 | 3.78 | 0.035316 | Low-fibre |
| 30S ribosomal protein S1 | B2UME1 | 3.90 | 0.035316 | Low-fibre |
| 30S ribosomal protein S1\|g__Akkermansia.s__Akkermansia_muciniphila | B2UME1 | 3.90 | 0.035316 | Low-fibre |
| Glutaminase | B2UL96 | 3.71 | 0.035316 | Low-fibre |
| Glutaminase\|g__Akkermansia.s__Akkermansia_muciniphila | B2UL96 | 3.71 | 0.035316 | Low-fibre |
| Sulfate adenylyltransferase subunit 2 | B2URN8 | 3.60 | 0.035436 | Low-fibre |
| Sulfate adenylyltransferase subunit 2\|g__Akkermansia.s__Akkermansia_muciniphila | B2URN8 | 3.60 | 0.035436 | Low-fibre |
| Peptidoglycan-binding LysM | B2UQB9 | 3.64 | 0.035436 | Low-fibre |
| Peptidoglycan-binding LysM\|g__Akkermansia.s__Akkermansia_muciniphila | B2UQB9 | 3.64 | 0.035436 | Low-fibre |
| NO_NAME | R7EG48 | -2.62 | 0.035474 | High-fibre |
| NO_NAME\|g__Bacteroides.s__Bacteroides_xylanisolvens | R7EG48 | -2.62 | 0.035474 | High-fibre |
| NO_NAME | R7E596 | 3.88 | 0.035592 | Low-fibre |
| NO_NAME\|g__Akkermansia.s__Akkermansia_muciniphila | R7E596 | 3.88 | 0.035592 | Low-fibre |
| NO_NAME | B2UL28 | 3.65 | 0.035723 | Low-fibre |
| NO_NAME\|g__Akkermansia.s__Akkermansia_muciniphila | B2UL28 | 3.65 | 0.035723 | Low-fibre |
| Methionyl-tRNA synthetase | B2UPN4 | 3.67 | 0.035848 | Low-fibre |
| Methionyl-tRNA synthetase\|g__Akkermansia.s__Akkermansia_muciniphila | B2UPN4 | 3.67 | 0.035848 | Low-fibre |
| Aldo/keto reductase | B2UQU7 | 3.78 | 0.035848 | Low-fibre |
| Aldo/keto reductase\|g__Akkermansia.s__Akkermansia_muciniphila | B2UQU7 | 3.78 | 0.035848 | Low-fibre |
| NO_NAME | B2UL74 | 3.83 | 0.035848 | Low-fibre |
| NO_NAME\|g__Akkermansia.s__Akkermansia_muciniphila | B2UL74 | 3.83 | 0.035848 | Low-fibre |
| NO_NAME | B2UQP9 | 3.76 | 0.036062 | Low-fibre |
| NO_NAME\|g__Akkermansia.s__Akkermansia_muciniphila | B2UQP9 | 3.76 | 0.036062 | Low-fibre |
| Agmatine deiminase | B2ULR8 | 3.67 | 0.036062 | Low-fibre |
| Agmatine deiminase\|g__Akkermansia.s__Akkermansia_muciniphila | B2ULR8 | 3.67 | 0.036062 | Low-fibre |
| Tetratricopeptide TPR_2 repeat protein | B2UP12 | 3.87 | 0.036062 | Low-fibre |
| Tetratricopeptide TPR_2 repeat protein\|g__Akkermansia.s__Akkermansia_muciniphila | B2UP12 | 3.87 | 0.036062 | Low-fibre |
| NO_NAME | B2ULE8 | 3.70 | 0.036333 | Low-fibre |
| NO_NAME\|g__Akkermansia.s__Akkermansia_muciniphila | B2ULE8 | 3.70 | 0.036333 | Low-fibre |
| Peptidase membrane zinc metallopeptidase putative | B2UQJ2 | 3.75 | 0.036431 | Low-fibre |
| Peptidase membrane zinc metallopeptidase putative\|g__Akkermansia.s__Akkermansia_muciniphila | B2UQJ2 | 3.75 | 0.036431 | Low-fibre |
| NO_NAME | B2UR40 | 3.57 | 0.036464 | Low-fibre |
| NO_NAME\|g__Akkermansia.s__Akkermansia_muciniphila | B2UR40 | 3.57 | 0.036464 | Low-fibre |
| Ferripyochelin binding protein (Fbp) | B2UQ68 | 3.67 | 0.03662 | Low-fibre |
| Ferripyochelin binding protein (Fbp)\|g__Akkermansia.s__Akkermansia_muciniphila | B2UQ68 | 3.67 | 0.03662 | Low-fibre |
| Dihydroorotate dehydrogenase (quinone) | B2UNA2 | 3.70 | 0.036846 | Low-fibre |
| Dihydroorotate dehydrogenase (quinone)\|g__Akkermansia.s__Akkermansia_muciniphila | B2UNA2 | 3.70 | 0.036846 | Low-fibre |
| FeS assembly SUF system protein SufT | B2ULN3 | 3.64 | 0.03699 | Low-fibre |
| FeS assembly SUF system protein SufT\|g__Akkermansia.s__Akkermansia_muciniphila | B2ULN3 | 3.64 | 0.03699 | Low-fibre |
| NADH-quinone oxidoreductase subunit H | B2ULY6 | 3.22 | 0.03699 | Low-fibre |
| NADH-quinone oxidoreductase subunit H\|g__Akkermansia.s__Akkermansia_muciniphila | B2ULY6 | 3.22 | 0.03699 | Low-fibre |
| Type II secretion system protein E | R7E3Q6 | 3.59 | 0.036997 | Low-fibre |
| Type II secretion system protein E\|g__Akkermansia.s__Akkermansia_muciniphila | R7E3Q6 | 3.59 | 0.036997 | Low-fibre |
| Glycosyl transferase group 1 | B2UPM1 | 3.78 | 0.037049 | Low-fibre |
| Glycosyl transferase group 1\|g__Akkermansia.s__Akkermansia_muciniphila | B2UPM1 | 3.78 | 0.037049 | Low-fibre |
| NO_NAME | B2UNN1 | 4.23 | 0.037062 | Low-fibre |
| NO_NAME\|g__Akkermansia.s__Akkermansia_muciniphila | B2UNN1 | 4.23 | 0.037062 | Low-fibre |
| ABC transporter related | B2ULV6 | 3.68 | 0.037082 | Low-fibre |
| ABC transporter related\|g__Akkermansia.s__Akkermansia_muciniphila | B2ULV6 | 3.68 | 0.037082 | Low-fibre |
| LAO/AO transport system ATPase | B2UP59 | 3.55 | 0.037098 | Low-fibre |
| LAO/AO transport system ATPase\|g__Akkermansia.s__Akkermansia_muciniphila | B2UP59 | 3.55 | 0.037098 | Low-fibre |
| NO_NAME | R6K1I1 | 3.88 | 0.037098 | Low-fibre |
| NO_NAME\|g__Akkermansia.s__Akkermansia_muciniphila | R6K1I1 | 3.88 | 0.037098 | Low-fibre |
| Radical SAM domain protein | B2UQ91 | 3.80 | 0.037098 | Low-fibre |
| Radical SAM domain protein\|g__Akkermansia.s__Akkermansia_muciniphila | B2UQ91 | 3.80 | 0.037098 | Low-fibre |
| Translation initiation factor IF-2 | R7E3Y6 | 3.87 | 0.037189 | Low-fibre |
| Translation initiation factor IF-2\|g__Akkermansia.s__Akkermansia_muciniphila | R7E3Y6 | 3.87 | 0.037189 | Low-fibre |
| Metal dependent phosphohydrolase\|g__Akkermansia.s__Akkermansia_muciniphila | R6JBM3 | 3.75 | 0.037262 | Low-fibre |
| NO_NAME | C6INL9 | 2.28 | 0.037307 | Low-fibre |
| NO_NAME | R7DW31 | 4.06 | 0.03761 | Low-fibre |
| NO_NAME\|g__Akkermansia.s__Akkermansia_muciniphila | R7DW31 | 4.06 | 0.03761 | Low-fibre |
| MotA/TolQ/ExbB proton channel | B2UP16 | 3.68 | 0.03761 | Low-fibre |
| MotA/TolQ/ExbB proton channel\|g__Akkermansia.s__Akkermansia_muciniphila | B2UP16 | 3.68 | 0.03761 | Low-fibre |
| Peptidase S1 and S6 chymotrypsin/Hap | B2ULX6 | 3.76 | 0.037615 | Low-fibre |
| Peptidase S1 and S6 chymotrypsin/Hap\|g__Akkermansia.s__Akkermansia_muciniphila | B2ULX6 | 3.76 | 0.037615 | Low-fibre |
| Peptidase S15 | B2UN08 | 3.71 | 0.037615 | Low-fibre |
| Peptidase S15\|g__Akkermansia.s__Akkermansia_muciniphila | B2UN08 | 3.71 | 0.037615 | Low-fibre |
| Carboxyl transferase | B2UM96 | 3.74 | 0.037615 | Low-fibre |
| Carboxyl transferase\|g__Akkermansia.s__Akkermansia_muciniphila | B2UM96 | 3.74 | 0.037615 | Low-fibre |
| Ribosomal RNA small subunit methyltransferase I | B2UPG8 | 3.67 | 0.037615 | Low-fibre |
| Ribosomal RNA small subunit methyltransferase I\|g__Akkermansia.s__Akkermansia_muciniphila | B2UPG8 | 3.67 | 0.037615 | Low-fibre |
| Protein serine/threonine phosphatase | B2UQ96 | 4.03 | 0.037615 | Low-fibre |
| Protein serine/threonine phosphatase\|g__Akkermansia.s__Akkermansia_muciniphila | B2UQ96 | 4.03 | 0.037615 | Low-fibre |
| NO_NAME | B2UMY6 | 3.89 | 0.037615 | Low-fibre |
| NO_NAME\|g__Akkermansia.s__Akkermansia_muciniphila | B2UMY6 | 3.89 | 0.037615 | Low-fibre |
| Dihydroorotase, multifunctional complex type | B2ULR2 | 3.94 | 0.037615 | Low-fibre |
| Dihydroorotase, multifunctional complex type\|g__Akkermansia.s__Akkermansia_muciniphila | B2ULR2 | 3.94 | 0.037615 | Low-fibre |
| Phosphate binding protein | R6J0B6 | 3.67 | 0.037615 | Low-fibre |
| Phosphate binding protein\|g__Akkermansia.s__Akkermansia_muciniphila | R6J0B6 | 3.67 | 0.037615 | Low-fibre |
| Beta-hydroxyacyl-(Acyl-carrier-protein) dehydratase FabZ | B2UNL5 | 3.67 | 0.037615 | Low-fibre |
| Beta-hydroxyacyl-(Acyl-carrier-protein) dehydratase FabZ\|g__Akkermansia.s__Akkermansia_muciniphila | B2UNL5 | 3.67 | 0.037615 | Low-fibre |
| Glycosyl hydrolase BNR repeat-containing protein | B2UPI3 | 3.77 | 0.037615 | Low-fibre |
| Glycosyl hydrolase BNR repeat-containing protein\|g__Akkermansia.s__Akkermansia_muciniphila | B2UPI3 | 3.77 | 0.037615 | Low-fibre |
| NO_NAME | B2UQP7 | 3.42 | 0.037615 | Low-fibre |
| NO_NAME\|g__Akkermansia.s__Akkermansia_muciniphila | B2UQP7 | 3.42 | 0.037615 | Low-fibre |
| Hydroxymethylpyrimidine phosphate synthase ThiC | W4PLE3 | 2.18 | 0.037615 | Low-fibre |
| NO_NAME | B2UM44 | 4.08 | 0.037738 | Low-fibre |
| NO_NAME\|g__Akkermansia.s__Akkermansia_muciniphila | B2UM44 | 4.08 | 0.037738 | Low-fibre |
| Alcohol dehydrogenase zinc-binding domain protein | B2URD0 | 3.67 | 0.037891 | Low-fibre |
| Alcohol dehydrogenase zinc-binding domain protein\|g__Akkermansia.s__Akkermansia_muciniphila | B2URD0 | 3.67 | 0.037891 | Low-fibre |
| Uncharacterized Fe-S center protein, putative ferredoxin | B2UN06 | 3.62 | 0.037891 | Low-fibre |
| Uncharacterized Fe-S center protein, putative ferredoxin\|g__Akkermansia.s__Akkermansia_muciniphila | B2UN06 | 3.62 | 0.037891 | Low-fibre |
| Peptide chain release factor 1 | B2UL99 | 3.91 | 0.037891 | Low-fibre |
| Peptide chain release factor 1\|g__Akkermansia.s__Akkermansia_muciniphila | B2UL99 | 3.91 | 0.037891 | Low-fibre |
| Chorismate binding-like protein | B2UNP6 | 3.66 | 0.037891 | Low-fibre |
| Chorismate binding-like protein\|g__Akkermansia.s__Akkermansia_muciniphila | B2UNP6 | 3.66 | 0.037891 | Low-fibre |
| Glycoside hydrolase family 16 | R6IZJ9 | 3.63 | 0.037891 | Low-fibre |
| Glycoside hydrolase family 16\|g__Akkermansia.s__Akkermansia_muciniphila | R6IZJ9 | 3.63 | 0.037891 | Low-fibre |
| Multiple antibiotic resistance (MarC)-related protein | B2UQV9 | 3.81 | 0.037891 | Low-fibre |
| Multiple antibiotic resistance (MarC)-related protein\|g__Akkermansia.s__Akkermansia_muciniphila | B2UQV9 | 3.81 | 0.037891 | Low-fibre |
| Glycoside hydrolase family 57 | B2UN71 | 3.70 | 0.037891 | Low-fibre |
| Glycoside hydrolase family 57\|g__Akkermansia.s__Akkermansia_muciniphila | B2UN71 | 3.70 | 0.037891 | Low-fibre |
| Transposase | R6V8K5 | 2.49 | 0.037891 | Low-fibre |
| NO_NAME | R7DTV7 | 3.67 | 0.037923 | Low-fibre |
| NO_NAME\|g__Akkermansia.s__Akkermansia_muciniphila | R7DTV7 | 3.67 | 0.037923 | Low-fibre |
| Metal dependent phosphohydrolase\|g__Bacteroides.s__Bacteroides_xylanisolvens | D4J4D6 | -3.40 | 0.037992 | High-fibre |
| Glycosyl transferase family 2 | B2UR11 | 3.72 | 0.038039 | Low-fibre |
| Glycosyl transferase family 2\|g__Akkermansia.s__Akkermansia_muciniphila | B2UR11 | 3.72 | 0.038039 | Low-fibre |
| Squalene-hopene cyclase-like protein | R7E0U1 | 3.76 | 0.038039 | Low-fibre |
| Squalene-hopene cyclase-like protein\|g__Akkermansia.s__Akkermansia_muciniphila | R7E0U1 | 3.76 | 0.038039 | Low-fibre |
| Glycoside hydrolase family 37 | B2URK0 | 3.72 | 0.038039 | Low-fibre |
| Glycoside hydrolase family 37\|g__Akkermansia.s__Akkermansia_muciniphila | B2URK0 | 3.72 | 0.038039 | Low-fibre |
| NO_NAME | R6JE05 | 3.96 | 0.038039 | Low-fibre |
| NO_NAME\|g__Akkermansia.s__Akkermansia_muciniphila | R6JE05 | 3.96 | 0.038039 | Low-fibre |
| Elongation factor P | B2UQH2 | 3.70 | 0.038039 | Low-fibre |
| Elongation factor P\|g__Akkermansia.s__Akkermansia_muciniphila | B2UQH2 | 3.70 | 0.038039 | Low-fibre |
| 6,7-dimethyl-8-ribityllumazine synthase | B2UNE9 | 3.54 | 0.038039 | Low-fibre |
| 6,7-dimethyl-8-ribityllumazine synthase\|g__Akkermansia.s__Akkermansia_muciniphila | B2UNE9 | 3.54 | 0.038039 | Low-fibre |
| NO_NAME | B2UMB0 | 3.69 | 0.038039 | Low-fibre |
| NO_NAME\|g__Akkermansia.s__Akkermansia_muciniphila | B2UMB0 | 3.69 | 0.038039 | Low-fibre |
| 6-phosphofructokinase | B2UL30 | 3.83 | 0.038067 | Low-fibre |
| 6-phosphofructokinase\|g__Akkermansia.s__Akkermansia_muciniphila | B2UL30 | 3.83 | 0.038067 | Low-fibre |
| NO_NAME | B2UP10 | 3.90 | 0.038263 | Low-fibre |
| NO_NAME\|g__Akkermansia.s__Akkermansia_muciniphila | B2UP10 | 3.90 | 0.038263 | Low-fibre |
| Type I phosphodiesterase/nucleotide pyrophosphatase | B2UNC3 | 3.69 | 0.038338 | Low-fibre |
| Type I phosphodiesterase/nucleotide pyrophosphatase\|g__Akkermansia.s__Akkermansia_muciniphila | B2UNC3 | 3.69 | 0.038338 | Low-fibre |
| Ribosomal RNA small subunit methyltransferase H | R7DUT1 | 3.73 | 0.038373 | Low-fibre |
| Ribosomal RNA small subunit methyltransferase H\|g__Akkermansia.s__Akkermansia_muciniphila | R7DUT1 | 3.73 | 0.038373 | Low-fibre |
| O-acetylhomoserine/O-acetylserine sulfhydrylase | B2UQ89 | 3.60 | 0.038373 | Low-fibre |
| O-acetylhomoserine/O-acetylserine sulfhydrylase\|g__Akkermansia.s__Akkermansia_muciniphila | B2UQ89 | 3.60 | 0.038373 | Low-fibre |
| Glycosyl hydrolase family 109 protein 1 | B2UL75 | 3.98 | 0.038373 | Low-fibre |
| Glycosyl hydrolase family 109 protein 1\|g__Akkermansia.s__Akkermansia_muciniphila | B2UL75 | 3.98 | 0.038373 | Low-fibre |
| Sulfate adenylyltransferase subunit 1 | B2URN7 | 4.36 | 0.038373 | Low-fibre |
| Sulfate adenylyltransferase subunit 1\|g__Akkermansia.s__Akkermansia_muciniphila | B2URN7 | 4.36 | 0.038373 | Low-fibre |
| NO_NAME | B2UQC1 | 3.80 | 0.038373 | Low-fibre |
| NO_NAME\|g__Akkermansia.s__Akkermansia_muciniphila | B2UQC1 | 3.80 | 0.038373 | Low-fibre |
| Site-specific DNA-methyltransferase (Adenine-specific) | B2UNK3 | 3.50 | 0.038373 | Low-fibre |
| Site-specific DNA-methyltransferase (Adenine-specific)\|g__Akkermansia.s__Akkermansia_muciniphila | B2UNK3 | 3.50 | 0.038373 | Low-fibre |
| 50S ribosomal protein L5 | B2UQM3 | 3.67 | 0.038373 | Low-fibre |
| 50S ribosomal protein L5\|g__Akkermansia.s__Akkermansia_muciniphila | B2UQM3 | 3.67 | 0.038373 | Low-fibre |
| Elongation factor Ts | B2UPX5 | 3.73 | 0.038373 | Low-fibre |
| Elongation factor Ts\|g__Akkermansia.s__Akkermansia_muciniphila | B2UPX5 | 3.73 | 0.038373 | Low-fibre |
| Binding-protein-dependent transport systems inner membrane component | B2ULU3 | 3.67 | 0.038373 | Low-fibre |
| Binding-protein-dependent transport systems inner membrane component\|g__Akkermansia.s__Akkermansia_muciniphila | B2ULU3 | 3.67 | 0.038373 | Low-fibre |
| NO_NAME\|g__Bacteroides.s__Bacteroides_ovatus | E4MER3 | 2.16 | 0.038373 | Low-fibre |
| integrase, partial | UPI000468EDAD | 2.42 | 0.038373 | Low-fibre |
| Aminopeptidase | B2URM0 | 3.84 | 0.038386 | Low-fibre |
| Aminopeptidase\|g__Akkermansia.s__Akkermansia_muciniphila | B2URM0 | 3.84 | 0.038386 | Low-fibre |
| Protein GrpE | B2UL08 | 3.55 | 0.03845 | Low-fibre |
| Protein GrpE\|g__Akkermansia.s__Akkermansia_muciniphila | B2UL08 | 3.55 | 0.03845 | Low-fibre |
| Excisionase family DNA binding domain-containing protein\|g__Bacteroides.s__Bacteroides_ovatus | I9RLL9 | 2.16 | 0.038497 | Low-fibre |
| NO_NAME | R9IDU3 | 3.06 | 0.038693 | Low-fibre |
| TatD-related deoxyribonuclease | B2URL7 | 3.74 | 0.038693 | Low-fibre |
| TatD-related deoxyribonuclease\|g__Akkermansia.s__Akkermansia_muciniphila | B2URL7 | 3.74 | 0.038693 | Low-fibre |
| NO_NAME | R9KE80 | 2.50 | 0.038812 | Low-fibre |
| FeS assembly ATPase SufC | B2UNV5 | 3.63 | 0.038878 | Low-fibre |
| FeS assembly ATPase SufC\|g__Akkermansia.s__Akkermansia_muciniphila | B2UNV5 | 3.63 | 0.038878 | Low-fibre |
| NO_NAME | K1SMZ8 | -3.46 | 0.039045 | High-fibre |
| Phosphate import ATP-binding protein PstB | B2URP2 | 3.49 | 0.03936 | Low-fibre |
| Phosphate import ATP-binding protein PstB\|g__Akkermansia.s__Akkermansia_muciniphila | B2URP2 | 3.49 | 0.03936 | Low-fibre |
| Cytochrome c class I | B2UN77 | 3.71 | 0.039623 | Low-fibre |
| Cytochrome c class I\|g__Akkermansia.s__Akkermansia_muciniphila | B2UN77 | 3.71 | 0.039623 | Low-fibre |
| NO_NAME | C6ZBB4 | 2.28 | 0.039623 | Low-fibre |
| NO_NAME | R6KB58 | 2.03 | 0.03964 | Low-fibre |
| NO_NAME\|g__Bacteroides.s__Bacteroides_ovatus | I9T2P7 | 2.25 | 0.03964 | Low-fibre |
| Exodeoxyribonuclease III Xth | B2UQQ5 | 3.70 | 0.039973 | Low-fibre |
| Exodeoxyribonuclease III Xth\|g__Akkermansia.s__Akkermansia_muciniphila | B2UQQ5 | 3.70 | 0.039973 | Low-fibre |
| Acyltransferase 3 | R6JDF9 | 3.60 | 0.039973 | Low-fibre |
| Acyltransferase 3\|g__Akkermansia.s__Akkermansia_muciniphila | R6JDF9 | 3.60 | 0.039973 | Low-fibre |
| 3-oxoacyl-[acyl-carrier-protein] synthase 3 | B2UPJ0 | 3.83 | 0.040129 | Low-fibre |
| 3-oxoacyl-[acyl-carrier-protein] synthase 3\|g__Akkermansia.s__Akkermansia_muciniphila | B2UPJ0 | 3.83 | 0.040129 | Low-fibre |
| Acetolactate synthase | B2UM25 | 3.83 | 0.040205 | Low-fibre |
| Acetolactate synthase\|g__Akkermansia.s__Akkermansia_muciniphila | B2UM25 | 3.83 | 0.040205 | Low-fibre |
| NO_NAME | B2UN82 | 3.52 | 0.040255 | Low-fibre |
| NO_NAME\|g__Akkermansia.s__Akkermansia_muciniphila | B2UN82 | 3.52 | 0.040255 | Low-fibre |
| Cell wall hydrolase/autolysin | B2UPW6 | 3.49 | 0.040602 | Low-fibre |
| Cell wall hydrolase/autolysin\|g__Akkermansia.s__Akkermansia_muciniphila | B2UPW6 | 3.49 | 0.040602 | Low-fibre |
| ATP-dependent helicase HrpA | B2UQA6 | 3.60 | 0.040908 | Low-fibre |
| ATP-dependent helicase HrpA\|g__Akkermansia.s__Akkermansia_muciniphila | B2UQA6 | 3.60 | 0.040908 | Low-fibre |
| RDD domain containing protein | B2ULY1 | 4.01 | 0.040908 | Low-fibre |
| RDD domain containing protein\|g__Akkermansia.s__Akkermansia_muciniphila | B2ULY1 | 4.01 | 0.040908 | Low-fibre |
| NO_NAME | B2UMB1 | 3.50 | 0.04114 | Low-fibre |
| NO_NAME\|g__Akkermansia.s__Akkermansia_muciniphila | B2UMB1 | 3.50 | 0.04114 | Low-fibre |
| Excisionase family DNA binding domain-containing protein | I9RLL9 | 2.13 | 0.041204 | Low-fibre |
| Acyltransferase 3 | R7E1D6 | 3.45 | 0.041278 | Low-fibre |
| Acyltransferase 3\|g__Akkermansia.s__Akkermansia_muciniphila | R7E1D6 | 3.45 | 0.041278 | Low-fibre |
| NO_NAME\|g__Bacteroides.s__Bacteroides_ovatus | C6Z6Z7 | 2.21 | 0.041278 | Low-fibre |
| NO_NAME | E4MER3 | 2.16 | 0.041314 | Low-fibre |
| NO_NAME | R7E254 | 3.60 | 0.04146 | Low-fibre |
| NO_NAME\|g__Akkermansia.s__Akkermansia_muciniphila | R7E254 | 3.60 | 0.04146 | Low-fibre |
| NO_NAME | B2UPE3 | 3.77 | 0.04146 | Low-fibre |
| NO_NAME\|g__Akkermansia.s__Akkermansia_muciniphila | B2UPE3 | 3.77 | 0.04146 | Low-fibre |
| DNA protecting protein DprA | B2UNG9 | 3.66 | 0.042078 | Low-fibre |
| DNA protecting protein DprA\|g__Akkermansia.s__Akkermansia_muciniphila | B2UNG9 | 3.66 | 0.042078 | Low-fibre |
| tRNA-dihydrouridine synthase | R7E3W1 | 3.83 | 0.042078 | Low-fibre |
| tRNA-dihydrouridine synthase\|g__Akkermansia.s__Akkermansia_muciniphila | R7E3W1 | 3.83 | 0.042078 | Low-fibre |
| Phenylalanine--tRNA ligase beta subunit | B2UP38 | 3.68 | 0.042078 | Low-fibre |
| Phenylalanine--tRNA ligase beta subunit\|g__Akkermansia.s__Akkermansia_muciniphila | B2UP38 | 3.68 | 0.042078 | Low-fibre |
| Non-canonical purine NTP pyrophosphatase | B2UQQ6 | 3.78 | 0.042078 | Low-fibre |
| Non-canonical purine NTP pyrophosphatase\|g__Akkermansia.s__Akkermansia_muciniphila | B2UQQ6 | 3.78 | 0.042078 | Low-fibre |
| Thioredoxin domain | B2UN47 | 3.48 | 0.042224 | Low-fibre |
| Thioredoxin domain\|g__Akkermansia.s__Akkermansia_muciniphila | B2UN47 | 3.48 | 0.042224 | Low-fibre |
| Sua5/YciO/YrdC/YwlC family protein | B2UMG3 | 3.81 | 0.042261 | Low-fibre |
| Sua5/YciO/YrdC/YwlC family protein\|g__Akkermansia.s__Akkermansia_muciniphila | B2UMG3 | 3.81 | 0.042261 | Low-fibre |
| NO_NAME | K5YTB0 | -2.01 | 0.042298 | High-fibre |
| L-threonine 3-dehydrogenase | B2UMC7 | 3.59 | 0.042298 | Low-fibre |
| L-threonine 3-dehydrogenase\|g__Akkermansia.s__Akkermansia_muciniphila | B2UMC7 | 3.59 | 0.042298 | Low-fibre |
| Transcriptional regulator, TraR/DksA family | B2ULP7 | 3.77 | 0.042298 | Low-fibre |
| Transcriptional regulator, TraR/DksA family\|g__Akkermansia.s__Akkermansia_muciniphila | B2ULP7 | 3.77 | 0.042298 | Low-fibre |
| HhH-GPD family protein | B2UNR9 | 3.67 | 0.042316 | Low-fibre |
| HhH-GPD family protein\|g__Akkermansia.s__Akkermansia_muciniphila | B2UNR9 | 3.67 | 0.042316 | Low-fibre |
| NO_NAME | R5B8B8 | 2.98 | 0.04238 | Low-fibre |
| Nuclease (SNase domain protein) | B2UQ78 | 3.42 | 0.042481 | Low-fibre |
| Nuclease (SNase domain protein)\|g__Akkermansia.s__Akkermansia_muciniphila | B2UQ78 | 3.42 | 0.042481 | Low-fibre |
| dTDP-4-dehydrorhamnose reductase | R6JBL5 | 3.47 | 0.042521 | Low-fibre |
| dTDP-4-dehydrorhamnose reductase\|g__Akkermansia.s__Akkermansia_muciniphila | R6JBL5 | 3.47 | 0.042521 | Low-fibre |
| RNP-1 like RNA-binding protein | B2UQR4 | 3.74 | 0.042895 | Low-fibre |
| RNP-1 like RNA-binding protein\|g__Akkermansia.s__Akkermansia_muciniphila | B2UQR4 | 3.74 | 0.042895 | Low-fibre |
| Arginine--tRNA ligase | R6JBR5 | 3.69 | 0.042895 | Low-fibre |
| Arginine--tRNA ligase\|g__Akkermansia.s__Akkermansia_muciniphila | R6JBR5 | 3.69 | 0.042895 | Low-fibre |
| NO_NAME | B2UQ61 | 3.42 | 0.042895 | Low-fibre |
| NO_NAME\|g__Akkermansia.s__Akkermansia_muciniphila | B2UQ61 | 3.42 | 0.042895 | Low-fibre |
| NO_NAME\|g__Bacteroides.s__Bacteroides_ovatus | C6ZBB4 | 2.44 | 0.042895 | Low-fibre |
| Helicase c2 | B2URD7 | 3.68 | 0.042961 | Low-fibre |
| Helicase c2\|g__Akkermansia.s__Akkermansia_muciniphila | B2URD7 | 3.68 | 0.042961 | Low-fibre |
| NO_NAME | A0A016IZ22 | 2.04 | 0.042961 | Low-fibre |
| MobN1\|g__Bacteroides.s__Bacteroides_ovatus | Q07719 | -2.11 | 0.043134 | High-fibre |
| NO_NAME | C6Z6X8 | 2.10 | 0.043169 | Low-fibre |
| NO_NAME | K9E5M8 | 2.01 | 0.043293 | Low-fibre |
| NO_NAME | R6VGK1 | 2.18 | 0.043325 | Low-fibre |
| NO_NAME\|g__Bacteroides.s__Bacteroides_ovatus | R6VGK1 | 2.18 | 0.043325 | Low-fibre |
| NO_NAME | B2ULD7 | 3.49 | 0.043608 | Low-fibre |
| NO_NAME\|g__Akkermansia.s__Akkermansia_muciniphila | B2ULD7 | 3.49 | 0.043608 | Low-fibre |
| NO_NAME\|g__Eubacterium.s__Eubacterium_plexicaudatum | N1ZYY3 | -3.65 | 0.043697 | High-fibre |
| 3-deoxy-manno-octulosonate cytidylyltransferase | B2ULW9 | 3.55 | 0.043905 | Low-fibre |
| 3-deoxy-manno-octulosonate cytidylyltransferase\|g__Akkermansia.s__Akkermansia_muciniphila | B2ULW9 | 3.55 | 0.043905 | Low-fibre |
| NO_NAME | C6INK7 | 2.09 | 0.043905 | Low-fibre |
| NO_NAME\|g__Bacteroides.s__Bacteroides_ovatus | C6INK7 | 2.09 | 0.043905 | Low-fibre |
| tRNA pseudouridine synthase A | B2UNB6 | 3.69 | 0.044674 | Low-fibre |
| tRNA pseudouridine synthase A\|g__Akkermansia.s__Akkermansia_muciniphila | B2UNB6 | 3.69 | 0.044674 | Low-fibre |
| 30S ribosomal protein S14 type Z | R7BRG3 | 2.51 | 0.044674 | Low-fibre |
| NO_NAME | UPI0004692D69 | 2.45 | 0.044674 | Low-fibre |
| NO_NAME | B2UNK5 | 3.42 | 0.044858 | Low-fibre |
| NO_NAME\|g__Akkermansia.s__Akkermansia_muciniphila | B2UNK5 | 3.42 | 0.044858 | Low-fibre |
| NO_NAME | B2UKW5 | 3.43 | 0.044935 | Low-fibre |
| NO_NAME\|g__Akkermansia.s__Akkermansia_muciniphila | B2UKW5 | 3.43 | 0.044935 | Low-fibre |
| NO_NAME | B2UNF9 | 3.45 | 0.045141 | Low-fibre |
| NO_NAME\|g__Akkermansia.s__Akkermansia_muciniphila | B2UNF9 | 3.45 | 0.045141 | Low-fibre |
| NO_NAME | B2UQ08 | 3.87 | 0.045141 | Low-fibre |
| NO_NAME\|g__Akkermansia.s__Akkermansia_muciniphila | B2UQ08 | 3.87 | 0.045141 | Low-fibre |
| Major facilitator superfamily MFS_1 | R6J5R9 | 3.43 | 0.045141 | Low-fibre |
| Major facilitator superfamily MFS_1\|g__Akkermansia.s__Akkermansia_muciniphila | R6J5R9 | 3.43 | 0.045141 | Low-fibre |
| NO_NAME | N2AT21 | 4.39 | 0.045141 | Low-fibre |
| NO_NAME | I9AEG6 | 2.93 | 0.045209 | Low-fibre |
| Metallophosphoesterase | B2UNJ9 | 3.83 | 0.045245 | Low-fibre |
| Metallophosphoesterase\|g__Akkermansia.s__Akkermansia_muciniphila | B2UNJ9 | 3.83 | 0.045245 | Low-fibre |
| NO_NAME | B2ULA3 | 3.36 | 0.045423 | Low-fibre |
| NO_NAME\|g__Akkermansia.s__Akkermansia_muciniphila | B2ULA3 | 3.36 | 0.045423 | Low-fibre |
| Peptidase M23 | R7E1P5 | 3.60 | 0.045457 | Low-fibre |
| Peptidase M23\|g__Akkermansia.s__Akkermansia_muciniphila | R7E1P5 | 3.60 | 0.045457 | Low-fibre |
| UDP-N-acetylmuramoyl-L-alanyl-D-glutamate--2,6-diaminopimelate ligase | B2UPL3 | 3.54 | 0.045697 | Low-fibre |
| UDP-N-acetylmuramoyl-L-alanyl-D-glutamate--2,6-diaminopimelate ligase\|g__Akkermansia.s__Akkermansia_muciniphila | B2UPL3 | 3.54 | 0.045697 | Low-fibre |
| Peptidase M20 | B2UMM1 | 3.38 | 0.045697 | Low-fibre |
| Peptidase M20\|g__Akkermansia.s__Akkermansia_muciniphila | B2UMM1 | 3.38 | 0.045697 | Low-fibre |
| NO_NAME | K5YSB6 | -6.15 | 0.045795 | High-fibre |
| tRNA (guanine-N(1)-)-methyltransferase | B2UMD8 | 3.55 | 0.045795 | Low-fibre |
| tRNA (guanine-N(1)-)-methyltransferase\|g__Akkermansia.s__Akkermansia_muciniphila | B2UMD8 | 3.55 | 0.045795 | Low-fibre |
| Argininosuccinate synthase | B2UP27 | 3.46 | 0.045795 | Low-fibre |
| Argininosuccinate synthase\|g__Akkermansia.s__Akkermansia_muciniphila | B2UP27 | 3.46 | 0.045795 | Low-fibre |
| Cell division protein ftsA | B2ULV3 | 3.74 | 0.045795 | Low-fibre |
| Cell division protein ftsA\|g__Akkermansia.s__Akkermansia_muciniphila | B2ULV3 | 3.74 | 0.045795 | Low-fibre |
| NO_NAME | B2ULW8 | 3.46 | 0.045795 | Low-fibre |
| NO_NAME\|g__Akkermansia.s__Akkermansia_muciniphila | B2ULW8 | 3.46 | 0.045795 | Low-fibre |
| NO_NAME | UPI000468651B | 2.67 | 0.045811 | Low-fibre |
| NO_NAME | C6INH5 | 2.17 | 0.045925 | Low-fibre |
| NO_NAME\|g__Bacteroides.s__Bacteroides_ovatus | C6INH5 | 2.17 | 0.045925 | Low-fibre |
| NO_NAME | B2UNR6 | 3.50 | 0.045942 | Low-fibre |
| NO_NAME\|g__Akkermansia.s__Akkermansia_muciniphila | B2UNR6 | 3.50 | 0.045942 | Low-fibre |
| Acyltransferase 3 | R6JCF8 | 3.70 | 0.04637 | Low-fibre |
| Acyltransferase 3\|g__Akkermansia.s__Akkermansia_muciniphila | R6JCF8 | 3.70 | 0.04637 | Low-fibre |
| Ubiquinone/menaquinone biosynthesis methyltransferase | B2ULA5 | 3.32 | 0.046787 | Low-fibre |
| Ubiquinone/menaquinone biosynthesis methyltransferase\|g__Akkermansia.s__Akkermansia_muciniphila | B2ULA5 | 3.32 | 0.046787 | Low-fibre |
| NO_NAME | B2ULV1 | 3.49 | 0.047151 | Low-fibre |
| NO_NAME\|g__Akkermansia.s__Akkermansia_muciniphila | B2ULV1 | 3.49 | 0.047151 | Low-fibre |
| NO_NAME | B2UQE9 | 3.43 | 0.047158 | Low-fibre |
| NO_NAME\|g__Akkermansia.s__Akkermansia_muciniphila | B2UQE9 | 3.43 | 0.047158 | Low-fibre |
| NO_NAME | B2UPC7 | 3.67 | 0.047247 | Low-fibre |
| NO_NAME\|g__Akkermansia.s__Akkermansia_muciniphila | B2UPC7 | 3.67 | 0.047247 | Low-fibre |
| Miro domain protein | B2UMM4 | 3.59 | 0.047247 | Low-fibre |
| Miro domain protein\|g__Akkermansia.s__Akkermansia_muciniphila | B2UMM4 | 3.59 | 0.047247 | Low-fibre |
| NO_NAME | B2UNW4 | 3.66 | 0.047384 | Low-fibre |
| NO_NAME\|g__Akkermansia.s__Akkermansia_muciniphila | B2UNW4 | 3.66 | 0.047384 | Low-fibre |
| NO_NAME | B2UPN0 | 3.39 | 0.047585 | Low-fibre |
| NO_NAME\|g__Akkermansia.s__Akkermansia_muciniphila | B2UPN0 | 3.39 | 0.047585 | Low-fibre |
| FAD dependent oxidoreductase | B2ULR7 | 3.72 | 0.04773 | Low-fibre |
| FAD dependent oxidoreductase\|g__Akkermansia.s__Akkermansia_muciniphila | B2ULR7 | 3.72 | 0.04773 | Low-fibre |
| Putative ATP-binding protein | B2UMY3 | 3.79 | 0.047747 | Low-fibre |
| Putative ATP-binding protein\|g__Akkermansia.s__Akkermansia_muciniphila | B2UMY3 | 3.79 | 0.047747 | Low-fibre |
| Methylmalonyl-CoA epimerase | R7E6F2 | 3.44 | 0.047747 | Low-fibre |
| Methylmalonyl-CoA epimerase\|g__Akkermansia.s__Akkermansia_muciniphila | R7E6F2 | 3.44 | 0.047747 | Low-fibre |
| Glycosyl transferase family 2 | B2UPJ2 | 3.60 | 0.047747 | Low-fibre |
| Glycosyl transferase family 2\|g__Akkermansia.s__Akkermansia_muciniphila | B2UPJ2 | 3.60 | 0.047747 | Low-fibre |
| NO_NAME | G9S2T6 | 2.38 | 0.047747 | Low-fibre |
| NO_NAME | UPI000469F30F | 2.48 | 0.047747 | Low-fibre |
| Phosphatidylserine decarboxylase-related | B2UM27 | 3.54 | 0.047841 | Low-fibre |
| Phosphatidylserine decarboxylase-related\|g__Akkermansia.s__Akkermansia_muciniphila | B2UM27 | 3.54 | 0.047841 | Low-fibre |
| NO_NAME | B2URL5 | 3.28 | 0.047841 | Low-fibre |
| NO_NAME\|g__Akkermansia.s__Akkermansia_muciniphila | B2URL5 | 3.28 | 0.047841 | Low-fibre |
| Transposase | R6LNS5 | 2.36 | 0.047841 | Low-fibre |
| NO_NAME | B2UMY5 | 3.46 | 0.047914 | Low-fibre |
| NO_NAME\|g__Akkermansia.s__Akkermansia_muciniphila | B2UMY5 | 3.46 | 0.047914 | Low-fibre |
| 2,3-bisphosphoglycerate-independent phosphoglycerate mutase | B2UMU5 | 3.77 | 0.047914 | Low-fibre |
| 2,3-bisphosphoglycerate-independent phosphoglycerate mutase\|g__Akkermansia.s__Akkermansia_muciniphila | B2UMU5 | 3.77 | 0.047914 | Low-fibre |
| Hydroxyethylthiazole kinase | B2UP55 | 3.39 | 0.048235 | Low-fibre |
| Hydroxyethylthiazole kinase\|g__Akkermansia.s__Akkermansia_muciniphila | B2UP55 | 3.39 | 0.048235 | Low-fibre |
| NO_NAME\|g__Parabacteroides.s__Parabacteroides_goldsteinii | K5YSB6 | -6.01 | 0.048965 | High-fibre |
| Beta-N-acetylhexosaminidase | B2UPP0 | 3.57 | 0.049254 | Low-fibre |
| Beta-N-acetylhexosaminidase\|g__Akkermansia.s__Akkermansia_muciniphila | B2UPP0 | 3.57 | 0.049254 | Low-fibre |
| Plasmid transfer protein\|g__Bacteroides.s__Bacteroides_ovatus | D7I7J6 | 2.21 | 0.049296 | Low-fibre |
| NO_NAME | R9NF06 | -3.43 | 0.049299 | High-fibre |
| NO_NAME | B2UR25 | 3.40 | 0.049299 | Low-fibre |
| Beta-lactamase domain protein | B2UMD2 | 3.31 | 0.049299 | Low-fibre |
| Beta-lactamase domain protein\|g__Akkermansia.s__Akkermansia_muciniphila | B2UMD2 | 3.31 | 0.049299 | Low-fibre |
| Eight transmembrane protein EpsH | B2UL19 | 3.61 | 0.04942 | Low-fibre |
| Eight transmembrane protein EpsH\|g__Akkermansia.s__Akkermansia_muciniphila | B2UL19 | 3.61 | 0.04942 | Low-fibre |
| NO_NAME | B2UL89 | 3.56 | 0.049619 | Low-fibre |
| NO_NAME\|g__Akkermansia.s__Akkermansia_muciniphila | B2UL89 | 3.56 | 0.049619 | Low-fibre |
| N-acetylmuramoyl-L-alanine amidase family 2 | B2UQK0 | 3.36 | 0.049739 | Low-fibre |
| N-acetylmuramoyl-L-alanine amidase family 2\|g__Akkermansia.s__Akkermansia_muciniphila | B2UQK0 | 3.36 | 0.049739 | Low-fibre |
| ATPase associated with various cellular activities AAA_3 | R6JC32 | 3.37 | 0.049739 | Low-fibre |
| ATPase associated with various cellular activities AAA_3\|g__Akkermansia.s__Akkermansia_muciniphila | R6JC32 | 3.37 | 0.049739 | Low-fibre |
| NO_NAME\|g__Akkermansia.s__Akkermansia_muciniphila | B2UR25 | 3.37 | 0.049739 | Low-fibre |
| NO_NAME | B2UP97 | 3.25 | 0.049739 | Low-fibre |
| NO_NAME\|g__Akkermansia.s__Akkermansia_muciniphila | B2UP97 | 3.25 | 0.049739 | Low-fibre |
| Conserved protein found in conjugate transposon | Q8A5E3 | 2.29 | 0.049953 | Low-fibre |

**Supplementary Table 4.** Integrated metagenomic pathway analysis showing differentially abundant bacteria.

| **Pathway ID** | **LogFC** | **Adjusted *P*-value** | **Direction** |
| --- | --- | --- | --- |
| UNINTEGRATED\|g__*Akkermansia.*s__*Akkermansia_muciniphila* | 5.39 | 0.0002 | Higher in Low-fibre |
| PWY-4702: phytate degradation I | 3.86 | 0.0002 | Higher in Low-fibre |
| P108-PWY: pyruvate fermentation to propanoate I | 3.39 | 0.0004 | Higher in Low-fibre |
| PYRIDOXSYN-PWY: pyridoxal 5'-phosphate biosynthesis I | 3.28 | 0.0019 | Higher in Low-fibre |
| PWY0-162: superpathway of pyrimidine ribonucleotides de novo biosynthesis | 2.50 | 0.0028 | Higher in Low-fibre |
| PWY-7221: guanosine ribonucleotides de novo biosynthesis\|g__*Akkermansia.s__Akkermansia_muciniphila* | 4.58 | 0.0028 | Higher in Low-fibre |
| PWY-7219: adenosine ribonucleotides de novo biosynthesis\|g__*Akkermansia.s__Akkermansia_muciniphila* | 4.07 | 0.0065 | Higher in Low-fibre |
| PWY0-845: superpathway of pyridoxal 5'-phosphate biosynthesis and salvage | 3.59 | 0.0329 | Higher in Low-fibre |
| PWY-5345: superpathway of L-methionine biosynthesis (by sulfhydrylation) | 2.16 | 0.0329 | Higher in Low-fibre |
| SO4ASSIM-PWY: sulfate reduction I (assimilatory) | 2.38 | 0.0493 | Higher in Low-fibre |

Legend: FC, fold change.

**Supplementary Table 5.** Top 10 genes per cell cluster of differentially expressed genes in the heart of low-fibre and high-fibre offspring detected using single-cell RNA-sequencing.

| **Gene name** | **Cell type** | **P-value** | **FDR P-value** | **Log2Fold change** | **Expression in High fibre** | **Expression in Low fibre** |
| --- | --- | --- | --- | --- | --- | --- |
| *Dnaja1* | B cells | 5.45E-34 | 2.35E-30 | 1.17 | 10.92 | 4.86 |
| *Hspa1b* | B cells | 1.13E-30 | 2.44E-27 | 1.97 | 9.18 | 2.34 |
| *Hspa1a* | B cells | 9.37E-23 | 1.34E-19 | 2.09 | 7.86 | 1.85 |
| *Hsp90aa1* | B cells | 7.17E-20 | 7.71E-17 | 0.92 | 13.63 | 7.20 |
| *Hsph1* | B cells | 1.88E-16 | 1.61E-13 | 1.51 | 2.82 | 0.99 |
| *Dnajb1* | B cells | 2.36E-14 | 1.69E-11 | 1.41 | 3.14 | 1.18 |
| *Hspa8* | B cells | 3.29E-13 | 2.02E-10 | 0.51 | 17.87 | 12.56 |
| *Hsp90ab1* | B cells | 3.05E-12 | 1.64E-09 | 0.52 | 21.90 | 15.30 |
| *Btg2* | B cells | 4.79E-12 | 2.29E-09 | 0.64 | 6.80 | 4.36 |
| *Apoe* | B cells | 3.42E-11 | 1.43E-08 | 1.09 | 6.81 | 3.20 |
| *Dnaja1* | DC-like cells | 1.17E-06 | 8.78E-03 | 0.75 | 10.24 | 6.10 |
| *Sec22a* | DC-like cells | 1.52E-05 | 5.69E-02 |  | 0.11 | 0.00 |
| *Lyz2* | Endocard | 5.71E-10 | 4.78E-06 | 1.61 | 4.67 | 1.53 |
| *Rbm3* | Endocard | 2.88E-09 | 1.21E-05 | 0.61 | 2.36 | 1.54 |
| *Gm47283* | Endocard | 4.99E-08 | 1.39E-04 | -0.99 | 0.69 | 1.37 |
| *Cd52* | Endocard | 9.11E-07 | 1.90E-03 | 2.05 | 0.24 | 0.06 |
| *Ost4* | Endocard | 7.02E-06 | 1.11E-02 | 0.16 | 2.93 | 2.62 |
| *Sult1a1* | Endocard | 7.97E-06 | 1.11E-02 | 0.70 | 1.67 | 1.03 |
| *Ahsg* | Endocard | 2.03E-05 | 2.43E-02 | 0.63 | 2.51 | 1.62 |
| *Hbb-bs* | Endocard | 3.07E-05 | 3.21E-02 | 2.68 | 0.46 | 0.07 |
| *G3bp2* | Endocard | 4.58E-05 | 4.26E-02 | -0.48 | 1.85 | 2.59 |
| *Fmo2* | Endocard | 5.89E-05 | 4.80E-02 | 0.12 | 1.95 | 1.79 |
| *Hsp90aa1* | Endothelial cells | 7.46E-147 | 4.69E-143 | 1.06 | 5.21 | 2.50 |
| *Hspa1a* | Endothelial cells | 2.32E-100 | 7.29E-97 | 0.69 | 13.53 | 8.41 |
| *Dnaja1* | Endothelial cells | 2.60E-88 | 5.45E-85 | 0.49 | 13.07 | 9.34 |
| *Xist* | Endothelial cells | 7.26E-88 | 1.14E-84 | -0.62 | 3.77 | 5.80 |
| *Hsph1* | Endothelial cells | 1.33E-87 | 1.56E-84 | 1.50 | 0.80 | 0.28 |
| *Hspa1b* | Endothelial cells | 1.49E-87 | 1.56E-84 | 0.65 | 14.33 | 9.11 |
| *Hspa8* | Endothelial cells | 3.65E-81 | 3.27E-78 | 0.34 | 17.35 | 13.68 |
| *Cacybp* | Endothelial cells | 5.68E-63 | 4.46E-60 | 0.70 | 1.90 | 1.17 |
| *Gm47283* | Endothelial cells | 6.96E-63 | 4.86E-60 | -0.76 | 1.05 | 1.77 |
| *Lyz2* | Endothelial cells | 1.82E-58 | 1.14E-55 | 1.28 | 1.19 | 0.49 |
| *Manf* | Fibroblasts | 4.39E-106 | 3.81E-102 | -0.68 | 2.42 | 3.89 |
| *Angptl4* | Fibroblasts | 1.26E-99 | 5.46E-96 | 1.42 | 0.63 | 0.23 |
| *AY036118* | Fibroblasts | 6.10E-83 | 1.52E-79 | 0.41 | 5.67 | 4.26 |
| *Hspb1* | Fibroblasts | 7.01E-83 | 1.52E-79 | 0.82 | 10.17 | 5.77 |
| *Hspa1b* | Fibroblasts | 1.93E-82 | 3.34E-79 | 0.89 | 5.66 | 3.07 |
| *Pdia6* | Fibroblasts | 5.97E-78 | 8.64E-75 | -0.41 | 3.40 | 4.53 |
| *Dnaja1* | Fibroblasts | 7.34E-75 | 9.10E-72 | 0.43 | 6.35 | 4.73 |
| *Gm47283* | Fibroblasts | 4.13E-70 | 4.48E-67 | -0.67 | 0.93 | 1.49 |
| *Hspa1a* | Fibroblasts | 2.12E-68 | 2.05E-65 | 0.90 | 4.23 | 2.27 |
| *Rbm3* | Fibroblasts | 2.77E-67 | 2.41E-64 | 0.41 | 1.88 | 1.41 |
| *AY036118* | Macrophages | 3.92E-50 | 2.86E-46 | 0.63 | 4.08 | 2.63 |
| *Xist* | Macrophages | 1.62E-40 | 5.89E-37 | -0.58 | 5.70 | 8.49 |
| *Dnaja1* | Macrophages | 1.66E-38 | 3.78E-35 | 0.64 | 11.72 | 7.54 |
| *Hsph1* | Macrophages | 2.07E-38 | 3.78E-35 | 1.14 | 1.69 | 0.77 |
| *Rbm3* | Macrophages | 6.43E-33 | 9.38E-30 | 0.46 | 2.26 | 1.64 |
| *Plin2* | Macrophages | 2.82E-32 | 3.43E-29 | 0.56 | 2.88 | 1.95 |
| *Gm47283* | Macrophages | 1.74E-28 | 1.81E-25 | -0.89 | 0.54 | 1.00 |
| *Hspa8* | Macrophages | 5.75E-28 | 5.24E-25 | 0.43 | 28.64 | 21.27 |
| *Hspa1b* | Macrophages | 3.18E-25 | 2.57E-22 | 0.55 | 26.97 | 18.41 |
| *Dnajb1* | Macrophages | 7.67E-25 | 5.59E-22 | 0.82 | 5.21 | 2.95 |
| *Dnaja1* | NK cells | 3.89E-09 | 2.04E-05 | 1.19 | 16.87 | 7.39 |
| *Hspa1b* | NK cells | 1.69E-06 | 4.45E-03 | 2.22 | 10.37 | 2.23 |
| *Hsph1* | NK cells | 5.01E-06 | 8.78E-03 | 2.26 | 3.88 | 0.81 |
| *Hsp90aa1* | NK cells | 7.13E-06 | 9.37E-03 | 1.21 | 15.46 | 6.70 |
| *Dnajb1* | NK cells | 9.04E-06 | 9.51E-03 | 1.61 | 4.60 | 1.51 |
| *Hsp90ab1* | NK cells | 1.19E-05 | 1.04E-02 | 0.80 | 27.46 | 15.75 |
| *Hbb-bs* | NK cells | 3.78E-05 | 2.42E-02 | 4.63 | 0.84 | 0.03 |
| *Rplp0* | NK cells | 3.80E-05 | 2.42E-02 | -0.56 | 15.97 | 23.53 |
| *Pdcl3* | NK cells | 4.14E-05 | 2.42E-02 | -1.03 | 0.41 | 0.83 |
| *Pfn1* | NK cells | 5.19E-05 | 2.73E-02 | -0.74 | 16.07 | 26.92 |
| *Dnaja1* | Pericytes | 1.92E-22 | 1.20E-18 | 0.67 | 11.62 | 7.28 |
| *Hspa8* | Pericytes | 8.21E-16 | 2.56E-12 | 0.31 | 14.43 | 11.65 |
| *Hsp90aa1* | Pericytes | 8.84E-15 | 1.84E-11 | 0.74 | 6.40 | 3.84 |
| *Ech1* | Pericytes | 1.22E-14 | 1.90E-11 | 0.68 | 2.25 | 1.40 |
| *Hspa1b* | Pericytes | 3.07E-14 | 3.83E-11 | 0.85 | 10.29 | 5.72 |
| *Hspa1a* | Pericytes | 2.91E-12 | 3.03E-09 | 0.72 | 9.54 | 5.78 |
| *Gm47283* | Pericytes | 1.19E-10 | 1.06E-07 | -0.61 | 1.89 | 2.88 |
| *Xist* | Pericytes | 2.03E-09 | 1.59E-06 | -0.29 | 7.84 | 9.62 |
| *Lyz2* | Pericytes | 2.39E-09 | 1.66E-06 | 1.65 | 1.37 | 0.44 |
| *Hbb-bs* | Pericytes | 3.36E-09 | 2.10E-06 | 1.80 | 0.70 | 0.20 |
| *Plod2* | Pro.cells | 5.51E-06 | 4.16E-02 | 1.27 | 0.50 | 0.21 |
| *C1qc* | Pro.cells | 7.76E-06 | 4.16E-02 | -1.25 | 2.58 | 6.14 |
| *Sfrp5* | Schwann cells | 3.57E-10 | 2.65E-06 |  | 0.89 | 0.00 |
| *Gm47283* | Schwann cells | 4.38E-09 | 1.63E-05 | -1.82 | 1.20 | 4.27 |
| *Hspa1a* | Schwann cells | 6.23E-06 | 1.54E-02 | 1.53 | 6.59 | 2.28 |
| *Tle1* | Schwann cells | 6.07E-05 | 1.13E-01 | -2.40 | 0.18 | 0.96 |
| *Lyz2* | Schwann cells | 1.06E-04 | 1.58E-01 | -0.90 | 0.99 | 1.85 |
| *Ppp1r12b* | Schwann cells | 1.60E-04 | 1.94E-01 | -1.29 | 0.88 | 2.16 |
| *Igkc* | Schwann cells | 1.86E-04 | 1.94E-01 |  | 0.42 | 0.00 |
| *Etfb* | Schwann cells | 2.27E-04 | 1.94E-01 | 1.25 | 1.30 | 0.54 |
| *Rbm3* | Schwann cells | 2.35E-04 | 1.94E-01 | 1.17 | 1.84 | 0.82 |
| *Aak1* | Schwann cells | 2.90E-04 | 2.15E-01 | -1.27 | 0.20 | 0.48 |
| *Atp1b2* | SMC | 3.54E-07 | 2.89E-03 | -0.32 | 3.41 | 4.26 |
| *Hspa1a* | SMC | 7.93E-07 | 3.23E-03 | 0.52 | 7.93 | 5.54 |
| *Lyz2* | SMC | 2.07E-05 | 4.91E-02 | 2.15 | 1.25 | 0.28 |
| *Atp6v0a2* | SMC | 2.41E-05 | 4.91E-02 | 0.60 | 0.35 | 0.23 |
| *Hsp90aa1* | T cells | 2.95E-21 | 1.49E-17 | 1.23 | 13.60 | 5.81 |
| *Dnaja1* | T cells | 8.45E-13 | 2.14E-09 | 0.68 | 13.88 | 8.64 |
| *Hspa1b* | T cells | 1.49E-11 | 2.06E-08 | 1.75 | 7.37 | 2.18 |
| *Hspa1a* | T cells | 1.62E-11 | 2.06E-08 | 2.02 | 6.05 | 1.49 |
| *H2-K1* | T cells | 2.07E-11 | 2.10E-08 | 0.54 | 30.74 | 21.21 |
| *Dnajb1* | T cells | 3.75E-11 | 3.17E-08 | 1.48 | 4.98 | 1.78 |
| *Gm47283* | T cells | 2.48E-10 | 1.79E-07 | -0.89 | 1.72 | 3.18 |
| *Hsph1* | T cells | 7.27E-09 | 4.61E-06 | 1.16 | 2.50 | 1.12 |
| *Xist* | T cells | 5.49E-08 | 3.09E-05 | -1.05 | 6.00 | 12.42 |
| *H2-D1* | T cells | 2.23E-07 | 1.13E-04 | 0.48 | 23.84 | 17.05 |

Legend: NK, natural killer cells; SMC, smooth muscle cells.

**Supplementary Table 6.** Gene ontology summary table of up- and down-regulated pathways**.**

| ID | Description | GeneRatio | Adjusted P-value | Cell type | Up/Down-regulated in high fibre |
| --- | --- | --- | --- | --- | --- |
| GO:0043062 | extracellular structure organization | 52/754 | 5.50106E-13 | Fibroblasts | Down-regulated |
| GO:0031589 | cell-substrate adhesion | 48/754 | 6.17204E-09 | Fibroblasts | Down-regulated |
| GO:0034446 | substrate adhesion-dependent cell spreading | 24/754 | 8.45664E-08 | Fibroblasts | Down-regulated |
| GO:0032970 | regulation of actin filament-based process | 47/754 | 5.3046E-06 | Fibroblasts | Down-regulated |
| GO:0022604 | regulation of cell morphogenesis | 55/754 | 5.83482E-06 | Fibroblasts | Down-regulated |
| GO:0035456 | response to interferon-beta | 14/511 | 1.16391E-05 | Endothelial cells | Down-regulated |
| GO:0007015 | actin filament organization | 45/754 | 3.67147E-05 | Fibroblasts | Down-regulated |
| GO:0032956 | regulation of actin cytoskeleton organization | 40/754 | 8.41395E-05 | Fibroblasts | Down-regulated |
| GO:0061448 | connective tissue development | 32/754 | 8.41395E-05 | Fibroblasts | Down-regulated |
| GO:0034976 | response to endoplasmic reticulum stress | 31/754 | 9.7657E-05 | Fibroblasts | Down-regulated |
| GO:0001501 | skeletal system development | 48/754 | 0.000120798 | Fibroblasts | Down-regulated |
| GO:0006888 | ER to Golgi vesicle-mediated transport | 20/754 | 0.000572564 | Fibroblasts | Down-regulated |
| GO:0071230 | cellular response to amino acid stimulus | 14/754 | 0.000572564 | Fibroblasts | Down-regulated |
| GO:0010810 | regulation of cell-substrate adhesion | 27/754 | 0.000572564 | Fibroblasts | Down-regulated |
| GO:0032527 | protein exit from endoplasmic reticulum | 11/754 | 0.000582699 | Fibroblasts | Down-regulated |
| GO:0007178 | transmembrane receptor protein serine/threonine kinase signaling pathway | 34/754 | 0.000582699 | Fibroblasts | Down-regulated |
| GO:0043200 | response to amino acid | 15/754 | 0.000582699 | Fibroblasts | Down-regulated |
| GO:0051607 | defense response to virus | 23/511 | 0.00067799 | Endothelial cells | Down-regulated |
| GO:0050792 | regulation of viral process | 20/511 | 0.000713551 | Endothelial cells | Down-regulated |
| GO:0001765 | membrane raft assembly | 6/511 | 0.000713551 | Endothelial cells | Down-regulated |
| GO:0072657 | protein localization to membrane | 49/754 | 0.000881937 | Fibroblasts | Down-regulated |
| GO:0001525 | angiogenesis | 47/754 | 0.000916534 | Fibroblasts | Down-regulated |
| GO:0009611 | response to wounding | 43/754 | 0.001039934 | Fibroblasts | Down-regulated |
| GO:0070208 | protein heterotrimerization | 7/754 | 0.00110933 | Fibroblasts | Down-regulated |
| GO:0051489 | regulation of filopodium assembly | 11/754 | 0.00110933 | Fibroblasts | Down-regulated |
| GO:0044403 | symbiont process | 29/511 | 0.001187652 | Endothelial cells | Down-regulated |
| GO:0035458 | cellular response to interferon-beta | 10/511 | 0.001228967 | Endothelial cells | Down-regulated |
| GO:0000280 | nuclear division | 12/94 | 0.001569694 | Pro.cells | Down-regulated |
| GO:0140014 | mitotic nuclear division | 10/94 | 0.001569694 | Pro.cells | Down-regulated |
| GO:0000910 | cytokinesis | 8/94 | 0.001569694 | Pro.cells | Down-regulated |
| GO:0032506 | cytokinetic process | 5/94 | 0.001569694 | Pro.cells | Down-regulated |
| GO:0032886 | regulation of microtubule-based process | 9/94 | 0.002296492 | Pro.cells | Down-regulated |
| GO:0016032 | viral process | 24/511 | 0.002444112 | Endothelial cells | Down-regulated |
| GO:0007059 | chromosome segregation | 10/94 | 0.002668261 | Pro.cells | Down-regulated |
| GO:0090068 | positive regulation of cell cycle process | 9/94 | 0.002668261 | Pro.cells | Down-regulated |
| GO:0001503 | ossification | 35/754 | 0.002712881 | Fibroblasts | Down-regulated |
| GO:0051271 | negative regulation of cellular component movement | 32/754 | 0.002739218 | Fibroblasts | Down-regulated |
| GO:1904951 | positive regulation of establishment of protein localization | 40/754 | 0.002772963 | Fibroblasts | Down-regulated |
| GO:0030970 | retrograde protein transport, ER to cytosol | 8/754 | 0.003078995 | Fibroblasts | Down-regulated |
| GO:1903513 | endoplasmic reticulum to cytosol transport | 8/754 | 0.003078995 | Fibroblasts | Down-regulated |
| GO:0035265 | organ growth | 24/754 | 0.003115417 | Fibroblasts | Down-regulated |
| GO:0090287 | regulation of cellular response to growth factor stimulus | 28/754 | 0.003301385 | Fibroblasts | Down-regulated |
| GO:0018279 | protein N-linked glycosylation via asparagine | 7/754 | 0.003398822 | Fibroblasts | Down-regulated |
| GO:0044706 | multi-multicellular organism process | 18/754 | 0.003555565 | Fibroblasts | Down-regulated |
| GO:0070507 | regulation of microtubule cytoskeleton organization | 8/94 | 0.003672505 | Pro.cells | Down-regulated |
| GO:0007565 | female pregnancy | 16/754 | 0.004073673 | Fibroblasts | Down-regulated |
| GO:0040013 | negative regulation of locomotion | 32/754 | 0.004073673 | Fibroblasts | Down-regulated |
| GO:0018196 | peptidyl-asparagine modification | 7/754 | 0.004177986 | Fibroblasts | Down-regulated |
| GO:0016126 | sterol biosynthetic process | 10/754 | 0.004307511 | Fibroblasts | Down-regulated |
| GO:0035437 | maintenance of protein localization in endoplasmic reticulum | 5/754 | 0.005085411 | Fibroblasts | Down-regulated |
| GO:0031667 | response to nutrient levels | 20/318 | 0.005214973 | Macrophages | Down-regulated |
| GO:0009617 | response to bacterium | 28/318 | 0.005214973 | Macrophages | Down-regulated |
| GO:0009991 | response to extracellular stimulus | 21/318 | 0.005214973 | Macrophages | Down-regulated |
| GO:0043254 | regulation of protein complex assembly | 32/511 | 0.005671314 | Endothelial cells | Down-regulated |
| GO:0001570 | vasculogenesis | 12/511 | 0.005671314 | Endothelial cells | Down-regulated |
| GO:0032963 | collagen metabolic process | 15/754 | 0.006080858 | Fibroblasts | Down-regulated |
| GO:1905244 | regulation of modification of synaptic structure | 6/754 | 0.006080858 | Fibroblasts | Down-regulated |
| GO:0090218 | positive regulation of lipid kinase activity | 8/754 | 0.00623736 | Fibroblasts | Down-regulated |
| GO:0110053 | regulation of actin filament organization | 27/754 | 0.00634205 | Fibroblasts | Down-regulated |
| GO:0045765 | regulation of angiogenesis | 24/511 | 0.007214666 | Endothelial cells | Down-regulated |
| GO:0120032 | regulation of plasma membrane bounded cell projection assembly | 21/754 | 0.007495136 | Fibroblasts | Down-regulated |
| GO:0050708 | regulation of protein secretion | 39/754 | 0.007526383 | Fibroblasts | Down-regulated |
| GO:0099010 | modification of postsynaptic structure | 7/754 | 0.008250984 | Fibroblasts | Down-regulated |
| GO:0070059 | intrinsic apoptotic signaling pathway in response to endoplasmic reticulum stress | 11/754 | 0.00825758 | Fibroblasts | Down-regulated |
| GO:0045834 | positive regulation of lipid metabolic process | 18/754 | 0.008932824 | Fibroblasts | Down-regulated |
| GO:0035966 | response to topologically incorrect protein | 17/754 | 0.009387897 | Fibroblasts | Down-regulated |
| GO:0000819 | sister chromatid segregation | 7/94 | 0.009558649 | Pro.cells | Down-regulated |
| GO:0090257 | regulation of muscle system process | 24/754 | 0.010047212 | Fibroblasts | Down-regulated |
| GO:0071229 | cellular response to acid chemical | 19/754 | 0.010047212 | Fibroblasts | Down-regulated |
| GO:1905897 | regulation of response to endoplasmic reticulum stress | 12/754 | 0.010189777 | Fibroblasts | Down-regulated |
| GO:0099159 | regulation of modification of postsynaptic structure | 5/754 | 0.010189777 | Fibroblasts | Down-regulated |
| GO:0071496 | cellular response to external stimulus | 25/754 | 0.011503933 | Fibroblasts | Down-regulated |
| GO:0001101 | response to acid chemical | 24/754 | 0.011503933 | Fibroblasts | Down-regulated |
| GO:0002791 | regulation of peptide secretion | 40/754 | 0.01187935 | Fibroblasts | Down-regulated |
| GO:0006900 | vesicle budding from membrane | 10/754 | 0.012207923 | Fibroblasts | Down-regulated |
| GO:0051345 | positive regulation of hydrolase activity | 42/754 | 0.012330995 | Fibroblasts | Down-regulated |
| GO:0035967 | cellular response to topologically incorrect protein | 14/754 | 0.012936955 | Fibroblasts | Down-regulated |
| GO:0001649 | osteoblast differentiation | 20/754 | 0.013073136 | Fibroblasts | Down-regulated |
| GO:0045862 | positive regulation of proteolysis | 30/754 | 0.013772303 | Fibroblasts | Down-regulated |
| GO:0006457 | protein folding | 18/754 | 0.013772303 | Fibroblasts | Down-regulated |
| GO:0002456 | T cell mediated immunity | 11/318 | 0.013850746 | Macrophages | Down-regulated |
| GO:0006875 | cellular metal ion homeostasis | 25/318 | 0.013850746 | Macrophages | Down-regulated |
| GO:0002709 | regulation of T cell mediated immunity | 9/318 | 0.014399114 | Macrophages | Down-regulated |
| GO:0031579 | membrane raft organization | 6/754 | 0.014605655 | Fibroblasts | Down-regulated |
| GO:0032611 | interleukin-1 beta production | 9/318 | 0.014608075 | Macrophages | Down-regulated |
| GO:0030316 | osteoclast differentiation | 10/318 | 0.014608075 | Macrophages | Down-regulated |
| GO:0031333 | negative regulation of protein complex assembly | 17/754 | 0.015864993 | Fibroblasts | Down-regulated |
| GO:0044091 | membrane biogenesis | 8/754 | 0.017560704 | Fibroblasts | Down-regulated |
| GO:0030968 | endoplasmic reticulum unfolded protein response | 10/754 | 0.017888793 | Fibroblasts | Down-regulated |
| GO:0120034 | positive regulation of plasma membrane bounded cell projection assembly | 14/754 | 0.018172893 | Fibroblasts | Down-regulated |
| GO:0042113 | B cell activation | 9/85 | 0.018421664 | B cells | Down-regulated |
| GO:0034341 | response to interferon-gamma | 14/511 | 0.01846068 | Endothelial cells | Down-regulated |
| GO:0071407 | cellular response to organic cyclic compound | 20/318 | 0.019788809 | Macrophages | Down-regulated |
| GO:0048002 | antigen processing and presentation of peptide antigen | 7/318 | 0.019788809 | Macrophages | Down-regulated |
| GO:0031668 | cellular response to extracellular stimulus | 14/318 | 0.019788809 | Macrophages | Down-regulated |
| GO:0006826 | iron ion transport | 6/318 | 0.019788809 | Macrophages | Down-regulated |
| GO:0032535 | regulation of cellular component size | 33/754 | 0.021247863 | Fibroblasts | Down-regulated |
| GO:1901216 | positive regulation of neuron death | 15/754 | 0.021720166 | Fibroblasts | Down-regulated |
| GO:0010761 | fibroblast migration | 9/754 | 0.021794913 | Fibroblasts | Down-regulated |
| GO:0098885 | modification of postsynaptic actin cytoskeleton | 5/754 | 0.022573919 | Fibroblasts | Down-regulated |
| GO:0007088 | regulation of mitotic nuclear division | 6/94 | 0.022737953 | Pro.cells | Down-regulated |
| GO:0071496 | cellular response to external stimulus | 15/318 | 0.022948023 | Macrophages | Down-regulated |
| GO:0006898 | receptor-mediated endocytosis | 14/318 | 0.022948023 | Macrophages | Down-regulated |
| GO:0009896 | positive regulation of catabolic process | 21/318 | 0.022948023 | Macrophages | Down-regulated |
| GO:0071559 | response to transforming growth factor beta | 21/754 | 0.024161729 | Fibroblasts | Down-regulated |
| GO:0070972 | protein localization to endoplasmic reticulum | 10/754 | 0.024458984 | Fibroblasts | Down-regulated |
| GO:0007030 | Golgi organization | 15/754 | 0.024737206 | Fibroblasts | Down-regulated |
| GO:0007033 | vacuole organization | 12/318 | 0.025210598 | Macrophages | Down-regulated |
| GO:0035774 | positive regulation of insulin secretion involved in cellular response to glucose stimulus | 7/754 | 0.025717643 | Fibroblasts | Down-regulated |
| GO:0051668 | localization within membrane | 19/754 | 0.026559301 | Fibroblasts | Down-regulated |
| GO:0032387 | negative regulation of intracellular transport | 10/754 | 0.026707593 | Fibroblasts | Down-regulated |
| GO:1903827 | regulation of cellular protein localization | 41/754 | 0.026707593 | Fibroblasts | Down-regulated |
| GO:1905475 | regulation of protein localization to membrane | 19/754 | 0.029307287 | Fibroblasts | Down-regulated |
| GO:0006970 | response to osmotic stress | 10/754 | 0.029307287 | Fibroblasts | Down-regulated |
| GO:0071470 | cellular response to osmotic stress | 7/754 | 0.029307287 | Fibroblasts | Down-regulated |
| GO:0008299 | isoprenoid biosynthetic process | 6/754 | 0.031729256 | Fibroblasts | Down-regulated |
| GO:0104004 | cellular response to environmental stimulus | 23/754 | 0.033433208 | Fibroblasts | Down-regulated |
| GO:0045672 | positive regulation of osteoclast differentiation | 5/318 | 0.034210166 | Macrophages | Down-regulated |
| GO:0045669 | positive regulation of osteoblast differentiation | 10/754 | 0.034255387 | Fibroblasts | Down-regulated |
| GO:0045666 | positive regulation of neuron differentiation | 35/754 | 0.034255387 | Fibroblasts | Down-regulated |
| GO:0098760 | response to interleukin-7 | 5/754 | 0.034255387 | Fibroblasts | Down-regulated |
| GO:0098761 | cellular response to interleukin-7 | 5/754 | 0.034255387 | Fibroblasts | Down-regulated |
| GO:1901699 | cellular response to nitrogen compound | 39/754 | 0.03545878 | Fibroblasts | Down-regulated |
| GO:0016125 | sterol metabolic process | 14/754 | 0.036978452 | Fibroblasts | Down-regulated |
| GO:0071396 | cellular response to lipid | 35/754 | 0.036978452 | Fibroblasts | Down-regulated |
| GO:0055074 | calcium ion homeostasis | 20/318 | 0.038157129 | Macrophages | Down-regulated |
| GO:0006935 | chemotaxis | 23/318 | 0.038157129 | Macrophages | Down-regulated |
| GO:1900038 | negative regulation of cellular response to hypoxia | 4/511 | 0.038573127 | Endothelial cells | Down-regulated |
| GO:0045453 | bone resorption | 7/318 | 0.039267848 | Macrophages | Down-regulated |
| GO:0050729 | positive regulation of inflammatory response | 10/318 | 0.039267848 | Macrophages | Down-regulated |
| GO:0045185 | maintenance of protein location | 13/754 | 0.039729393 | Fibroblasts | Down-regulated |
| GO:1903034 | regulation of response to wounding | 17/754 | 0.04037888 | Fibroblasts | Down-regulated |
| GO:0051781 | positive regulation of cell division | 4/94 | 0.04057666 | Pro.cells | Down-regulated |
| GO:0031345 | negative regulation of cell projection organization | 17/511 | 0.04068917 | Endothelial cells | Down-regulated |
| GO:0034329 | cell junction assembly | 17/511 | 0.04074593 | Endothelial cells | Down-regulated |
| GO:0047484 | regulation of response to osmotic stress | 4/754 | 0.041173859 | Fibroblasts | Down-regulated |
| GO:0060394 | negative regulation of pathway-restricted SMAD protein phosphorylation | 4/754 | 0.041173859 | Fibroblasts | Down-regulated |
| GO:0032651 | regulation of interleukin-1 beta production | 7/318 | 0.042557995 | Macrophages | Down-regulated |
| GO:0046849 | bone remodeling | 8/318 | 0.042557995 | Macrophages | Down-regulated |
| GO:0090066 | regulation of anatomical structure size | 40/754 | 0.042697464 | Fibroblasts | Down-regulated |
| GO:0061640 | cytoskeleton-dependent cytokinesis | 12/754 | 0.043138032 | Fibroblasts | Down-regulated |
| GO:0030855 | epithelial cell differentiation | 31/511 | 0.043365694 | Endothelial cells | Down-regulated |
| GO:0140253 | cell-cell fusion | 9/754 | 0.044219598 | Fibroblasts | Down-regulated |
| GO:0090162 | establishment of epithelial cell polarity | 6/511 | 0.046595264 | Endothelial cells | Down-regulated |
| GO:0031333 | negative regulation of protein complex assembly | 10/318 | 0.048692003 | Macrophages | Down-regulated |
| GO:0097191 | extrinsic apoptotic signaling pathway | 13/318 | 0.048817323 | Macrophages | Down-regulated |
| GO:0044089 | positive regulation of cellular component biogenesis | 40/754 | 0.049205954 | Fibroblasts | Down-regulated |
| GO:0042026 | protein refolding | 2/33 | 0.314202405 | Schwann cells | Down-regulated |
| GO:0051085 | chaperone cofactor-dependent protein refolding | 2/33 | 0.314202405 | Schwann cells | Down-regulated |
| GO:0044839 | cell cycle G2/M phase transition | 3/33 | 0.314202405 | Schwann cells | Down-regulated |
| GO:0051084 | 'de novo' posttranslational protein folding | 2/33 | 0.314202405 | Schwann cells | Down-regulated |
| GO:0006458 | 'de novo' protein folding | 2/33 | 0.314202405 | Schwann cells | Down-regulated |
| GO:0061077 | chaperone-mediated protein folding | 2/33 | 0.314202405 | Schwann cells | Down-regulated |
| GO:1902750 | negative regulation of cell cycle G2/M phase transition | 2/33 | 0.314202405 | Schwann cells | Down-regulated |
| GO:0034605 | cellular response to heat | 2/33 | 0.314202405 | Schwann cells | Down-regulated |
| GO:0050830 | defense response to Gram-positive bacterium | 2/33 | 0.314202405 | Schwann cells | Down-regulated |
| GO:0032507 | maintenance of protein location in cell | 2/33 | 0.314202405 | Schwann cells | Down-regulated |
| GO:0034620 | cellular response to unfolded protein | 2/33 | 0.314202405 | Schwann cells | Down-regulated |
| GO:0051651 | maintenance of location in cell | 2/33 | 0.314202405 | Schwann cells | Down-regulated |
| GO:1902749 | regulation of cell cycle G2/M phase transition | 2/33 | 0.314202405 | Schwann cells | Down-regulated |
| GO:0090101 | negative regulation of transmembrane receptor protein serine/threonine kinase signaling pathway | 2/33 | 0.314202405 | Schwann cells | Down-regulated |
| GO:0035967 | cellular response to topologically incorrect protein | 2/33 | 0.314202405 | Schwann cells | Down-regulated |
| GO:0009408 | response to heat | 2/33 | 0.314202405 | Schwann cells | Down-regulated |
| GO:0045185 | maintenance of protein location | 2/33 | 0.314202405 | Schwann cells | Down-regulated |
| GO:0006986 | response to unfolded protein | 2/33 | 0.314202405 | Schwann cells | Down-regulated |
| GO:0006913 | nucleocytoplasmic transport | 3/33 | 0.314202405 | Schwann cells | Down-regulated |
| GO:0051169 | nuclear transport | 3/33 | 0.314202405 | Schwann cells | Down-regulated |
| GO:0071453 | cellular response to oxygen levels | 2/33 | 0.314202405 | Schwann cells | Down-regulated |
| GO:0000086 | G2/M transition of mitotic cell cycle | 2/33 | 0.314202405 | Schwann cells | Down-regulated |
| GO:0006611 | protein export from nucleus | 2/33 | 0.314202405 | Schwann cells | Down-regulated |
| GO:0035966 | response to topologically incorrect protein | 2/33 | 0.314202405 | Schwann cells | Down-regulated |
| GO:0009267 | cellular response to starvation | 2/33 | 0.314202405 | Schwann cells | Down-regulated |
| GO:0051168 | nuclear export | 2/33 | 0.314202405 | Schwann cells | Down-regulated |
| GO:0030902 | hindbrain development | 2/33 | 0.314202405 | Schwann cells | Down-regulated |
| GO:0031333 | negative regulation of protein complex assembly | 2/33 | 0.314202405 | Schwann cells | Down-regulated |
| GO:0006457 | protein folding | 2/33 | 0.314202405 | Schwann cells | Down-regulated |
| GO:0009266 | response to temperature stimulus | 2/33 | 0.314202405 | Schwann cells | Down-regulated |
| GO:0042594 | response to starvation | 2/33 | 0.314202405 | Schwann cells | Down-regulated |
| GO:1901988 | negative regulation of cell cycle phase transition | 2/33 | 0.314202405 | Schwann cells | Down-regulated |
| GO:0007033 | vacuole organization | 2/33 | 0.333059899 | Schwann cells | Down-regulated |
| GO:0022613 | ribonucleoprotein complex biogenesis | 74/862 | 1.44988E-15 | Fibroblasts | Up-regulated |
| GO:0006457 | protein folding | 17/100 | 1.50762E-13 | B cells | Up-regulated |
| GO:0006457 | protein folding | 22/237 | 2.42258E-12 | Macrophages | Up-regulated |
| GO:0006457 | protein folding | 29/535 | 1.06586E-10 | Endothelial cells | Up-regulated |
| GO:0006457 | protein folding | 14/87 | 1.87379E-10 | Pericytes | Up-regulated |
| GO:0006457 | protein folding | 11/43 | 2.38356E-10 | NK cells | Up-regulated |
| GO:0002181 | cytoplasmic translation | 25/862 | 3.88824E-09 | Fibroblasts | Up-regulated |
| GO:0016072 | rRNA metabolic process | 41/862 | 3.8946E-09 | Fibroblasts | Up-regulated |
| GO:0006457 | protein folding | 12/71 | 5.53301E-09 | T cells | Up-regulated |
| GO:0043484 | regulation of RNA splicing | 30/862 | 1.59646E-08 | Fibroblasts | Up-regulated |
| GO:0061077 | chaperone-mediated protein folding | 7/39 | 4.59025E-08 | DC-like cells | Up-regulated |
| GO:0006457 | protein folding | 9/39 | 4.59025E-08 | DC-like cells | Up-regulated |
| GO:0051085 | chaperone cofactor-dependent protein refolding | 6/39 | 4.59025E-08 | DC-like cells | Up-regulated |
| GO:0006986 | response to unfolded protein | 15/237 | 5.00461E-08 | Macrophages | Up-regulated |
| GO:0006986 | response to unfolded protein | 8/39 | 5.00684E-08 | DC-like cells | Up-regulated |
| GO:0051085 | chaperone cofactor-dependent protein refolding | 12/535 | 2.43834E-07 | Endothelial cells | Up-regulated |
| GO:0022618 | ribonucleoprotein complex assembly | 37/862 | 3.72983E-07 | Fibroblasts | Up-regulated |
| GO:0008380 | RNA splicing | 38/535 | 4.26766E-07 | Endothelial cells | Up-regulated |
| GO:0006986 | response to unfolded protein | 10/100 | 4.84822E-07 | B cells | Up-regulated |
| GO:0071826 | ribonucleoprotein complex subunit organization | 38/862 | 5.54016E-07 | Fibroblasts | Up-regulated |
| GO:0009408 | response to heat | 7/39 | 6.34975E-07 | DC-like cells | Up-regulated |
| GO:0002181 | cytoplasmic translation | 9/100 | 6.70014E-06 | Endocardium | Up-regulated |
| GO:0006417 | regulation of translation | 43/862 | 1.05244E-05 | Fibroblasts | Up-regulated |
| GO:0034248 | regulation of cellular amide metabolic process | 47/862 | 1.05244E-05 | Fibroblasts | Up-regulated |
| GO:0043484 | regulation of RNA splicing | 20/535 | 1.09183E-05 | Endothelial cells | Up-regulated |
| GO:0010608 | posttranscriptional regulation of gene expression | 52/862 | 1.15423E-05 | Fibroblasts | Up-regulated |
| GO:0042326 | negative regulation of phosphorylation | 53/862 | 1.15999E-05 | Fibroblasts | Up-regulated |
| GO:0051085 | chaperone cofactor-dependent protein refolding | 6/100 | 3.97943E-05 | Endocardium | Up-regulated |
| GO:0043062 | extracellular structure organization | 14/119 | 6.17027E-05 | Pro.cells | Up-regulated |
| GO:0030198 | extracellular matrix organization | 13/119 | 6.17027E-05 | Pro.cells | Up-regulated |
| GO:0001667 | ameboidal-type cell migration | 35/535 | 7.93193E-05 | Endothelial cells | Up-regulated |
| GO:0030099 | myeloid cell differentiation | 34/535 | 7.93193E-05 | Endothelial cells | Up-regulated |
| GO:0006457 | protein folding | 7/40 | 9.7623E-05 | Schwann cells | Up-regulated |
| GO:0034620 | cellular response to unfolded protein | 5/39 | 0.000115676 | DC-like cells | Up-regulated |
| GO:2001233 | regulation of apoptotic signaling pathway | 47/862 | 0.000143319 | Fibroblasts | Up-regulated |
| GO:0051348 | negative regulation of transferase activity | 33/862 | 0.000179624 | Fibroblasts | Up-regulated |
| GO:0009266 | response to temperature stimulus | 13/237 | 0.000203603 | Macrophages | Up-regulated |
| GO:0034504 | protein localization to nucleus | 35/862 | 0.000210664 | Fibroblasts | Up-regulated |
| GO:0006457 | protein folding | 24/862 | 0.000323062 | Fibroblasts | Up-regulated |
| GO:0060249 | anatomical structure homeostasis | 31/535 | 0.000467713 | Endothelial cells | Up-regulated |
| GO:0055094 | response to lipoprotein particle | 8/535 | 0.000467713 | Endothelial cells | Up-regulated |
| GO:2000573 | positive regulation of DNA biosynthetic process | 9/237 | 0.00048551 | Macrophages | Up-regulated |
| GO:0009266 | response to temperature stimulus | 8/87 | 0.000550251 | Pericytes | Up-regulated |
| GO:0006986 | response to unfolded protein | 7/87 | 0.000550251 | Pericytes | Up-regulated |
| GO:0060537 | muscle tissue development | 47/862 | 0.000626315 | Fibroblasts | Up-regulated |
| GO:1903828 | negative regulation of cellular protein localization | 5/39 | 0.00062782 | DC-like cells | Up-regulated |
| GO:0090132 | epithelium migration | 26/535 | 0.000637773 | Endothelial cells | Up-regulated |
| GO:0090130 | tissue migration | 26/535 | 0.000637773 | Endothelial cells | Up-regulated |
| GO:0071402 | cellular response to lipoprotein particle stimulus | 8/535 | 0.000637773 | Endothelial cells | Up-regulated |
| GO:0034620 | cellular response to unfolded protein | 9/237 | 0.000707943 | Macrophages | Up-regulated |
| GO:0048511 | rhythmic process | 32/862 | 0.000759477 | Fibroblasts | Up-regulated |
| GO:0009266 | response to temperature stimulus | 18/535 | 0.000800883 | Endothelial cells | Up-regulated |
| GO:0010594 | regulation of endothelial cell migration | 18/535 | 0.000800883 | Endothelial cells | Up-regulated |
| GO:1901343 | negative regulation of vasculature development | 16/535 | 0.0008065 | Endothelial cells | Up-regulated |
| GO:0034605 | cellular response to heat | 6/100 | 0.000819945 | B cells | Up-regulated |
| GO:0009408 | response to heat | 7/100 | 0.000819945 | B cells | Up-regulated |
| GO:1903707 | negative regulation of hemopoiesis | 18/535 | 0.000821051 | Endothelial cells | Up-regulated |
| GO:0001933 | negative regulation of protein phosphorylation | 44/862 | 0.000832073 | Fibroblasts | Up-regulated |
| GO:0001525 | angiogenesis | 37/535 | 0.000832635 | Endothelial cells | Up-regulated |
| GO:1903706 | regulation of hemopoiesis | 43/862 | 0.000867453 | Fibroblasts | Up-regulated |
| GO:0051251 | positive regulation of lymphocyte activation | 9/71 | 0.0009169 | T cells | Up-regulated |
| GO:0043620 | regulation of DNA-templated transcription in response to stress | 13/862 | 0.001084055 | Fibroblasts | Up-regulated |
| GO:0048762 | mesenchymal cell differentiation | 28/862 | 0.001084055 | Fibroblasts | Up-regulated |
| GO:0051438 | regulation of ubiquitin-protein transferase activity | 12/862 | 0.001191864 | Fibroblasts | Up-regulated |
| GO:0043484 | regulation of RNA splicing | 11/237 | 0.001214494 | Macrophages | Up-regulated |
| GO:0009408 | response to heat | 5/43 | 0.001369455 | NK cells | Up-regulated |
| GO:0009408 | response to heat | 6/71 | 0.00137989 | T cells | Up-regulated |
| GO:0006959 | humoral immune response | 8/100 | 0.001392503 | Endocardium | Up-regulated |
| GO:2000573 | positive regulation of DNA biosynthetic process | 6/100 | 0.001461726 | B cells | Up-regulated |
| GO:0006986 | response to unfolded protein | 5/43 | 0.00161714 | NK cells | Up-regulated |
| GO:0001525 | angiogenesis | 50/862 | 0.001746431 | Fibroblasts | Up-regulated |
| GO:0006986 | response to unfolded protein | 6/71 | 0.001748508 | T cells | Up-regulated |
| GO:0098760 | response to interleukin-7 | 3/43 | 0.001849785 | NK cells | Up-regulated |
| GO:0098761 | cellular response to interleukin-7 | 3/43 | 0.001849785 | NK cells | Up-regulated |
| GO:0050821 | protein stabilization | 12/237 | 0.001929687 | Macrophages | Up-regulated |
| GO:0072594 | establishment of protein localization to organelle | 7/39 | 0.001940029 | DC-like cells | Up-regulated |
| GO:0010498 | proteasomal protein catabolic process | 34/535 | 0.001957038 | Endothelial cells | Up-regulated |
| GO:0006986 | response to unfolded protein | 14/535 | 0.001957038 | Endothelial cells | Up-regulated |
| GO:0051973 | positive regulation of telomerase activity | 8/535 | 0.002078704 | Endothelial cells | Up-regulated |
| GO:0002456 | T cell mediated immunity | 6/71 | 0.002103696 | T cells | Up-regulated |
| GO:0070203 | regulation of establishment of protein localization to telomere | 4/237 | 0.002299172 | Macrophages | Up-regulated |
| GO:1901214 | regulation of neuron death | 11/100 | 0.00234305 | B cells | Up-regulated |
| GO:0050870 | positive regulation of T cell activation | 7/71 | 0.002513717 | T cells | Up-regulated |
| GO:0016072 | rRNA metabolic process | 9/100 | 0.002564252 | Endocardium | Up-regulated |
| GO:0042254 | ribosome biogenesis | 10/100 | 0.002564252 | Endocardium | Up-regulated |
| GO:1903362 | regulation of cellular protein catabolic process | 14/237 | 0.002628482 | Macrophages | Up-regulated |
| GO:1903050 | regulation of proteolysis involved in cellular protein catabolic process | 13/237 | 0.002628482 | Macrophages | Up-regulated |
| GO:0061136 | regulation of proteasomal protein catabolic process | 12/237 | 0.002628482 | Macrophages | Up-regulated |
| GO:1901998 | toxin transport | 6/237 | 0.002628482 | Macrophages | Up-regulated |
| GO:0001935 | endothelial cell proliferation | 16/535 | 0.002837603 | Endothelial cells | Up-regulated |
| GO:0010632 | regulation of epithelial cell migration | 28/862 | 0.003021561 | Fibroblasts | Up-regulated |
| GO:0002709 | regulation of T cell mediated immunity | 5/71 | 0.003034908 | T cells | Up-regulated |
| GO:0010743 | regulation of macrophage derived foam cell differentiation | 6/535 | 0.00308254 | Endothelial cells | Up-regulated |
| GO:0006606 | protein import into nucleus | 21/862 | 0.003221387 | Fibroblasts | Up-regulated |
| GO:0006913 | nucleocytoplasmic transport | 33/862 | 0.003221387 | Fibroblasts | Up-regulated |
| GO:0051169 | nuclear transport | 33/862 | 0.003221387 | Fibroblasts | Up-regulated |
| GO:1903039 | positive regulation of leukocyte cell-cell adhesion | 7/71 | 0.003293012 | T cells | Up-regulated |
| GO:0033002 | muscle cell proliferation | 28/862 | 0.003366313 | Fibroblasts | Up-regulated |
| GO:0061136 | regulation of proteasomal protein catabolic process | 8/100 | 0.00340966 | B cells | Up-regulated |
| GO:0042326 | negative regulation of phosphorylation | 12/100 | 0.00340966 | B cells | Up-regulated |
| GO:1903362 | regulation of cellular protein catabolic process | 9/100 | 0.00340966 | B cells | Up-regulated |
| GO:0006749 | glutathione metabolic process | 6/119 | 0.003507026 | Pro.cells | Up-regulated |
| GO:0017038 | protein import | 26/862 | 0.003951824 | Fibroblasts | Up-regulated |
| GO:0001937 | negative regulation of endothelial cell proliferation | 8/535 | 0.004112606 | Endothelial cells | Up-regulated |
| GO:0070997 | neuron death | 11/100 | 0.004154535 | B cells | Up-regulated |
| GO:0035116 | embryonic hindlimb morphogenesis | 8/862 | 0.004188948 | Fibroblasts | Up-regulated |
| GO:0006278 | RNA-dependent DNA biosynthetic process | 7/237 | 0.00425141 | Macrophages | Up-regulated |
| GO:0007004 | telomere maintenance via telomerase | 7/237 | 0.00425141 | Macrophages | Up-regulated |
| GO:0072655 | establishment of protein localization to mitochondrion | 5/71 | 0.00507597 | T cells | Up-regulated |
| GO:0043618 | regulation of transcription from RNA polymerase II promoter in response to stress | 9/535 | 0.005201522 | Endothelial cells | Up-regulated |
| GO:1903523 | negative regulation of blood circulation | 5/119 | 0.005267701 | Pro.cells | Up-regulated |
| GO:0032963 | collagen metabolic process | 7/119 | 0.005267701 | Pro.cells | Up-regulated |
| GO:0046677 | response to antibiotic | 27/862 | 0.005499587 | Fibroblasts | Up-regulated |
| GO:0009948 | anterior/posterior axis specification | 8/535 | 0.005779444 | Endothelial cells | Up-regulated |
| GO:0007339 | binding of sperm to zona pellucida | 5/237 | 0.005824337 | Macrophages | Up-regulated |
| GO:0010812 | negative regulation of cell-substrate adhesion | 12/862 | 0.005892661 | Fibroblasts | Up-regulated |
| GO:0031050 | dsRNA processing | 10/862 | 0.005962419 | Fibroblasts | Up-regulated |
| GO:0001933 | negative regulation of protein phosphorylation | 9/71 | 0.006139027 | T cells | Up-regulated |
| GO:0007159 | leukocyte cell-cell adhesion | 8/71 | 0.006139027 | T cells | Up-regulated |
| GO:1902107 | positive regulation of leukocyte differentiation | 6/71 | 0.006139027 | T cells | Up-regulated |
| GO:0031341 | regulation of cell killing | 5/71 | 0.006139027 | T cells | Up-regulated |
| GO:0002706 | regulation of lymphocyte mediated immunity | 6/71 | 0.006139027 | T cells | Up-regulated |
| GO:0043524 | negative regulation of neuron apoptotic process | 6/71 | 0.006139027 | T cells | Up-regulated |
| GO:0071900 | regulation of protein serine/threonine kinase activity | 9/71 | 0.006139027 | T cells | Up-regulated |
| GO:0010657 | muscle cell apoptotic process | 16/862 | 0.0062114 | Fibroblasts | Up-regulated |
| GO:0051131 | chaperone-mediated protein complex assembly | 4/237 | 0.006234299 | Macrophages | Up-regulated |
| GO:0070200 | establishment of protein localization to telomere | 4/237 | 0.006234299 | Macrophages | Up-regulated |
| GO:1902253 | regulation of intrinsic apoptotic signaling pathway by p53 class mediator | 4/100 | 0.006585512 | Endocardium | Up-regulated |
| GO:0061614 | pri-miRNA transcription by RNA polymerase II | 9/535 | 0.006805232 | Endothelial cells | Up-regulated |
| GO:0002455 | humoral immune response mediated by circulating immunoglobulin | 5/100 | 0.00683149 | Endocardium | Up-regulated |
| GO:0031647 | regulation of protein stability | 23/535 | 0.007219925 | Endothelial cells | Up-regulated |
| GO:0097201 | negative regulation of transcription from RNA polymerase II promoter in response to stress | 3/100 | 0.007220989 | B cells | Up-regulated |
| GO:0031331 | positive regulation of cellular catabolic process | 10/100 | 0.007827806 | B cells | Up-regulated |
| GO:0009988 | cell-cell recognition | 6/237 | 0.007993922 | Macrophages | Up-regulated |
| GO:0000723 | telomere maintenance | 14/535 | 0.008206947 | Endothelial cells | Up-regulated |
| GO:0071480 | cellular response to gamma radiation | 8/862 | 0.008234091 | Fibroblasts | Up-regulated |
| GO:0032609 | interferon-gamma production | 5/71 | 0.008252427 | T cells | Up-regulated |
| GO:1905897 | regulation of response to endoplasmic reticulum stress | 7/237 | 0.008294493 | Macrophages | Up-regulated |
| GO:0098760 | response to interleukin-7 | 4/237 | 0.008294493 | Macrophages | Up-regulated |
| GO:0098761 | cellular response to interleukin-7 | 4/237 | 0.008294493 | Macrophages | Up-regulated |
| GO:0001503 | ossification | 36/862 | 0.008764906 | Fibroblasts | Up-regulated |
| GO:0072655 | establishment of protein localization to mitochondrion | 4/43 | 0.00883371 | NK cells | Up-regulated |
| GO:0046825 | regulation of protein export from nucleus | 3/39 | 0.008835075 | DC-like cells | Up-regulated |
| GO:0032200 | telomere organization | 9/237 | 0.008995406 | Macrophages | Up-regulated |
| GO:0032200 | telomere organization | 14/535 | 0.009344515 | Endothelial cells | Up-regulated |
| GO:0003158 | endothelium development | 13/535 | 0.009676144 | Endothelial cells | Up-regulated |
| GO:0002683 | negative regulation of immune system process | 31/535 | 0.009916153 | Endothelial cells | Up-regulated |
| GO:0019886 | antigen processing and presentation of exogenous peptide antigen via MHC class II | 3/100 | 0.010057108 | Endocardium | Up-regulated |
| GO:0045446 | endothelial cell differentiation | 12/535 | 0.010294175 | Endothelial cells | Up-regulated |
| GO:0072594 | establishment of protein localization to organelle | 10/100 | 0.010384022 | B cells | Up-regulated |
| GO:0002449 | lymphocyte mediated immunity | 7/71 | 0.010544531 | T cells | Up-regulated |
| GO:0051702 | interaction with symbiont | 4/71 | 0.010544531 | T cells | Up-regulated |
| GO:0032516 | positive regulation of phosphoprotein phosphatase activity | 3/100 | 0.010567242 | B cells | Up-regulated |
| GO:0051131 | chaperone-mediated protein complex assembly | 3/100 | 0.010567242 | B cells | Up-regulated |
| GO:0034620 | cellular response to unfolded protein | 5/100 | 0.010567242 | B cells | Up-regulated |
| GO:0010498 | proteasomal protein catabolic process | 18/237 | 0.010849364 | Macrophages | Up-regulated |
| GO:0051347 | positive regulation of transferase activity | 49/862 | 0.010925004 | Fibroblasts | Up-regulated |
| GO:0010660 | regulation of muscle cell apoptotic process | 15/862 | 0.010978354 | Fibroblasts | Up-regulated |
| GO:0045601 | regulation of endothelial cell differentiation | 7/535 | 0.010980508 | Endothelial cells | Up-regulated |
| GO:0042698 | ovulation cycle | 10/862 | 0.011118639 | Fibroblasts | Up-regulated |
| GO:0010498 | proteasomal protein catabolic process | 11/100 | 0.011357471 | B cells | Up-regulated |
| GO:0006986 | response to unfolded protein | 16/862 | 0.011608976 | Fibroblasts | Up-regulated |
| GO:0060148 | positive regulation of posttranscriptional gene silencing | 7/862 | 0.011608976 | Fibroblasts | Up-regulated |
| GO:0001667 | ameboidal-type cell migration | 39/862 | 0.011608976 | Fibroblasts | Up-regulated |
| GO:0001649 | osteoblast differentiation | 22/862 | 0.011608976 | Fibroblasts | Up-regulated |
| GO:0006403 | RNA localization | 22/862 | 0.011608976 | Fibroblasts | Up-regulated |
| GO:2001235 | positive regulation of apoptotic signaling pathway | 7/100 | 0.011795267 | B cells | Up-regulated |
| GO:0017038 | protein import | 6/71 | 0.012051842 | T cells | Up-regulated |
| GO:0030278 | regulation of ossification | 25/862 | 0.012686705 | Fibroblasts | Up-regulated |
| GO:0017038 | protein import | 18/535 | 0.012963352 | Endothelial cells | Up-regulated |
| GO:0001906 | cell killing | 5/71 | 0.012982591 | T cells | Up-regulated |
| GO:0051260 | protein homooligomerization | 15/237 | 0.013426867 | Macrophages | Up-regulated |
| GO:0022613 | ribonucleoprotein complex biogenesis | 17/237 | 0.013426867 | Macrophages | Up-regulated |
| GO:0098727 | maintenance of cell number | 15/535 | 0.013536529 | Endothelial cells | Up-regulated |
| GO:0042326 | negative regulation of phosphorylation | 30/535 | 0.014121158 | Endothelial cells | Up-regulated |
| GO:0021954 | central nervous system neuron development | 4/71 | 0.014165283 | T cells | Up-regulated |
| GO:0009266 | response to temperature stimulus | 5/40 | 0.014243994 | Schwann cells | Up-regulated |
| GO:0001933 | negative regulation of protein phosphorylation | 10/100 | 0.01438749 | B cells | Up-regulated |
| GO:0048569 | post-embryonic animal organ development | 5/535 | 0.014442442 | Endothelial cells | Up-regulated |
| GO:0031396 | regulation of protein ubiquitination | 11/237 | 0.014747784 | Macrophages | Up-regulated |
| GO:1901861 | regulation of muscle tissue development | 20/862 | 0.014865065 | Fibroblasts | Up-regulated |
| GO:0048524 | positive regulation of viral process | 4/71 | 0.015174951 | T cells | Up-regulated |
| GO:0002521 | leukocyte differentiation | 9/71 | 0.015174951 | T cells | Up-regulated |
| GO:1903708 | positive regulation of hemopoiesis | 7/100 | 0.015228576 | B cells | Up-regulated |
| GO:0045445 | myoblast differentiation | 13/862 | 0.015307509 | Fibroblasts | Up-regulated |
| GO:0061684 | chaperone-mediated autophagy | 2/39 | 0.015323477 | DC-like cells | Up-regulated |
| GO:0071276 | cellular response to cadmium ion | 3/100 | 0.015363312 | B cells | Up-regulated |
| GO:0050821 | protein stabilization | 21/862 | 0.015580687 | Fibroblasts | Up-regulated |
| GO:0032799 | low-density lipoprotein receptor particle metabolic process | 4/535 | 0.015853052 | Endothelial cells | Up-regulated |
| GO:0070203 | regulation of establishment of protein localization to telomere | 4/535 | 0.015853052 | Endothelial cells | Up-regulated |
| GO:0060416 | response to growth hormone | 6/862 | 0.01596135 | Fibroblasts | Up-regulated |
| GO:0051085 | chaperone cofactor-dependent protein refolding | 3/40 | 0.016021384 | Schwann cells | Up-regulated |
| GO:0046329 | negative regulation of JNK cascade | 3/40 | 0.016021384 | Schwann cells | Up-regulated |
| GO:1903573 | negative regulation of response to endoplasmic reticulum stress | 5/237 | 0.016066711 | Macrophages | Up-regulated |
| GO:0006403 | RNA localization | 10/237 | 0.016066711 | Macrophages | Up-regulated |
| GO:0002460 | adaptive immune response based on somatic recombination of immune receptors built from immunoglobulin superfamily domains | 31/862 | 0.016362204 | Fibroblasts | Up-regulated |
| GO:0045785 | positive regulation of cell adhesion | 39/862 | 0.016362204 | Fibroblasts | Up-regulated |
| GO:0016032 | viral process | 29/862 | 0.016406043 | Fibroblasts | Up-regulated |
| GO:0050870 | positive regulation of T cell activation | 22/862 | 0.016823164 | Fibroblasts | Up-regulated |
| GO:0010332 | response to gamma radiation | 10/862 | 0.016823164 | Fibroblasts | Up-regulated |
| GO:0009612 | response to mechanical stimulus | 6/87 | 0.016968079 | Pericytes | Up-regulated |
| GO:0051345 | positive regulation of hydrolase activity | 11/100 | 0.017043701 | B cells | Up-regulated |
| GO:0009896 | positive regulation of catabolic process | 10/100 | 0.017043701 | B cells | Up-regulated |
| GO:0070198 | protein localization to chromosome, telomeric region | 6/535 | 0.017118821 | Endothelial cells | Up-regulated |
| GO:0060009 | Sertoli cell development | 5/535 | 0.017485311 | Endothelial cells | Up-regulated |
| GO:0048660 | regulation of smooth muscle cell proliferation | 20/862 | 0.018203251 | Fibroblasts | Up-regulated |
| GO:0032870 | cellular response to hormone stimulus | 42/862 | 0.018388583 | Fibroblasts | Up-regulated |
| GO:1990823 | response to leukemia inhibitory factor | 17/862 | 0.018498729 | Fibroblasts | Up-regulated |
| GO:1990830 | cellular response to leukemia inhibitory factor | 17/862 | 0.018498729 | Fibroblasts | Up-regulated |
| GO:0015909 | long-chain fatty acid transport | 5/237 | 0.018532177 | Macrophages | Up-regulated |
| GO:0009123 | nucleoside monophosphate metabolic process | 12/237 | 0.018791701 | Macrophages | Up-regulated |
| GO:0044319 | wound healing, spreading of cells | 7/862 | 0.018837497 | Fibroblasts | Up-regulated |
| GO:0090505 | epiboly involved in wound healing | 7/862 | 0.018837497 | Fibroblasts | Up-regulated |
| GO:0070303 | negative regulation of stress-activated protein kinase signaling cascade | 3/40 | 0.018957005 | Schwann cells | Up-regulated |
| GO:0021955 | central nervous system neuron axonogenesis | 3/71 | 0.018993317 | T cells | Up-regulated |
| GO:0040013 | negative regulation of locomotion | 32/862 | 0.019018053 | Fibroblasts | Up-regulated |
| GO:0042445 | hormone metabolic process | 20/862 | 0.019311853 | Fibroblasts | Up-regulated |
| GO:0006958 | complement activation, classical pathway | 4/100 | 0.019325181 | Endocardium | Up-regulated |
| GO:0010717 | regulation of epithelial to mesenchymal transition | 13/862 | 0.019543008 | Fibroblasts | Up-regulated |
| GO:0051348 | negative regulation of transferase activity | 6/71 | 0.019611322 | T cells | Up-regulated |
| GO:0002711 | positive regulation of T cell mediated immunity | 4/100 | 0.019631884 | B cells | Up-regulated |
| GO:0031647 | regulation of protein stability | 8/100 | 0.019631884 | B cells | Up-regulated |
| GO:0045428 | regulation of nitric oxide biosynthetic process | 9/535 | 0.019716254 | Endothelial cells | Up-regulated |
| GO:2001252 | positive regulation of chromosome organization | 16/535 | 0.019716254 | Endothelial cells | Up-regulated |
| GO:0044419 | interspecies interaction between organisms | 25/535 | 0.019716254 | Endothelial cells | Up-regulated |
| GO:0048871 | multicellular organismal homeostasis | 28/535 | 0.019716254 | Endothelial cells | Up-regulated |
| GO:0007062 | sister chromatid cohesion | 8/535 | 0.019716254 | Endothelial cells | Up-regulated |
| GO:0007611 | learning or memory | 6/71 | 0.019783115 | T cells | Up-regulated |
| GO:0140014 | mitotic nuclear division | 20/535 | 0.019897189 | Endothelial cells | Up-regulated |
| GO:0097201 | negative regulation of transcription from RNA polymerase II promoter in response to stress | 2/39 | 0.020451648 | DC-like cells | Up-regulated |
| GO:0009896 | positive regulation of catabolic process | 28/535 | 0.020929929 | Endothelial cells | Up-regulated |
| GO:0031341 | regulation of cell killing | 5/100 | 0.021022393 | B cells | Up-regulated |
| GO:0009266 | response to temperature stimulus | 19/862 | 0.021225379 | Fibroblasts | Up-regulated |
| GO:1901214 | regulation of neuron death | 9/87 | 0.021345566 | Pericytes | Up-regulated |
| GO:0072655 | establishment of protein localization to mitochondrion | 5/87 | 0.02157866 | Pericytes | Up-regulated |
| GO:1903747 | regulation of establishment of protein localization to mitochondrion | 3/100 | 0.021593407 | B cells | Up-regulated |
| GO:0001933 | negative regulation of protein phosphorylation | 27/535 | 0.021750953 | Endothelial cells | Up-regulated |
| GO:0009404 | toxin metabolic process | 6/862 | 0.021834352 | Fibroblasts | Up-regulated |
| GO:1903409 | reactive oxygen species biosynthetic process | 12/535 | 0.022715035 | Endothelial cells | Up-regulated |
| GO:0043902 | positive regulation of multi-organism process | 5/71 | 0.022774184 | T cells | Up-regulated |
| GO:0010977 | negative regulation of neuron projection development | 5/71 | 0.023122431 | T cells | Up-regulated |
| GO:0060412 | ventricular septum morphogenesis | 9/862 | 0.024075665 | Fibroblasts | Up-regulated |
| GO:0042110 | T cell activation | 8/71 | 0.024176045 | T cells | Up-regulated |
| GO:1901214 | regulation of neuron death | 6/43 | 0.024662273 | NK cells | Up-regulated |
| GO:1901215 | negative regulation of neuron death | 5/43 | 0.024662273 | NK cells | Up-regulated |
| GO:0007179 | transforming growth factor beta receptor signaling pathway | 6/100 | 0.024906817 | B cells | Up-regulated |
| GO:0002696 | positive regulation of leukocyte activation | 32/862 | 0.025215757 | Fibroblasts | Up-regulated |
| GO:0071236 | cellular response to antibiotic | 17/862 | 0.025826767 | Fibroblasts | Up-regulated |
| GO:0001912 | positive regulation of leukocyte mediated cytotoxicity | 4/100 | 0.026162118 | B cells | Up-regulated |
| GO:0007004 | telomere maintenance via telomerase | 4/100 | 0.026162118 | B cells | Up-regulated |
| GO:1904950 | negative regulation of establishment of protein localization | 10/237 | 0.026274327 | Macrophages | Up-regulated |
| GO:0044419 | interspecies interaction between organisms | 34/862 | 0.026384683 | Fibroblasts | Up-regulated |
| GO:2000737 | negative regulation of stem cell differentiation | 6/862 | 0.026384683 | Fibroblasts | Up-regulated |
| GO:0010463 | mesenchymal cell proliferation | 10/862 | 0.026596639 | Fibroblasts | Up-regulated |
| GO:0072593 | reactive oxygen species metabolic process | 6/71 | 0.026782282 | T cells | Up-regulated |
| GO:0048024 | regulation of mRNA splicing, via spliceosome | 5/87 | 0.026916413 | Pericytes | Up-regulated |
| GO:0048675 | axon extension | 4/43 | 0.027110534 | NK cells | Up-regulated |
| GO:0042632 | cholesterol homeostasis | 9/535 | 0.027221593 | Endothelial cells | Up-regulated |
| GO:1903332 | regulation of protein folding | 2/43 | 0.027958053 | NK cells | Up-regulated |
| GO:1903362 | regulation of cellular protein catabolic process | 5/43 | 0.027958053 | NK cells | Up-regulated |
| GO:1903364 | positive regulation of cellular protein catabolic process | 4/43 | 0.027958053 | NK cells | Up-regulated |
| GO:0016032 | viral process | 6/71 | 0.029186481 | T cells | Up-regulated |
| GO:0050890 | cognition | 6/71 | 0.029186481 | T cells | Up-regulated |
| GO:0046686 | response to cadmium ion | 3/100 | 0.029230499 | B cells | Up-regulated |
| GO:0070997 | neuron death | 9/87 | 0.02993281 | Pericytes | Up-regulated |
| GO:0070997 | neuron death | 6/43 | 0.030473732 | NK cells | Up-regulated |
| GO:0097201 | negative regulation of transcription from RNA polymerase II promoter in response to stress | 2/71 | 0.030515532 | T cells | Up-regulated |
| GO:0050679 | positive regulation of epithelial cell proliferation | 21/862 | 0.031388918 | Fibroblasts | Up-regulated |
| GO:0031331 | positive regulation of cellular catabolic process | 24/535 | 0.03190526 | Endothelial cells | Up-regulated |
| GO:0050684 | regulation of mRNA processing | 4/40 | 0.032064456 | Schwann cells | Up-regulated |
| GO:0090307 | mitotic spindle assembly | 3/40 | 0.032064456 | Schwann cells | Up-regulated |
| GO:0033523 | histone H2B ubiquitination | 2/40 | 0.032064456 | Schwann cells | Up-regulated |
| GO:0031331 | positive regulation of cellular catabolic process | 14/237 | 0.032085246 | Macrophages | Up-regulated |
| GO:0043371 | negative regulation of CD4-positive, alpha-beta T cell differentiation | 6/862 | 0.032121735 | Fibroblasts | Up-regulated |
| GO:0051702 | interaction with symbiont | 4/100 | 0.032261627 | B cells | Up-regulated |
| GO:0032092 | positive regulation of protein binding | 4/71 | 0.032363658 | T cells | Up-regulated |
| GO:0051402 | neuron apoptotic process | 6/71 | 0.032363658 | T cells | Up-regulated |
| GO:0062012 | regulation of small molecule metabolic process | 31/862 | 0.0324078 | Fibroblasts | Up-regulated |
| GO:0051101 | regulation of DNA binding | 12/535 | 0.032550007 | Endothelial cells | Up-regulated |
| GO:0032355 | response to estradiol | 8/862 | 0.032592596 | Fibroblasts | Up-regulated |
| GO:0032507 | maintenance of protein location in cell | 3/39 | 0.032675157 | DC-like cells | Up-regulated |
| GO:1902176 | negative regulation of oxidative stress-induced intrinsic apoptotic signaling pathway | 2/39 | 0.032675157 | DC-like cells | Up-regulated |
| GO:0001819 | positive regulation of cytokine production | 38/862 | 0.033048079 | Fibroblasts | Up-regulated |
| GO:0019395 | fatty acid oxidation | 10/535 | 0.033075072 | Endothelial cells | Up-regulated |
| GO:0006809 | nitric oxide biosynthetic process | 9/535 | 0.034027968 | Endothelial cells | Up-regulated |
| GO:0050657 | nucleic acid transport | 18/862 | 0.034435901 | Fibroblasts | Up-regulated |
| GO:0050658 | RNA transport | 18/862 | 0.034435901 | Fibroblasts | Up-regulated |
| GO:0051101 | regulation of DNA binding | 5/100 | 0.034763123 | B cells | Up-regulated |
| GO:0071407 | cellular response to organic cyclic compound | 35/862 | 0.035049619 | Fibroblasts | Up-regulated |
| GO:1904019 | epithelial cell apoptotic process | 11/535 | 0.035449927 | Endothelial cells | Up-regulated |
| GO:0000054 | ribosomal subunit export from nucleus | 3/237 | 0.035581975 | Macrophages | Up-regulated |
| GO:0033750 | ribosome localization | 3/237 | 0.035581975 | Macrophages | Up-regulated |
| GO:0046661 | male sex differentiation | 12/535 | 0.035914999 | Endothelial cells | Up-regulated |
| GO:0019058 | viral life cycle | 6/100 | 0.036534452 | B cells | Up-regulated |
| GO:0097193 | intrinsic apoptotic signaling pathway | 6/71 | 0.037015463 | T cells | Up-regulated |
| GO:0008361 | regulation of cell size | 5/71 | 0.037016152 | T cells | Up-regulated |
| GO:0090151 | establishment of protein localization to mitochondrial membrane | 2/71 | 0.037022234 | T cells | Up-regulated |
| GO:0002274 | myeloid leukocyte activation | 5/71 | 0.037212297 | T cells | Up-regulated |
| GO:0031345 | negative regulation of cell projection organization | 5/71 | 0.037212297 | T cells | Up-regulated |
| GO:0030099 | myeloid cell differentiation | 35/862 | 0.037521783 | Fibroblasts | Up-regulated |
| GO:0006413 | translational initiation | 7/237 | 0.03813348 | Macrophages | Up-regulated |
| GO:0016032 | viral process | 5/23 | 0.038291503 | Granulocytes | Up-regulated |
| GO:0044403 | symbiont process | 5/23 | 0.038291503 | Granulocytes | Up-regulated |
| GO:0044419 | interspecies interaction between organisms | 5/23 | 0.038291503 | Granulocytes | Up-regulated |
| GO:0019058 | viral life cycle | 4/23 | 0.038291503 | Granulocytes | Up-regulated |
| GO:0034620 | cellular response to unfolded protein | 9/535 | 0.038652311 | Endothelial cells | Up-regulated |
| GO:1990823 | response to leukemia inhibitory factor | 12/535 | 0.038652311 | Endothelial cells | Up-regulated |
| GO:1990830 | cellular response to leukemia inhibitory factor | 12/535 | 0.038652311 | Endothelial cells | Up-regulated |
| GO:0090083 | regulation of inclusion body assembly | 4/535 | 0.038652311 | Endothelial cells | Up-regulated |
| GO:0030449 | regulation of complement activation | 5/862 | 0.038878874 | Fibroblasts | Up-regulated |
| GO:1901361 | organic cyclic compound catabolic process | 40/862 | 0.039028973 | Fibroblasts | Up-regulated |
| GO:0035690 | cellular response to drug | 29/862 | 0.039353378 | Fibroblasts | Up-regulated |
| GO:0046631 | alpha-beta T cell activation | 18/862 | 0.039353378 | Fibroblasts | Up-regulated |
| GO:0002486 | antigen processing and presentation of endogenous peptide antigen via MHC class I via ER pathway, TAP-independent | 2/71 | 0.039614193 | T cells | Up-regulated |
| GO:0051131 | chaperone-mediated protein complex assembly | 2/71 | 0.039614193 | T cells | Up-regulated |
| GO:1990000 | amyloid fibril formation | 2/71 | 0.039614193 | T cells | Up-regulated |
| GO:0048285 | organelle fission | 27/535 | 0.040037162 | Endothelial cells | Up-regulated |
| GO:0032200 | telomere organization | 5/100 | 0.040156093 | B cells | Up-regulated |
| GO:0007162 | negative regulation of cell adhesion | 20/535 | 0.040189494 | Endothelial cells | Up-regulated |
| GO:0001935 | endothelial cell proliferation | 17/862 | 0.040575122 | Fibroblasts | Up-regulated |
| GO:0008585 | female gonad development | 13/862 | 0.040575122 | Fibroblasts | Up-regulated |
| GO:0051881 | regulation of mitochondrial membrane potential | 4/100 | 0.040992928 | B cells | Up-regulated |
| GO:0045793 | positive regulation of cell size | 2/43 | 0.04128127 | NK cells | Up-regulated |
| GO:0051131 | chaperone-mediated protein complex assembly | 2/43 | 0.04128127 | NK cells | Up-regulated |
| GO:0070988 | demethylation | 5/237 | 0.041305393 | Macrophages | Up-regulated |
| GO:0050880 | regulation of blood vessel size | 7/119 | 0.041506724 | Pro.cells | Up-regulated |
| GO:0035150 | regulation of tube size | 7/119 | 0.041506724 | Pro.cells | Up-regulated |
| GO:0033555 | multicellular organismal response to stress | 3/23 | 0.042271459 | Granulocytes | Up-regulated |
| GO:0051347 | positive regulation of transferase activity | 8/71 | 0.042367991 | T cells | Up-regulated |
| GO:0002476 | antigen processing and presentation of endogenous peptide antigen via MHC class Ib | 2/71 | 0.04249084 | T cells | Up-regulated |
| GO:0045732 | positive regulation of protein catabolic process | 5/71 | 0.04249084 | T cells | Up-regulated |
| GO:0006839 | mitochondrial transport | 9/237 | 0.042913208 | Macrophages | Up-regulated |
| GO:0021954 | central nervous system neuron development | 4/100 | 0.042978087 | B cells | Up-regulated |
| GO:0009060 | aerobic respiration | 8/535 | 0.0429982 | Endothelial cells | Up-regulated |
| GO:0002521 | leukocyte differentiation | 10/100 | 0.043221751 | B cells | Up-regulated |
| GO:0006110 | regulation of glycolytic process | 3/100 | 0.043221751 | B cells | Up-regulated |
| GO:0030811 | regulation of nucleotide catabolic process | 3/100 | 0.043221751 | B cells | Up-regulated |
| GO:1901998 | toxin transport | 3/100 | 0.043221751 | B cells | Up-regulated |
| GO:0071559 | response to transforming growth factor beta | 6/100 | 0.043221751 | B cells | Up-regulated |
| GO:0048638 | regulation of developmental growth | 34/862 | 0.043534292 | Fibroblasts | Up-regulated |
| GO:0010458 | exit from mitosis | 6/862 | 0.043534292 | Fibroblasts | Up-regulated |
| GO:0061311 | cell surface receptor signaling pathway involved in heart development | 6/862 | 0.043534292 | Fibroblasts | Up-regulated |
| GO:0001906 | cell killing | 5/100 | 0.043540198 | B cells | Up-regulated |
| GO:1905710 | positive regulation of membrane permeability | 3/100 | 0.044223421 | B cells | Up-regulated |
| GO:0031589 | cell-substrate adhesion | 31/862 | 0.04497235 | Fibroblasts | Up-regulated |
| GO:0042537 | benzene-containing compound metabolic process | 5/862 | 0.0453509 | Fibroblasts | Up-regulated |
| GO:2000257 | regulation of protein activation cascade | 5/862 | 0.0453509 | Fibroblasts | Up-regulated |
| GO:0032922 | circadian regulation of gene expression | 10/862 | 0.0453509 | Fibroblasts | Up-regulated |
| GO:0002698 | negative regulation of immune effector process | 16/862 | 0.0453509 | Fibroblasts | Up-regulated |
| GO:0006622 | protein targeting to lysosome | 2/71 | 0.045759557 | T cells | Up-regulated |
| GO:0098760 | response to interleukin-7 | 2/71 | 0.045759557 | T cells | Up-regulated |
| GO:0098761 | cellular response to interleukin-7 | 2/71 | 0.045759557 | T cells | Up-regulated |
| GO:0010656 | negative regulation of muscle cell apoptotic process | 5/237 | 0.046658106 | Macrophages | Up-regulated |
| GO:0050729 | positive regulation of inflammatory response | 4/71 | 0.046911814 | T cells | Up-regulated |
| GO:0045862 | positive regulation of proteolysis | 6/71 | 0.046911814 | T cells | Up-regulated |
| GO:0045123 | cellular extravasation | 3/71 | 0.046911814 | T cells | Up-regulated |
| GO:0051851 | modification by host of symbiont morphology or physiology | 3/71 | 0.046911814 | T cells | Up-regulated |
| GO:0044794 | positive regulation by host of viral process | 2/23 | 0.047495806 | Granulocytes | Up-regulated |
| GO:0045930 | negative regulation of mitotic cell cycle | 4/23 | 0.047495806 | Granulocytes | Up-regulated |
| GO:1904950 | negative regulation of establishment of protein localization | 4/39 | 0.048017013 | DC-like cells | Up-regulated |
| GO:0014015 | positive regulation of gliogenesis | 12/862 | 0.048083375 | Fibroblasts | Up-regulated |
| GO:2001038 | regulation of cellular response to drug | 7/862 | 0.048369566 | Fibroblasts | Up-regulated |
| GO:0071466 | cellular response to xenobiotic stimulus | 13/862 | 0.048909814 | Fibroblasts | Up-regulated |
| GO:0044270 | cellular nitrogen compound catabolic process | 37/862 | 0.049187216 | Fibroblasts | Up-regulated |
| GO:0032870 | cellular response to hormone stimulus | 9/100 | 0.049606139 | B cells | Up-regulated |
| GO:0055078 | sodium ion homeostasis | 4/119 | 0.049670852 | Pro.cells | Up-regulated |
| GO:0060070 | canonical Wnt signaling pathway | 19/535 | 0.04968826 | Endothelial cells | Up-regulated |
| GO:1901673 | regulation of mitotic spindle assembly | 2/39 | 0.049800536 | DC-like cells | Up-regulated |
| GO:0009896 | positive regulation of catabolic process | 15/237 | 0.049966456 | Macrophages | Up-regulated |

**Supplementary Figures**

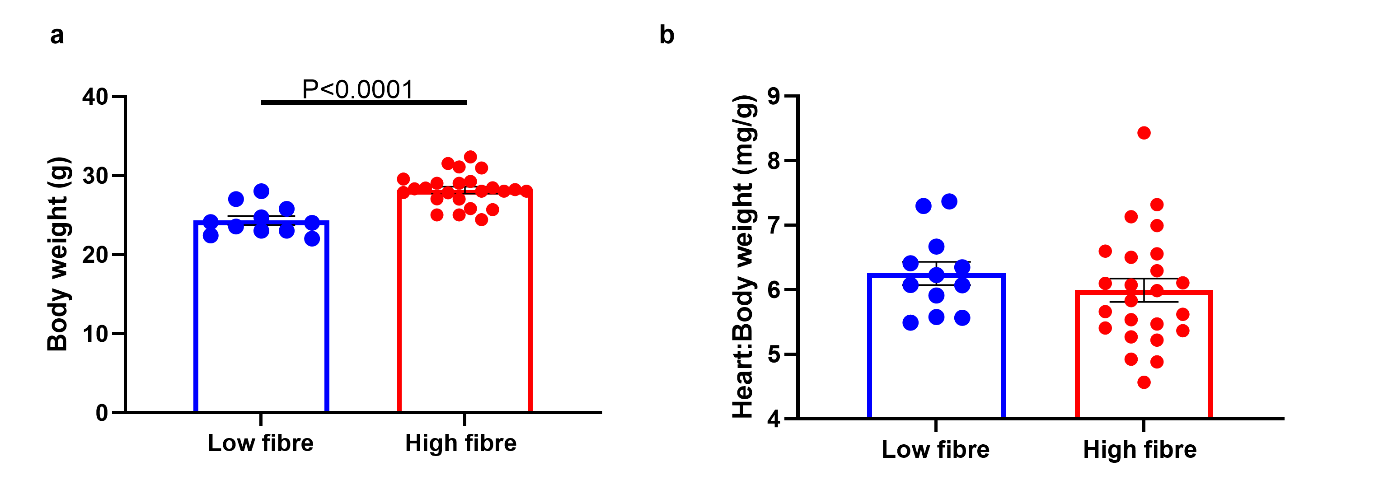

**Supplementary Figure 1.** Post-weaning a) body weight and b) heart to body weight ratio of high- and low-fibre fed dams. All data is shown as ±SEM.n=11-24/group.

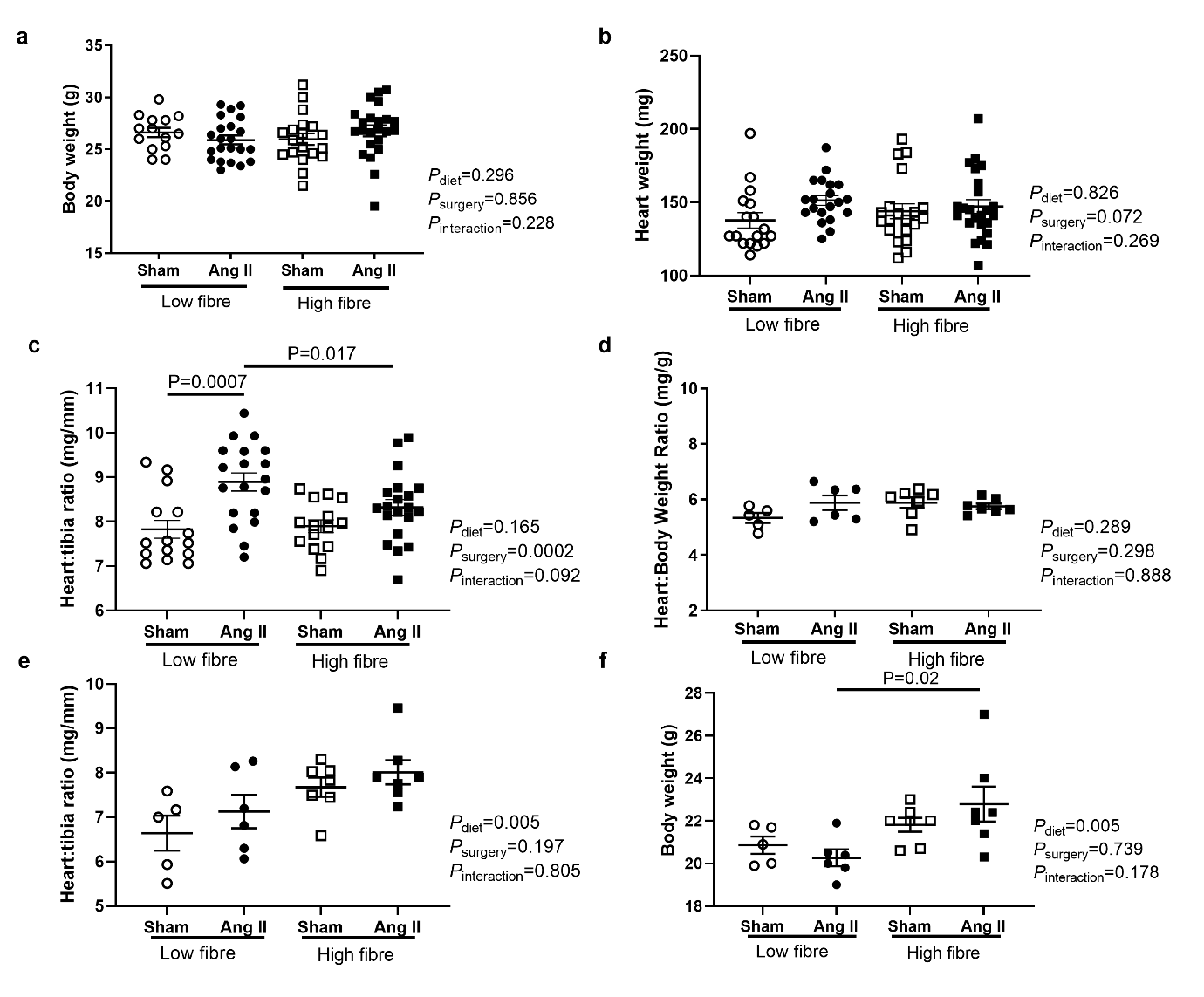

**Supplementary Figure 2.** a) Body weight, b) heart weight, and c) heart to tibia ratio of male high-fibre and low-fibre offspring. d) body weight, heart to body weight ratio, and f) heart to tibia ratio of female offspring from high-fibre and low-fibre female offspring. Sample size: a-c n=15-23. d-f n=5-7. All data is shown as mean±SEM. 2-way ANOVA with FDR adjustment for multiple comparisons.

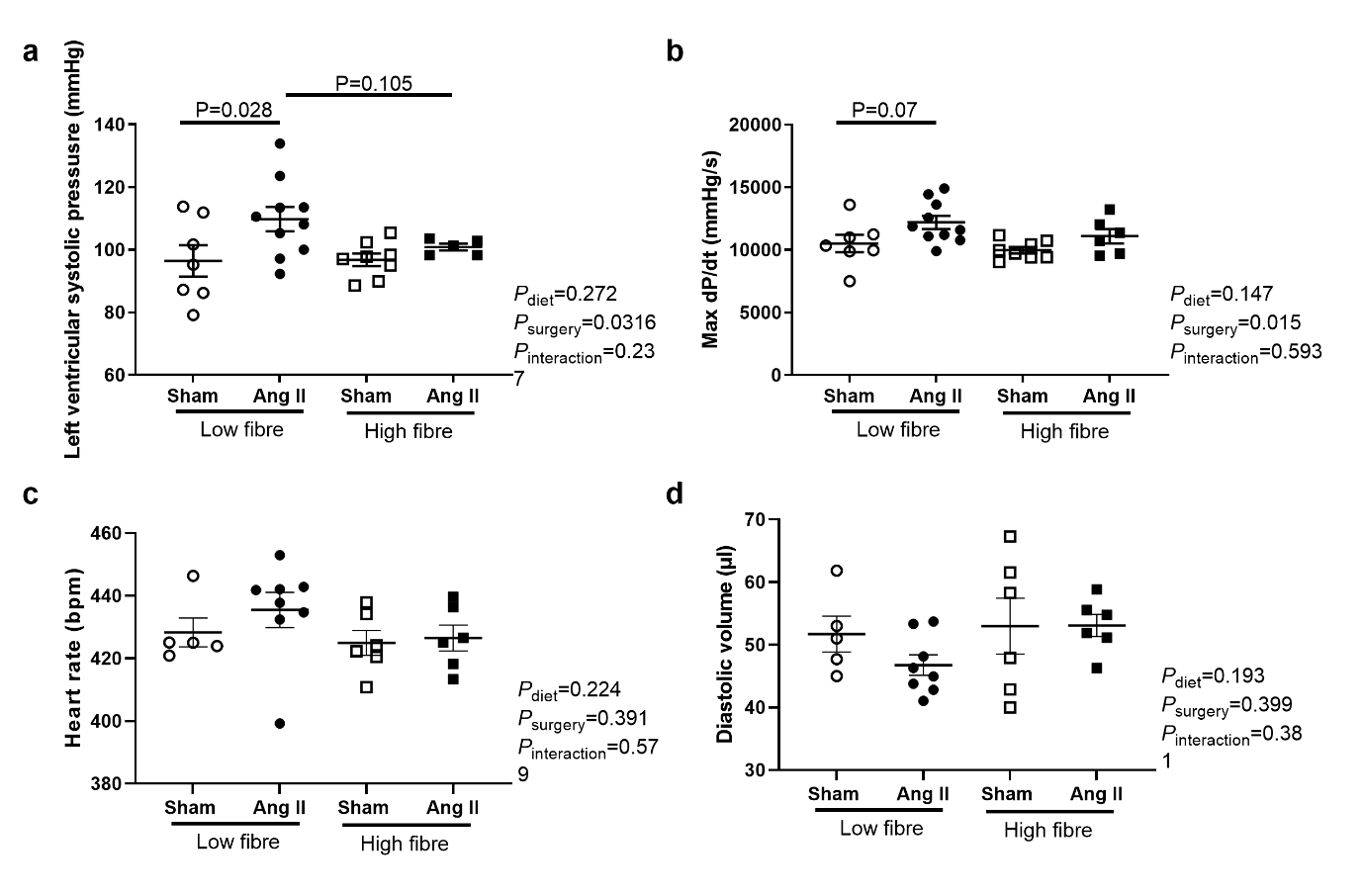
**Supplementary Figure 3.** Cardiac catherization measurements showing a) left ventricular systolic pressure, and b) cardiac contractility (dp/dt) in male high-fibre and low-fibre offspring. Echocardiography measurements in independent male cohort showing c) heart rate and d) diastolic volume. All data is shown as mean±SEM. 2-way ANOVA with FDR adjustment for multiple comparisons. n=5-9.

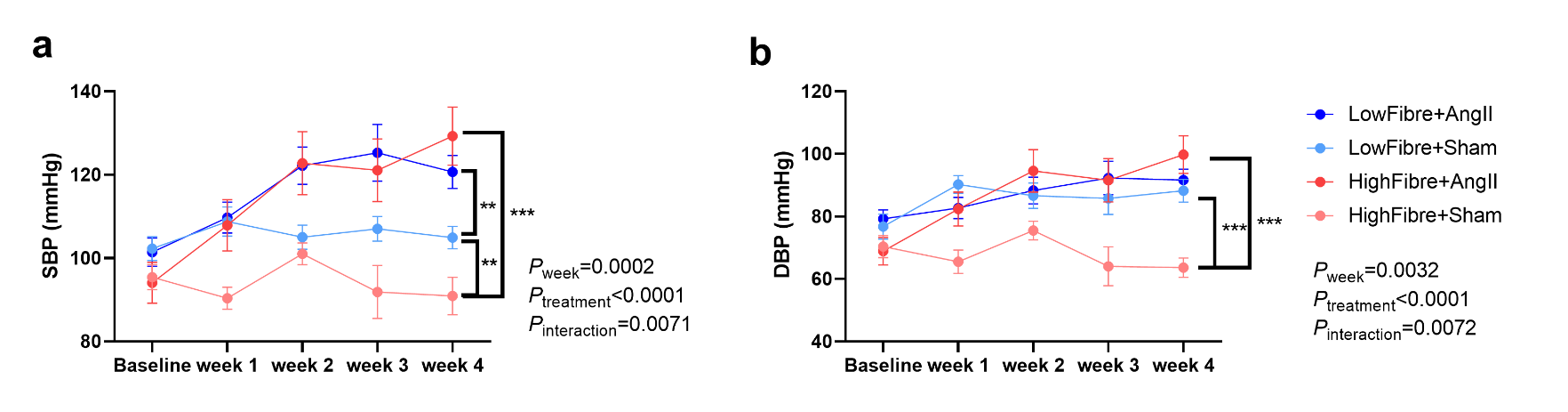

**Supplementary Figure 4.** Non-invasive tail-cuff blood pressure measurements in male high-fibre and low-fibre offspring showing a) systolic and b) diastolic blood pressure over 4 weeks. All data is shown as mean±SEM. 2-way ANOVA (repeated measures) with FDR adjustment for multiple comparisons. n=7-14/group. ***P*<0.01, ****P*<0.001.

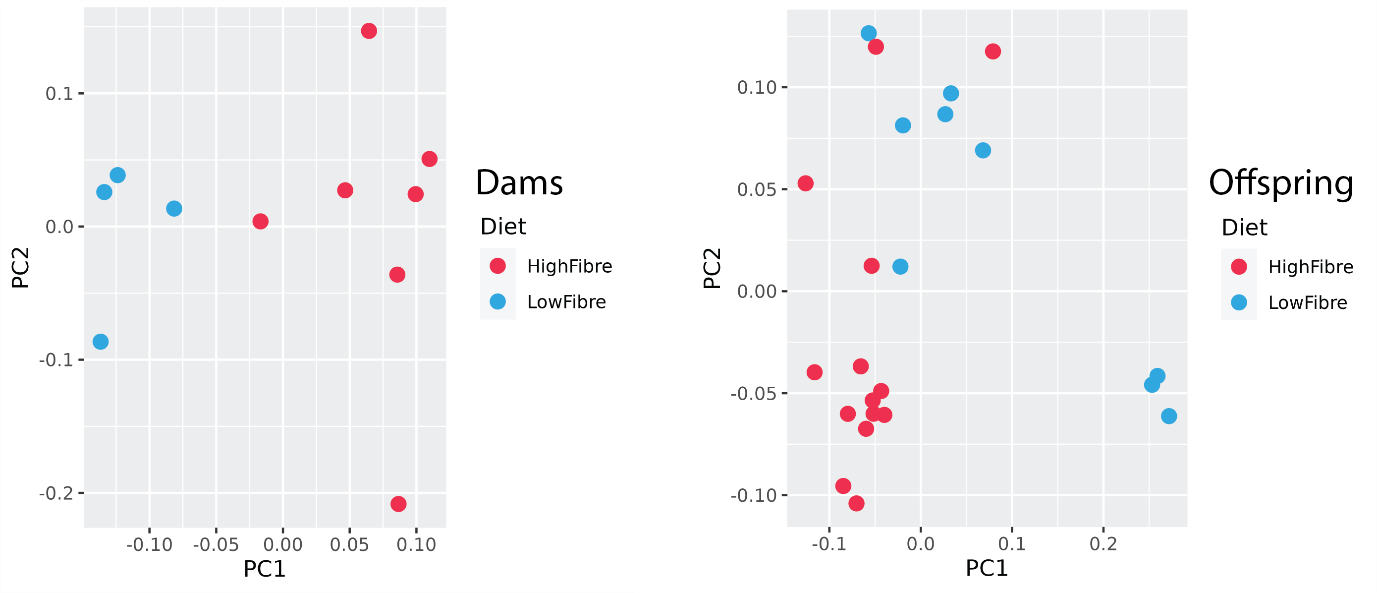

**Supplementary Figure 5.** Principal coordinate analysis plot showing differences in gut microbiome composition of dams (left) and offspring (right) of independent cohort. Dams n=4-7, offspring n= 9-14.

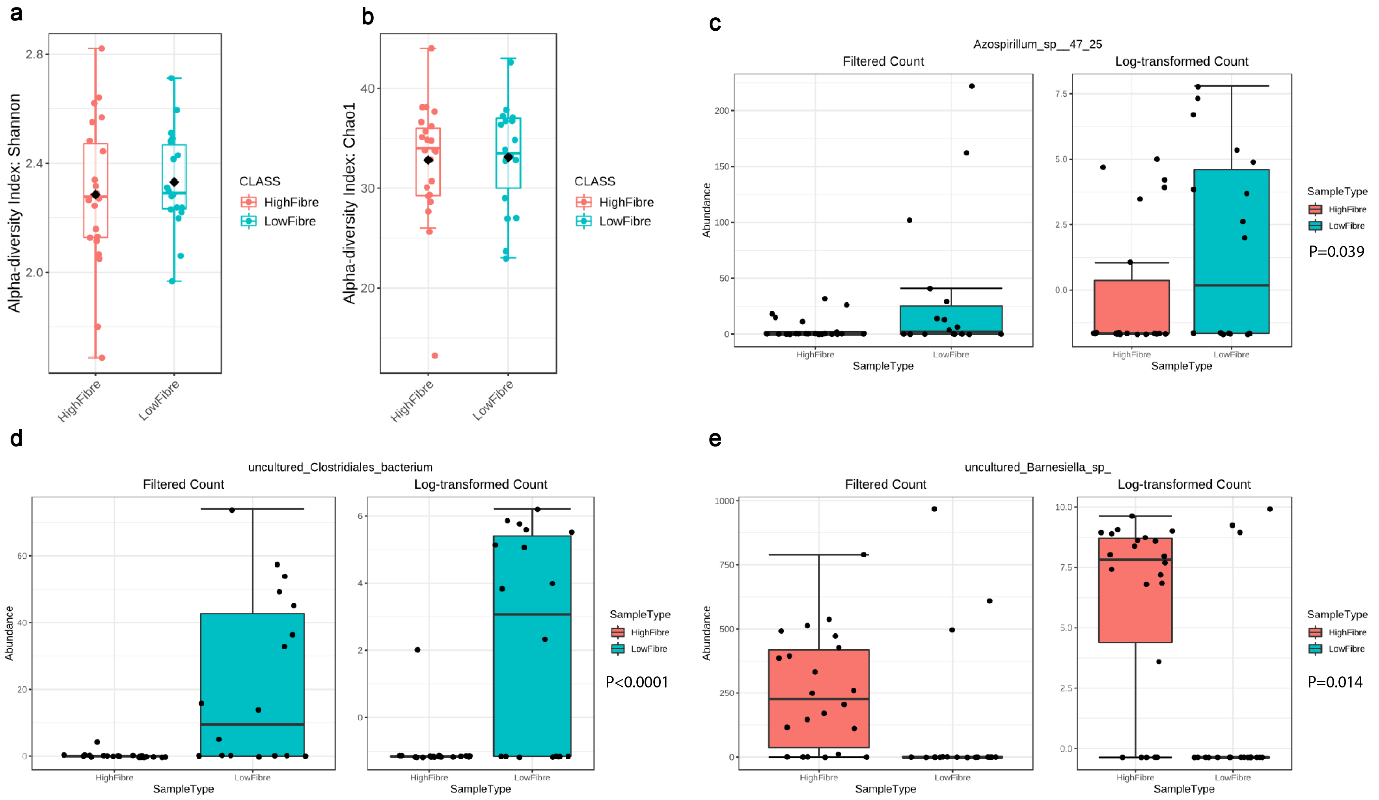

**Supplementary Figure 6.** 16S α-diversity metrics: a) Shannon index and b) Chao1 10WOA male offspring. Relative abundance of c) *Azospirillum*_sp__47_25, d) uncultured_*Clostridiales_bacterium*, and e) uncultured_*Barnesiella_*sp_ in caecal samples of male offspring. n=6-15/group

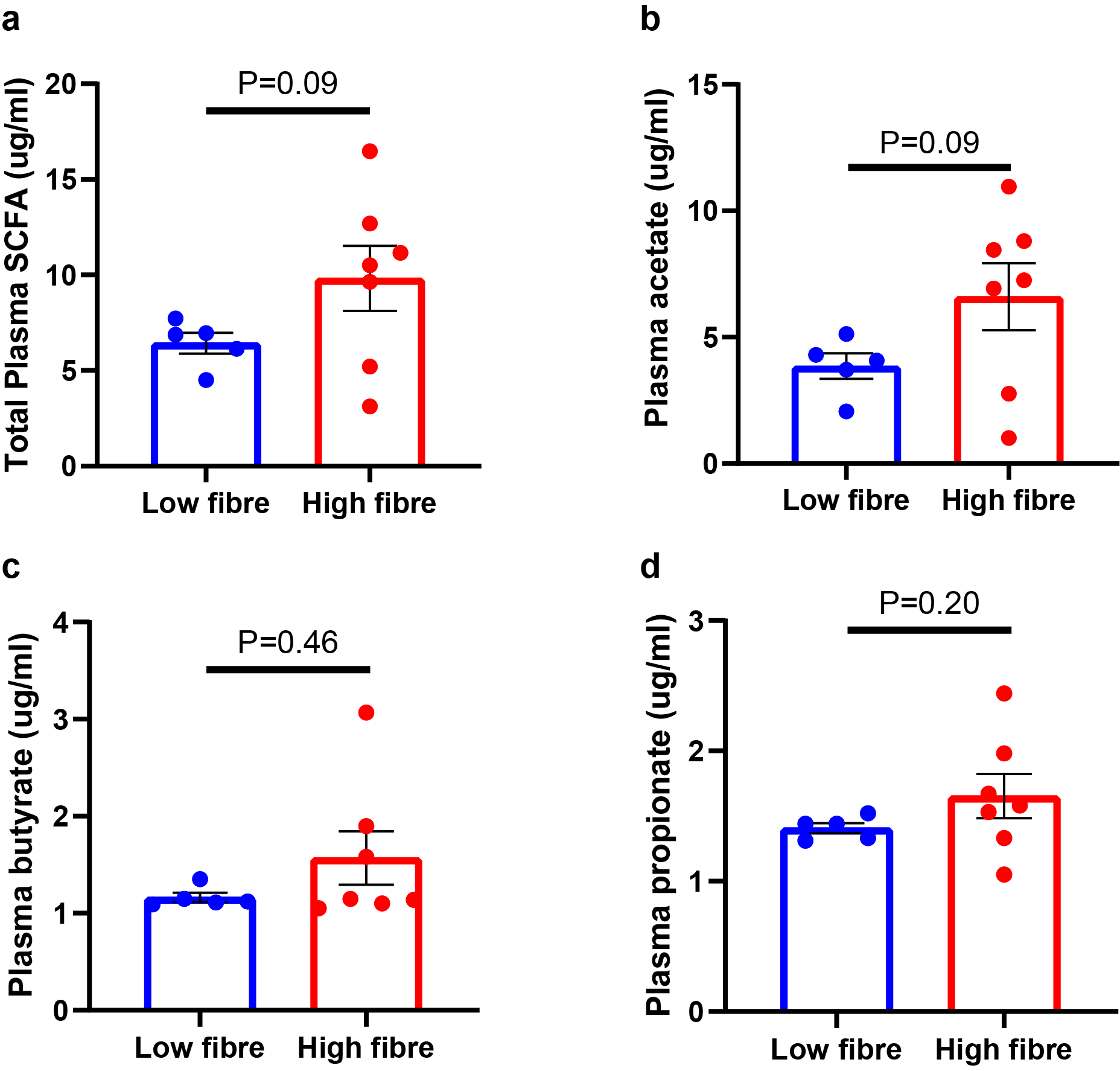
**Supplementary Figure 7.** Plasma short-chain fatty acid (SCFA) quantification of low fibre and high fibre fed mothers. a) Total plasma SCFA, b) plasma acetate, c) plasma butyrate, and d) Plasma propionate. All data is shown as mean±SEM. Dams n=5-7.

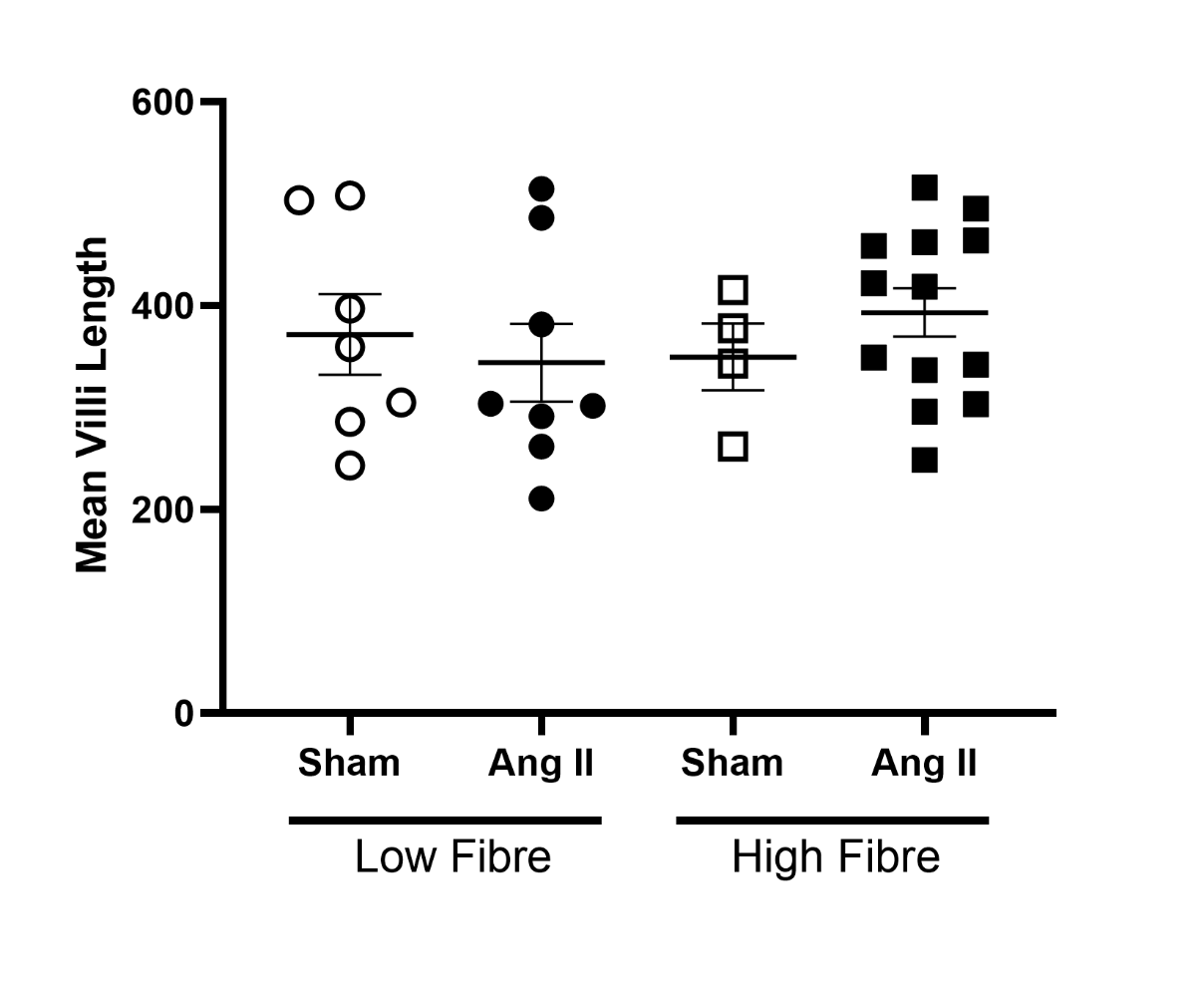
**Supplementary Figure 8.** Intestinal villi length in the offspring. All data is shown as mean±SEM. n=4-13

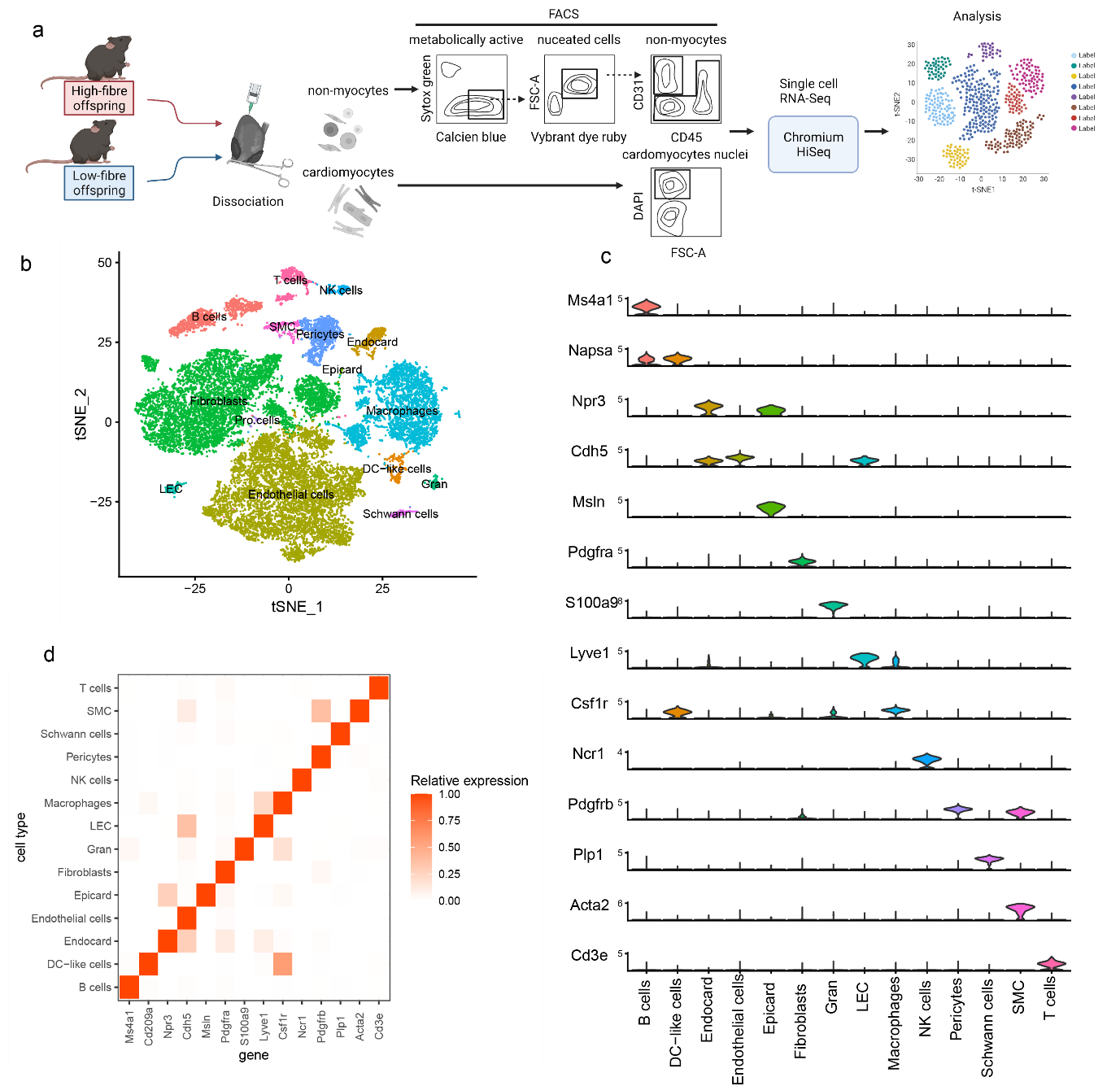
**Supplementary Figure 9.** Single-cell RNA sequencing analysis of cardiac tissue. a) Schematic outline of cell isolation protocol, b) t-distributed stochastic neighbour embedding visualisation of cell clusters, c) plots showing cell-specific gene markers used, d) heat map showing expression of each gene marker. n=8/diet (4 male and 4 female).

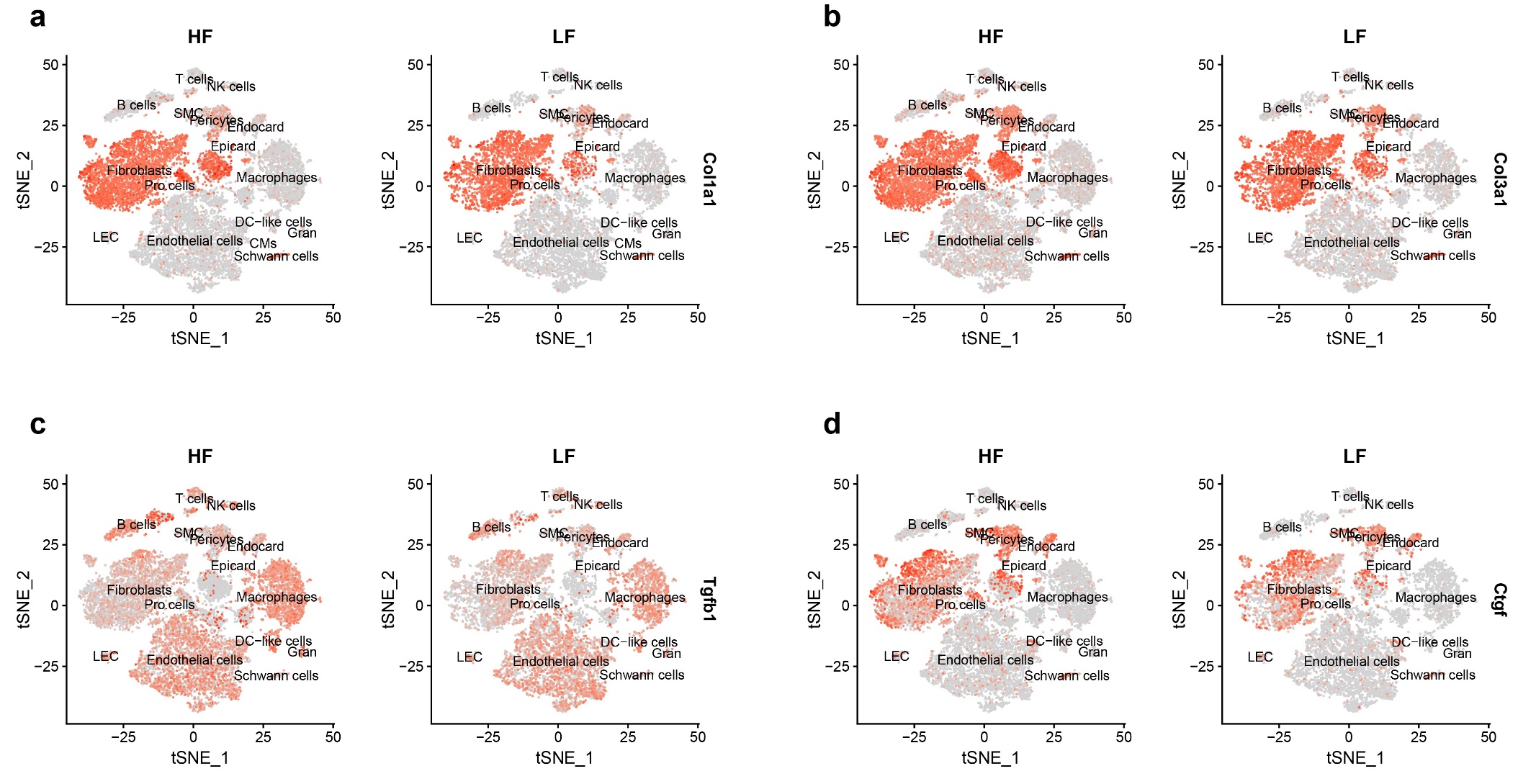
**Supplementary Figure 10.** Global expression of a) collagen 1a1, b) collagen 3a1, c) transforming growth factor beta, and d) connective tissue growth factor in the cardiac cellulome. n=8/diet (4 male and 4 female).

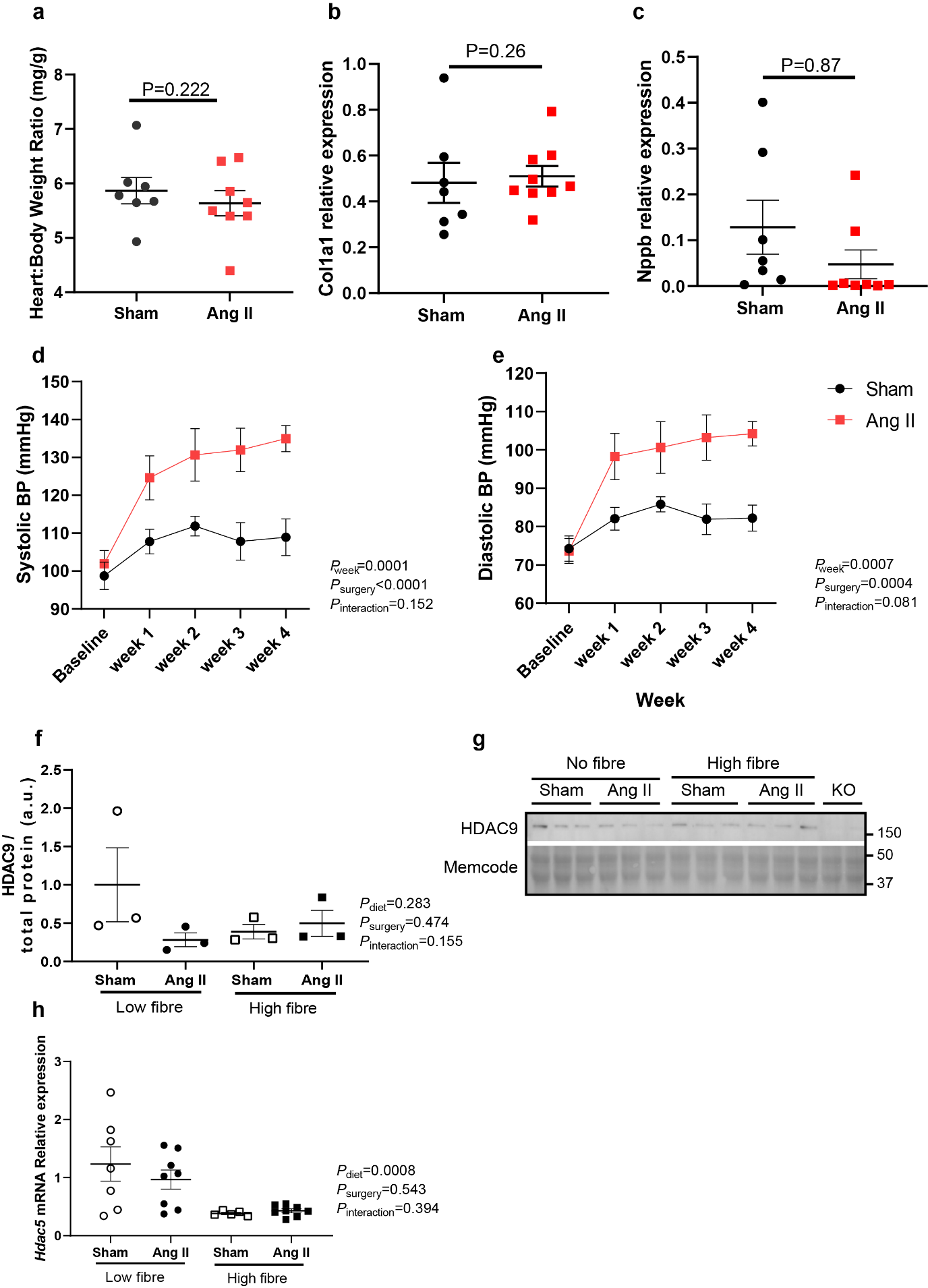
**Supplementary Figure 11. HDAC9 role in the offspring’s cardioprotection following maternal fibre intake.** a) heart to body weight ratio of HDAC9 KO male offspring from high-fibre fed mothers. Relative expression of b) collagen 1 a1 and c) natriuretic peptide b mRNA relative to *Gapdh*. d) Systolic and e) diastolic blood pressure of HDAC9KO male mice. f) HDAC9 protein quantification relative to total protein in wildtypes high-fibre and low-fibre male offspring and g) Western blot showing HDAC9 protein levels. h) HDAC5 mRNA gene expression in wildtype high-fibre and low-fibre male offspring. All data is shown as mean±SEM. n=3-8/group, 2-way ANOVA with false discovery rate adjustment for multiple comparisons.

**References**

1.	Kaye, D. M. *et al.* Deficiency of Prebiotic Fiber and Insufficient Signaling Through Gut Metabolite-Sensing Receptors Leads to Cardiovascular Disease. *Circulation* **141**, 1393-1403 (2020).

2.	Jama, H. A. *et al.* Manipulation of the gut microbiota by the use of prebiotic fibre does not override a genetic predisposition to heart failure. *Scientific Reports* **10**, 17919 (2020).

3.	Bolyen, E. *et al.* Reproducible, interactive, scalable and extensible microbiome data science using QIIME 2. *Nature Biotechnology* **37**, 852-857 (2019).

4.	Callahan, B. J. *et al.* DADA2: High-resolution sample inference from Illumina amplicon data. *Nat Methods* **13**, 581-583 (2016).

5.	Price, M. N., Dehal, P. S. & Arkin, A. P. FastTree 2--approximately maximum-likelihood trees for large alignments. *PLoS One* **5**, e9490 (2010).

6.	Katoh, K., Misawa, K., Kuma, K. & Miyata, T. MAFFT: a novel method for rapid multiple sequence alignment based on fast Fourier transform. *Nucleic Acids Res* **30**, 3059-3066 (2002).

7.	Chong, J., Liu, P., Zhou, G. & Xia, J. Using MicrobiomeAnalyst for comprehensive statistical, functional, and meta-analysis of microbiome data. *Nat Protoc* **15**, 799-821 (2020).

8.	Dhariwal, A. *et al.* MicrobiomeAnalyst: a web-based tool for comprehensive statistical, visual and meta-analysis of microbiome data. *Nucleic Acids Res* **45**, W180-W188 (2017).

9.	FastQC: a quality control tool for high throughput sequence data v. 0.11.9 (2010).

10.	Bolger, A. M., Lohse, M. & Usadel, B. Trimmomatic: a flexible trimmer for Illumina sequence data. *Bioinformatics* **30**, 2114-2120 (2014).

11.	Nurk, S., Meleshko, D., Korobeynikov, A. & Pevzner, P. A. metaSPAdes: a new versatile metagenomic assembler. *Genome Res* **27**, 824-834 (2017).

12.	Franzosa, E. A. *et al.* Species-level functional profiling of metagenomes and metatranscriptomes. *Nat Methods* **15**, 962-968 (2018).

13.	Langmead, B. & Salzberg, S. L. Fast gapped-read alignment with Bowtie 2. *Nat Methods* **9**, 357-359 (2012).

14.	Phipson, B., Lee, S., Majewski, I. J., Alexander, W. S. & Smyth, G. K. ROBUST HYPERPARAMETER ESTIMATION PROTECTS AGAINST HYPERVARIABLE GENES AND IMPROVES POWER TO DETECT DIFFERENTIAL EXPRESSION. *Ann Appl Stat* **10**, 946-963 (2016).

15.	R: A language and environment for statistical computing v. 4.0.3 (R Foundation for Statistical Computing, Vienna, Austria, 2010).

16.	Ritchie, M. E. *et al.* limma powers differential expression analyses for RNA-sequencing and microarray studies. *Nucleic acids research* **43**, e47-e47 (2015).

17.	Wickham, H. *et al.* Welcome to the Tidyverse. *Journal of Open Source Software* **4**, 1686 (2019).

18.	Fernandes, A. D., Macklaim, J. M., Linn, T. G., Reid, G. & Gloor, G. B. ANOVA-Like Differential Expression (ALDEx) Analysis for Mixed Population RNA-Seq. *PLOS ONE* **8**, e67019 (2013).

19.	Wickham, H. *ggplot2: Elegant Graphics for Data Analysis* 213 (Springer-Verlag, New York, United States of America, 2009).

20.	K N, H. *et al.* Sex-Based Mhrt Methylation Chromatinizes MeCP2 in the Heart. *iScience* **17**, 288-301 (2019).

21.	Khurana, I. *et al.* SAHA attenuates Takotsubo-like myocardial injury by targeting an epigenetic Ac/Dc axis. *Signal Transduction and Targeted Therapy* **6**, 159 (2021).
